# Mechanism of K63-linked polyubiquitin recognition and cleavage by the BRCA1-A complex

**DOI:** 10.64898/2026.06.05.730395

**Authors:** Martina Foglizzo, Arindam Datta, Oksana Degtjarik, Hirunika Perera, Jordan Liburd, Upasana M. Sykora, Sri Ranjani Ganji, Gemma Wildsmith, Francesca Chandler, Lisa J. Campbell, Antonio N. Calabrese, Roger A. Greenberg, Elton Zeqiraj

## Abstract

Deubiquitylases modulate cellular processes by cleaving monoubiquitin or polyubiquitin chains. The ARISC–RAP80 complex partners with BRCA1–BARD1 to form the BRCA1-A super-complex, which recognizes K63-linked ubiquitin chains at DNA damage sites. ARISC–RAP80 contains multiple ubiquitin-binding sites, yet how these influence recognition and cleavage of K63-polyubiquitylated substrates remains unknown. We discover that a composite three-subunit interface allows ARISC–RAP80 to position K63-linked polyubiquitin chains in its catalytic site. Substrate recognition is further supported by RAP80 and non-catalytic ubiquitin-binding sites that impose a compact conformation to K63-polyubiquitylated substrates. This mechanism exploits the inherent flexibility of long ubiquitin chains and differs considerably from other deubiquitylases. Structure-guided mutagenesis validate ubiquitin chain interactions, and cell-based assays demonstrate a functional role of the observed interfaces in chromatin recruitment. Our findings define mechanisms of polyubiquitin chain decoding and cleavage by ARISC–RAP80, linking ubiquitin reading and erasing functions to BRCA1-A mediated DNA damage responses.

## Introduction

The covalent attachment of ubiquitin to substrate proteins and to other ubiquitin molecules generates an ensemble of structurally and functionally diverse post-translational modifications, commonly referred to as the “ubiquitin code”^1,2^. Polyubiquitin chains linked via any of the seven lysine (K) residues in ubiquitin or its N-terminal methionine (M1) have been identified in human cells, with K48- and K63-linked chains being the most abundant^3,4^. While K48-linked chains have been primarily implicated in proteasomal degradation of the target substrates, non-degradative functions have been described for K63-linked ubiquitin moieties, including intracellular trafficking, immune signalling, and maintenance of genome stability^1,5–9^. Structural studies have highlighted that K48-linked chains assume compact conformations while K63-linked polymers form extended structures that are inherently more flexible in their conformations^10,11^. The topology adopted by ubiquitin polymers is thought to direct how ubiquitin chains of different linkages are “read out” and decoded by specific ubiquitin binding domains (UBDs), enabling signal transmission and downstream cellular processes^12–17^. However, relationships between K63-linked ubiquitin polymers recognition and cleavage beyond diubiquitin chains^18–22^ is still poorly understood.

Deubiquitylating enzymes (DUBs) cleave polyubiquitin chains or ubiquitin-substrate linkages, thereby controlling protein activity, localisation, stability, and function^1,23–25^. The human genome encodes for over 100 DUBs, which are classified as either cysteine proteases or zinc metalloproteases based on their catalytic mechanisms^23,26^. The Jab1/Mov34/Mpr1 Pad1 N-terminal (JAMM/MPN) metalloenzyme family consist of 14 known members in humans, and comprises both catalytically active (MPN^+^) DUBs and catalytically dead (MPN^-^) pseudo-DUBs^24^.

BRCC36 is a JAMM/MPN metalloenzyme that selectively cleaves K63-linked ubiquitin chains^27,28^ as part of two distinct macromolecular assemblies – the nuclear Abraxas1-regulated isopeptidase complex (ARISC) and the cytoplasmic BRCC36 isopeptidase complex (BRISC). ARISC interacts with the ubiquitin E3 ligase heterodimer composed of breast cancer type 1 susceptibility protein (BRCA1) and BRCA1-associated RING domain protein 1 (BARD1) to form the BRCA1-A supercomplex that regulates DNA damage repair. The cytoplasmic BRISC complex promotes interferon-dependent signaling by stabilising the type I interferon receptor (IFNAR1)^29–32^. In ARISC and BRISC assemblies, BRCC36 (MPN^+^) requires selective pairing with an inactive pseudo-DUB partner (MPN^-^), Abraxas1 and Abraxas2, for activity^28,33–35^. In addition, both complexes are characterised by accessory subunits, BRCC45 and MERIT40, forming 2:2:2:2 heterotetramers with characteristic U-shaped architectures^33–38^. The DUB activity of ARISC and BRISC is also regulated via selective interactions with protein partners, namely receptor-associated protein 80 (RAP80) and serine hydroxymethyltransferase 2 (SHMT2). RAP80 and SHMT2 target ARISC and BRISC complexes to their biological substrates in the nucleus and cytoplasm, respectively. RAP80 mediates the response of ARISC to DNA double-strand breaks (DSBs) while SHMT2 guides BRISC regulation of cytokine signalling^29–32^. However, the molecular details that govern ARISC and BRISC interaction with ubiquitin and specificity for hydrolysing K63-linked polyubiquitin chains are undetermined.

Here, we describe molecular mechanisms underpinning ARISC and ARISC–RAP80 recognition and cleavage of K63-linked ubiquitin chains. Biochemical experiments demonstrate that RAP80 confers enhanced activity to ARISC, specifically in the presence of long polyubiquitin substrates. Cryogenic-electron microscopy (cryo-EM) structures of ARISC and ARISC–RAP80 in complex with K63-linked ubiquitin chains reveal a conserved substrate binding mode, consisting of multi-subunit interactions within the ARISC catalytic core. Substrate recognition is aided by multiple domains in RAP80, and by non-catalytic ubiquitin sites poised to bind mono or polyubiquitylated chromatin substrates. Structure-guided mutagenesis and cross linking-mass spectrometry confirm the mode of ubiquitin chain interaction with ARISC, and cell-based studies validate the importance of ubiquitin-binding surfaces in ARISC–RAP80 recruitment to DSBs. Taken together, our findings establish that coordination between ubiquitin-binding moieties on different subunits promotes ARISC and ARISC–RAP80 dependent decoding of K63-linked polyubiquitin chains, with direct implications for understanding BRCA1-A complex recruitment to chromatin and its functions in DNA repair.

## Results

### RAP80 stimulates ARISC activity on long polyubiquitin chains

RAP80 contains tandem ubiquitin-interacting motifs (UIMs) at its N-terminus (**Fig. 1a**), which recognise lysine 63 (K63)-linked ubiquitin chains^29–31,39^. We hypothesised that interactions between the RAP80 UIMs and polyubiquitin (polyUb) chains favour the subsequent engagement of polyUb with the BRCC36 active site, promoting ubiquitin chain hydrolysis by ARISC. To assess this, we purified stoichiometric amounts of full-length (FL) ARISC and ARISC–RAP80 complexes from insect cells (**Fig. 1a,b**), and measured their activities against a panel of K63-linked di and polyUb substrates (Ub2-Ub9) generated in-house (**Supplementary Fig. 1a**, *left panel*). The difference in ARISC and ARISC–RAP80 deubiquitylase (DUB) activity was modest when using di and triUb (Ub2 and Ub3) chains (**Fig. 1c** and **Supplementary Fig. 1b,c**). By contrast, the activities of ARISC and ARISC–RAP80 increased in the presence of tetraUb (Ub4) chains as previously reported^37^, with a clear enhancement in substrate cleavage by ARISC–RAP80 over ARISC (**Fig. 1c** and **Supplementary Fig. 1b,c**). Curiously, we observed a reduction of ARISC and ARISC–RAP80 DUB activities with pentaUb (Ub5) chains, while both complexes showed a progressive increase in ubiquitin chain hydrolysis when tested against hexa up to nonaUb (Ub6-Ub9) (**Fig. 1c** and **Supplementary Fig. 1b,c**). Of note, the activity of ARISC–RAP80 was visibly higher than ARISC in the presence of Ub6 and longer chains (**Fig. 1c** and **Supplementary Fig. 1b,c**), supporting our initial hypothesis. Taken together, these results demonstrate that: 1) ARISC and ARISC-RAP80 cleave ubiquitin chains equal to, or longer than, Ub6 more efficiently than shorter chains; and that 2) FL RAP80 confers enhanced activity to ARISC, particularly in the presence of Ub6 and longer chains.

**Fig. 1:**
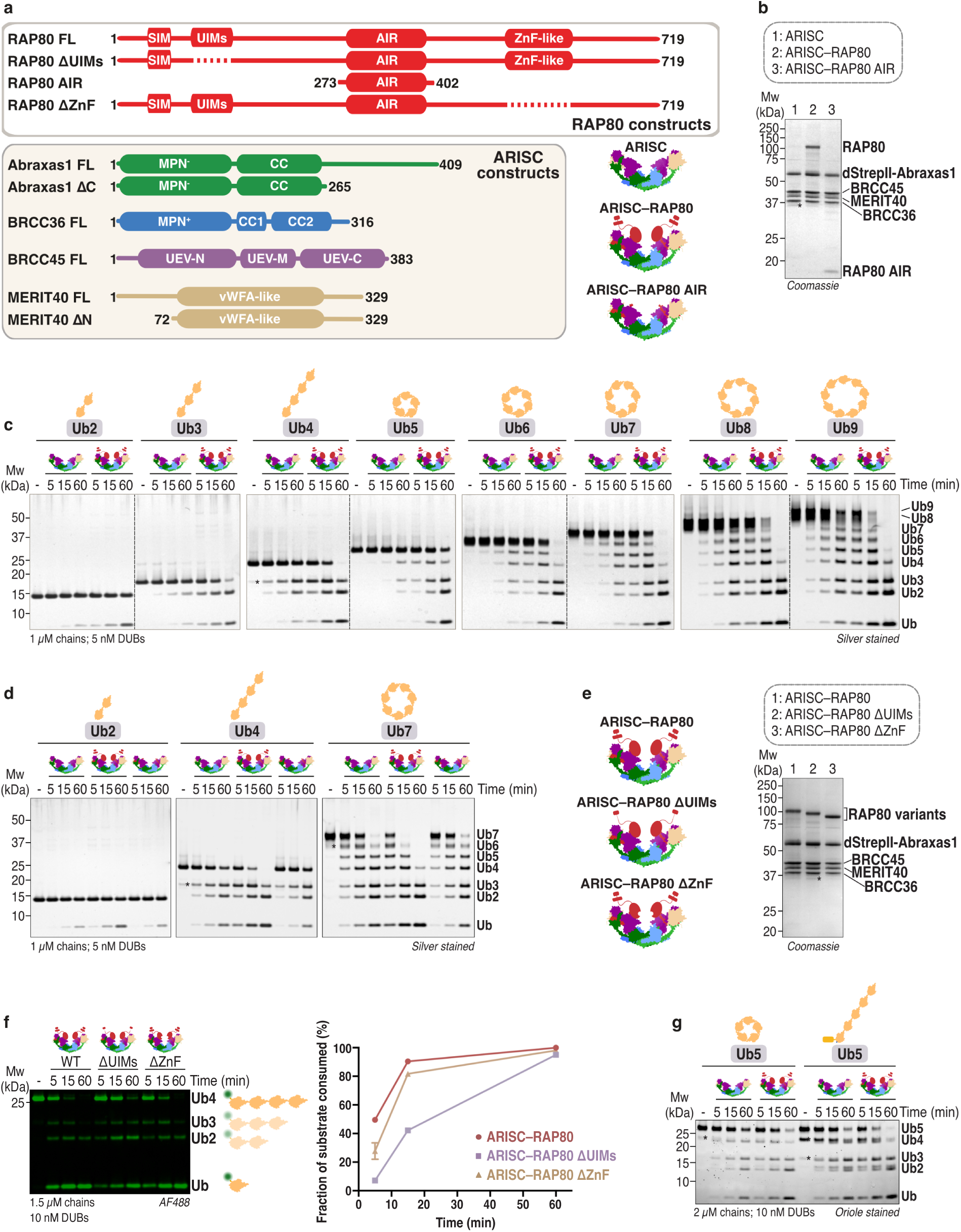
ARISC deubiquitylase activity is regulated by FL RAP80. **a**, Domain architecture of RAP80 and ARISC constructs. FL, full-length; SIM, small ubiquitin-like modifier (SUMO)-interacting motif; UIM, ubiquitin-interacting motif; AIR, Abraxas1-interacting region; ZnF, zinc finger; MPN, Mpr1, Pad1 N-terminal; CC, coiled coil; UEV, ubiquitin E2 variant; vWFA, von Willebrand factor type A (*left*). Schematics of indicated complexes (*right*). **b**, SDS-PAGE analysis of ARISC, ARISC–RAP80, and ARISC–RAP80 AIR. **c**, K63-linked ubiquitin chains (1 µM) were incubated with ARISC or ARISC–RAP80 (5 nM) for the indicated time points. Cleavage activity was analysed by SDS-PAGE and silver staining. Data are representative of two independent experiments. **d**, K63-Ub2, -Ub4, and - Ub7 chains (1 µM) were incubated with ARISC, ARISC–RAP80, or ARISC–RAP80 AIR (5 nM) for the indicated time points. Cleavage activity was analysed as in **c**. Data are representative of three independent experiments. **e**, Schematics (*left*) and SDS-PAGE analysis (*right*) of indicated complexes. dStrepII, double StrepII tag. * indicates Abraxas1 degradation product. **f**, Alexa-Fluor 488 (AF488) labelled distally (AF488-^Cys^Ub4^K63R^) blocked K63-Ub4 chains (1.5 µM) were incubated with ARISC–RAP80, ARISC–RAP80 ΔUIMs, or ARISC–RAP80 ΔZnF (10 nM) for the indicated time points. Cleavage activity was analysed by SDS-PAGE and fluorescence scanning (*left*; see **Methods**). The disappearance of the K63-Ub4 parent band was quantified using densitometry, and plotted as fraction of substrate consumed (%). Data points are mean ± SEM of two independent experiments (*right*). **g**, Cyclical and linear K63-Ub5 chains (2 µM) were incubated with ARISC or ARISC–RAP80 (10 nM) for the indicated time points. Cleavage activity was analysed by SDS-PAGE and Oriole staining. Data are representative of two independent experiments. Ub, ubiquitin; DUB, deubiquitylating enzyme. * indicates lower molecular weight ubiquitin species.

The RAP80 Abraxas1-interacting region (AIR) represents the minimal domain that stably binds to ARISC^37^. Previous studies have shown that RAP80 AIR engages with the arm regions in ARISC, making extensive contacts with the C-terminal ubiquitin E2 variant (UEV-C) domains of BRCC45, MERIT40, and the Abraxas1 C-termini^37,40^. To determine if the minimal RAP80 AIR fragment confers increased activity to ARISC, we assayed a RAP80 variant (ARISC–RAP80 AIR) (**Fig. 1a,b**), alongside ARISC and ARISC–RAP80, against purified K63-Ub2, -Ub4, and -Ub7 chains (**Supplementary Fig. 1a**, *left panel*). The activity of ARISC–RAP80 AIR was more similar to ARISC in the presence of all chain lengths tested, and was considerably reduced when compared to ARISC–RAP80 (**Fig. 1d**). To better assess the catalytic efficiencies and substrate binding abilities of our DUB complexes, we introduced a fluorescent label at the distal end of a K63-Ub4 chain^37^ (**Supplementary Fig. 2a,c**) and measured the activities of ARISC and ARISC–RAP80 variants against non-saturating and saturating substrate concentrations. In agreement with our initial analyses, the activity of ARISC–RAP80 was visibly increased when compared to ARISC and ARISC–RAP80 AIR in the presence of non-saturating and saturating substrate (**Fig. 1d** and **Supplementary Fig. 1d,e**). Similarly, ARISC and ARISC–RAP80 AIR displayed comparable enzymatic activities at non-saturating and saturating concentrations of K63-Ub4 (**Fig. 1d** and **Supplementary Fig. 1d,e**). These combined data indicate that: 1) ARISC–RAP80 possess higher catalytic efficiency and ability to bind ubiquitin chains than ARISC and ARISC–RAP80 AIR; and that 2) in isolation, the minimal RAP80 AIR fragment does not enhance ARISC activity nor contribute to ARISC-mediated ubiquitin chain binding. Our results therefore suggest that other ubiquitin binding regions or domains within RAP80 are responsible for modulating ARISC enzymatic function.

To identify the region in RAP80 that supports ARISC interaction with ubiquitin chains and contributes to sustained DUB activity, we purified stoichiometric amounts of two ARISC–RAP80 variants in which the RAP80 UIM and zinc finger (ZnF)-like domains were deleted (here indicated as ARISC–RAP80 ΔUIMs and ARISC–RAP80 ΔZnF) (**Fig. 1a,e**). We subsequently assayed these protein complexes, alongside ARISC–RAP80, against distally-labelled K63-Ub4 chain. The activity of ARISC–RAP80 ΔUIMs was considerably decreased when compared to ARISC–RAP80, while we observed only a marginal reduction in the activity of ARISC–RAP80 ΔZnF under the same experimental conditions (**Fig. 1f**). These results indicate that the UIM domains represent the major ubiquitin-interacting site in RAP80. Our data also suggest that the ZnF region may provide a secondary surface for ubiquitin chain coordination, albeit with reduced efficiency compared to the UIM domains.

Lastly, we compared the enzymatic activities of ARISC and ARISC–RAP80 with the cytoplasmic BRISC complex^37,38,41^. We tested all recombinantly purified complexes against K63-Ub2, -Ub4, and -Ub7 chains (**Supplementary Fig. 1a**, *left panel*, **f**), and detected increased activity of BRISC over ARISC and ARISC-RAP80 when K63-Ub2 was used as substrate (**Supplementary Fig. 1g**). BRISC activity remained higher than ARISC in the presence of K63-Ub4 and -Ub7 chains, but only moderately enhanced compared to ARISC–RAP80 (**Supplementary Fig. 1g**). These results suggest a specific role for RAP80 in the recognition of polyubiquitin chains, and highlight differences in the regulatory mechanisms underpinning ARISC and BRISC enzymatic activities.

### Lysine 63-polyubiquitin form linear and cyclical chains

Intrigued by the unexpected decrease of ARISC and ARISC–RAP80 catalytic activities towards K63-Ub5 (**Fig. 1c** and **Supplementary Fig. 1b**), we characterised our purified ubiquitin chains using intact mass spectrometry analyses. Results indicated a strong agreement between measured and calculated masses for K63-linked Ub2, Ub3, and Ub4 chains (**Supplementary Fig. 1a**, *left panel*, **Supplementary Fig. 2d**, and **Supplementary Fig. 3a-c**), confirming the quality of our reagents. Surprisingly, we detected a mass difference of approximately 18 Da, equivalent to the loss of one water molecule, when comparing measured and calculated molecular weights for Ub5-Ub9 chains (**Supplementary Fig. 1a**, *left panel*, **Supplementary Fig. 2d**, and **Supplementary Fig. 3d-h**). These analyses suggest that our purified K63-Ub5 and longer chains assemble in a cyclical conformation, whereby the C-terminal G76 of the distal ubiquitin forms an isopeptide bond with K63 of the proximal ubiquitin moiety.

To further assess the nature of ubiquitin chain cyclisation, we purified AMSH* (STAM2–AMSH fusion), a related JAMM/MPN DUB that, like BRCC36, selectively cleaves K63-linked polyubiquitin chains^15,18,42^, and OTULIN, a member of the Ovarian Tumor (OTU) family with exclusive specificity for M1-linked chains^43,44^ (**Supplementary Fig. 4a**). We then assayed the resulting proteins, alongside ARISC and ARISC–RAP80, against cyclical K63-Ub5 and linear M1-Ub4 chains (**Supplementary Fig. 1a**, *left panel*, and **Supplementary Fig. 4a**). We detected robust activity for ARISC, ARISC–RAP80, and AMSH* in the presence of our cyclical K63-Ub5 substrate, with full conversion of Ub5 to monoubiquitin. Likewise, we observed high activity for OTULIN against M1-Ub4 chains (**Supplementary Fig. 4b-d**). Importantly, ARISC, ARISC–RAP80, and AMSH* did not display any activity towards M1-Ub4 and, accordingly, OTULIN did not hydrolyse our cyclical K63-Ub5 substrate (**Supplementary Fig. 4b-d**). These experiments strongly indicate that cyclisation of our ubiquitin chains does not involve M1 linkages, confirming our earlier hypothesis.

To circumvent this ubiquitin chain cyclisation *in vitro*, we generated a ubiquitin construct blocked at the N-terminus via an 8xHis tag and a lysine to arginine mutation at position 63 (K63R)^45^. We then used the resulting 8xHis-ubiquitin K63R variant and wild type (WT) ubiquitin to assemble distally blocked (here also referred to as “linear”) K63-linked ubiquitin chains of different lengths (**Supplementary Fig. 1a**, *right panel*). Intact mass spectrometry analyses detected a nearly perfect agreement between measured and calculated masses for all ubiquitin chains (**Supplementary Fig. 1a**, *right panel*, **Supplementary Fig. 2d**, and **Supplementary Fig. 3i-n**), confirming linear K63-linked ubiquitin chain assembly. We then assayed our DUB complexes against cyclical and linear K63-linked Ub5 chains (**Supplementary Fig. 1a**), and observed a moderately higher activity of ARISC and ARISC–RAP80 towards linear K63-Ub5 compared to cyclical substrate (**Fig. 1g**).

Collectively, these analyses indicate that ARISC and ARISC–RAP80 can interact with, and hydrolyse, cyclical and linear ubiquitin chains. These data also suggest that our DUB complexes employ different modes of interactions with their ubiquitin substrates, with cyclical chains likely adopting a more constrained conformation compared to their linear counterpart.

### Cryo-EM structure of the ARISC(E33A)–RAP80:K63-Ub7 complex

To determine the precise mechanisms underpinning ARISC–RAP80 interaction with ubiquitin chains, we performed single-particle cryogenic-electron microscopy (cryo-EM) analyses of our purified complexes in the presence of ubiquitin chains. We took advantage of an ARISC–RAP80 variant where the BRCC36 active site glutamate was mutated to alanine (E33A), rendering it inactive (**Supplementary Fig. 5a**)^35,40^. We set up cryo-EM grids with ARISC(E33A)–RAP80 and cyclical K63-Ub7, and used mild glutaraldehyde cross-linking to stabilise ARISC–RAP80 interactions (**Supplementary Fig. 5b**, *left panel*)^46^. This approach yielded high-quality data and allowed us to obtain a structure with the characteristic U-shaped architecture, previously seen for ARISC and BRISC assemblies^36–38^, as well as additional globular densities that could be attributed to ubiquitin (**Supplementary Fig. 6a,b**). Further three-dimensional (3D) refinement resulted in a consensus cryo-EM map at an overall resolution of 2.92 Å, although the local resolution around the BRCC45-MERIT40 arm regions and bound ubiquitin molecules was poor (**Supplementary Fig. 6b,c** and **Supplementary Table 1**). The particle orientation distribution was largely homogeneous across the map, with some preferential orientation mainly corresponding to top views of the complex (**Supplementary Fig. 7a**). Subsequent focused masking and local refinements improved the regional quality of the map, resulting in local resolutions ranging from 3.16 Å to 4.09 Å (**Supplementary Fig. 6b,d-j**, **Supplementary Fig. 7b-g**, and **Supplementary Table 1**).

We observe the highest resolution in the regions comprising the BRCC36-Abraxas1 super dimer and the N-terminal UEV (UEV-N) domain of both BRCC45 subunits (**Fig. 2a** and **Supplementary Fig. 6b,c**). The resolution is slightly decreased around the middle and last BRCC45 UEV domains (UEV-M and UEV-C), and becomes lower for the Abraxas1 C-terminal domains, MERIT40, and RAP80 AIR (**Fig. 2a** and **Supplementary Fig. 6b,c**). This is particularly evident for the left arm, and in agreement with the flexible nature of the ARISC and BRISC arm regions observed in previous structures^37,38,40,41^. Local refinements using masks comprising these regions improved the corresponding densities on both ARISC arms (**Supplementary Fig. 6b,d,e**). We therefore model built the entire BRCC45 molecules, and rigid-body fitted the Abraxas1 C-terminal domains and RAP80 AIR (residues 278-331 and 272-329, respectively), and MERIT40 using a previous model of the human ARISC–RAP80 AIR complex (**Fig. 2a**)^40^. Although clear density for the linker region connecting the Abraxas1 MPN^-^ to its C-termini was absent in the refined map (**Fig. 2a** and **Supplementary Fig. 6b**), this was clearly traceable in the lower-resolution ab-initio map (**Supplementary Fig. 5c**). Interestingly, apart from RAP80 AIR, we could not visualise any additional RAP80 domains, which is consistent with the highly dynamic nature of most RAP80 regions and conformational heterogeneity we observed previously (**Fig. 1a**, **Fig. 2a**, **Supplementary Fig. 5c**, and **Supplementary Fig. 6b**)^40^.

**Fig. 2:**
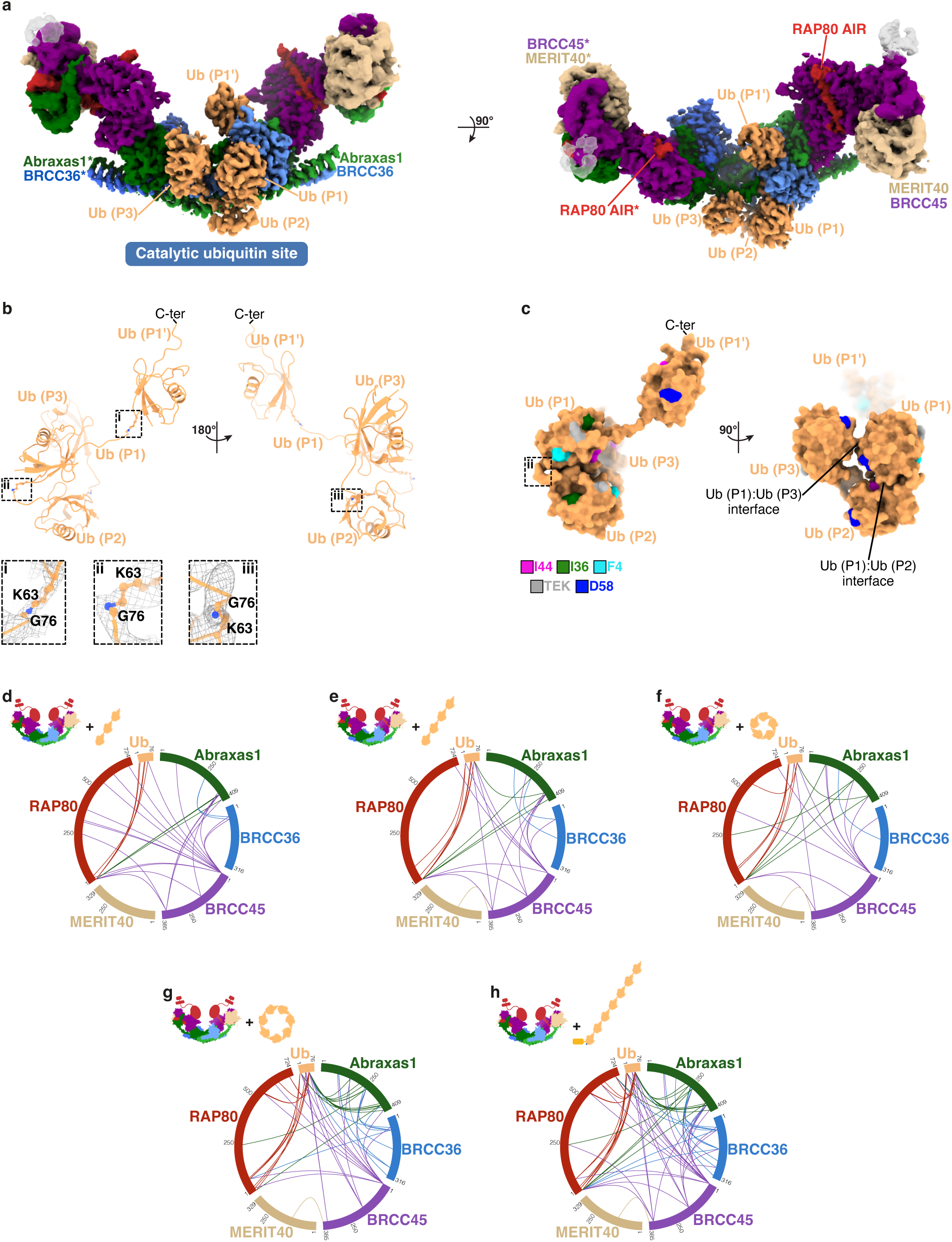
Cryo-EM structure of the ARISC(E33A)–RAP80:K63-Ub7 complex. **a**, Cryo-EM composite map of ARISC(E33A)–RAP80:K63-Ub7 at contour level of 0.0432. ARISC–RAP80 subunits and ubiquitin molecules are colored as in Fig. 1a and light orange; transparent densities correspond to the non-catalytic ubiquitin sites (see **Fig. 7a,b** and **Supplementary Fig. 14a,b**). AIR, Abraxas1-interacting region. **b,** Cartoon model of the K63-linked ubiquitin chains visualised in the ARISC(E33A)–RAP80:K63-Ub7 structure, with ubiquitin moieties colored as in **a**. Dashed black rectangles highlight the isopeptide bonds between each ubiquitin molecule (*top*). Close-up views and structural details of the isopeptide bonds between Ub (P1’):Ub (P1) (i), Ub (P1):Ub (P2) (ii), and Ub (P2):Ub (P3) (iii). Residues are shown as ball & sticks and the composite cryo-EM map as mesh (contoured at 0.0432) (*bottom*). **c,** Surface representation of the K63-linked ubiquitin chains depicted in **b**, with ubiquitin moieties colored as in **a**. The F4, I36, I44, and D58 patches as well as the TEK box are mapped onto each ubiquitin moiety and colored as indicated. The dashed black rectangle highlights the Ub (P1):Ub (P2) isopeptide bond (see panel **b**). C-ter, C-terminus. **d-h,** Chemical cross-linking and mass spectrometry analyses of ARISC(E33A)–RAP80 in complex with wild type K63-Ub3 (**d**), -Ub4 (**e**), -Ub5 (**f**), and -Ub7 (**g**) chains as well as with distally blocked K63-Ub7 (**h**). Proteins are labelled and their corresponding inter-subunit cross-links color-coded accordingly. Communal cross-links obtained from two independent experiments are shown. Crosslink sites involving ubiquitin were searched and mapped against a single ubiquitin sequence. Ub, ubiquitin.

Next, we focused on identifying the position of the K63-Ub7 chain in our ARISC–RAP80 consensus map. We immediately observed two well-defined globular regions on either side of one BRCC36 molecule, spanning its active site, which we attributed to ubiquitin molecules (**Fig. 2a** and **Supplementary Fig. 6b**). We defined the different ubiquitin moieties in the Ub7 chain based on their orientations relative to the BRCC36 active site, with the distal (P) ubiquitin providing the C-terminal carboxyl group (G76) and the proximal (P’) ubiquitin providing the lysine residue (K63) required for isopeptide bond formation^47,48^. We rigid-body fitted ubiquitin into the density corresponding to the distal site [referred to as Ub (P1)] (**Fig. 2a**), and performed local refinement on the proximal ubiquitin region [referred to as Ub (P1’)] to improve density connectivity (**Supplementary Fig. 6b,f**). This allowed us to rigid-body fit a second ubiquitin into this site and orient it so that K63 of Ub (P1’) forms an isopeptide bond with G76 of Ub (P1) (**Fig. 2a,b**). Further analyses of our consensus and ab-initio cryo-EM maps revealed additional densities extending towards the Abraxas1 subunit of the adjacent ARISC protomer (ARISC*, Abraxas1*). We could confidently place two additional ubiquitin molecules extending from Ub (P1) towards Abraxas1* [referred to as Ub (P2) and Ub (P3), respectively] (**Fig. 2a**, **Supplementary Fig. 5c**, and **Supplementary Fig. 6b,c,g**). The quality of the map in these regions was sufficient to visualise the isopeptide bonds between the ubiquitin moieties and connect them via K63 linkages (**Fig. 2a,b** and **Supplementary Fig. 6b,g**). Further visualisation of the ab-initio map revealed a globular density extending from Ub (P3) towards the BRCC45* arm (**Supplementary Fig. 5c**), which we attributed to a fifth ubiquitin molecule [Ub (P4)]. However, local refinement did not result in density improvement, likely due to the absence of stable contacts between Ub (P4) and ARISC. We therefore rigid-body fitted Ub (P4) in the low-resolution ab-initio map and oriented it so its C-terminal G76 residue is in proximity to K63 of Ub (P3) (**Supplementary Fig. 5c**). Although the precise position or orientation of Ub (P4) could not be determined due to poor map density and unresolved regions within both BRCC36 and RAP80, the cyclic conformation of K63-Ub7 suggests that this ubiquitin, together with the last two molecules, likely extend from Ub (P3) toward Ub (P1’) (**Supplementary Fig. 5d**).

Notably, ubiquitin molecules are bound to only one BRCC36 protomer in our structure while the corresponding BRCC36* interface appears unbound (**Fig. 2a**). Overlaying of the ubiquitin chains onto the BRCC36* active site showed only a small clash between the alpha 1 (α1) C-terminal regions of the two Ub (P1’) molecules (**Supplementary Fig. 5e**). Although we observe no clear ubiquitin density in both BRCC36 active sites simultaneously, considering this is a modest overlap, it is possible that structural re-arrangements of the ubiquitin moieties within ARISC may allow both BRCC36 protomers to concomitantly engage with their ubiquitin substrates.

### Tightly packed ubiquitins form a unique chain architecture

Next, we analysed the arrangement adopted by the ubiquitin monomers present in our ARISC–RAP80 ubiquitin-bound structure. As expected, P1’ and P1 ubiquitin display a long and extended conformation across the BRCC36 active site that is amenable to substrate engagement and cleavage (**Fig. 2a-c**). In addition, Ub (P1), Ub (P2), and Ub (P3) arrange in a tightly packed conformation that uses different surfaces in each of the ubiquitin molecules. The F4 patch in Ub (P1) together with loop4, the loop5 region surrounding the K63-linked isopeptide bond, and β4 at the C-terminus make extensive contacts with β1, β3, β4, and the C-terminal tail in Ub (P2) (**Fig. 2b,c**). Notably, Ub (P1) loop4 is positioned directly in front of the Ub (P2) I44 hydrophobic patch^11^ (**Fig. 2b,c**). Interactions between Ub (P1) and Ub (P3) are instead largely mediated by the region preceding the negatively charged D58 cluster^11^ of both ubiquitin molecules (**Fig. 2b,c**). These analyses reveal a highly interconnected ubiquitin architecture not previously reported for K63-linked chains in the absence or presence of binding partners^10,11,18–21,49,50^. By exploiting the inherent flexibility of K63-linked chains, this architecture exposes distinct ubiquitin surfaces along a single chain. Different ubiquitin-binding domains could therefore engage different interfaces on the same chain, allowing one chain type to potentially drive diverse signalling outcomes.

### RAP80 promotes selective recognition of polyubiquitin chains

To independently validate our structural model and understand how ARISC–RAP80 interacts with ubiquitin chains of different lengths and types, we performed chemical cross-linking mass spectrometry analysis (XL-MS) of ARISC(E33A)–RAP80 in the absence and presence of WT and distally blocked K63-linked Ub3, Ub4, Ub5, and Ub7 chains (**Fig. 2d-h**, **Supplementary Fig. 1a**, and **Supplementary Fig. 5f**). We used the XL-MS-cleavable cross-linker disuccinimidyl dibutyric urea (DSBU), which has a spacer length of 12.5 Å and a reachable Cα-Cα distance between residues of ∼ 32 Å^51^, and initially identified intermolecular cross-links between all ARISC–RAP80 subunits (**Fig. 2d-h** and **Supplementary Fig. 5f**). These data are consistent with our previous work^40^, and validate the structural arrangement adopted by the heterodecameric ARISC–RAP80 complex. In addition, we detected several cross-links between ubiquitin and the Abraxas1, BRCC45, and RAP80 subunits (**Fig. 2d-h**). These observations are consistent with our structural model (**Fig. 2a** and **Supplementary Fig. 5c**) and indicate that the BRCC45 arm regions interact with both linear and cyclical chains, likely via its multiple UEV domains. Mapping of the cross-link sites in our cryo-EM structure revealed direct interactions between BRCC45 K162 and ubiquitin K6, K33, and K48, supporting the idea that Ub (P4) is positioned near BRCC45* (**Supplementary Fig. 5c,g**). Strikingly, we detected a higher number of cross-links between RAP80 and ubiquitin as a function of increased ubiquitin chain length (**Fig. 2d-g**), and comparable cross-link numbers and/or sites between Abraxas1, BRCC45, or RAP80 and cyclical and linear K63-Ub7 substrates (**Fig. 2g,h**). This suggests that ARISC–RAP80 adopts a conserved mode of interaction with both ubiquitin chain types. Our analyses also indicate that ubiquitin molecules within a K63-linked chain, whether in its cyclical or linear form, localise in proximity to known RAP80 ubiquitin-interacting motifs we previously reported^40^, such as its N-terminus and UIM domains, but also to regions not previously associated with ubiquitin binding, including the linker region between the RAP80 small ubiquitin-like modifier (SUMO)-interacting motif (SIM) and UIMs, RAP80 AIR, and the C-terminal ZnF domain (**Fig. 1a** and **Fig. 2d-h**). Our data therefore indicate a prominent role played by RAP80 in polyUb chain binding, consistent with the increased activity detected for ARISC–RAP80 over ARISC towards K63-Ub6 and longer chains as well as with the reduced activity observed upon deletion of the RAP80 UIM and ZnF domains (**Fig. 1c,f** and **Supplementary Fig. 1b,c**).

Collectively, these analyses describe the first high-resolution structure of ARISC–RAP80 bound to K63-linked ubiquitin chains. ARISC interacts with ubiquitin via a composite surface of three proteins: the BRCC36 active site of one ARISC monomer, and the Abraxas1* and BRCC45* subunits of the adjacent protomer. Substrate recognition is also aided by multiple domains in RAP80, especially in the context of long polyUb chains. This demonstrates that ARISC–RAP80 must assemble with a defined stoichiometry and undergo super-dimerisation to function as a single macromolecular unit that is capable of selectively recognising and cleaving K63-linked polyubiquitin substrates.

### Molecular mechanisms of BRCC36 engagement with diubiquitin

To understand the molecular details of how BRCC36 engages with ubiquitin chains at its active site, and compare it with other JAMM/MPN DUBs, we used the ubiquitin-bound BRCC36 protomer present in our human ARISC(E33A)–RAP80:K63-Ub7 structure and overlayed this with previously determined BRCC36 structures from metazoan orthologues (mouse, *Mus musculus*; zebrafish, *Danio rerio*; ant, *Camponotus floridanus*)^35,37^. Similarities were observed across all models (**Supplementary Fig. 8a**, *left panel*) and, notably, the active site residues and corresponding secondary structural elements in human BRCC36 were all captured in an active-like configuration (**Supplementary Fig. 8a**, *left panel*). Analogous comparisons between human BRCC36 MPN^+^ and the corresponding domain of AMSH-LP, CSN5, and RPN11^18,52,53^ revealed shared positioning of both active site residues and secondary structural elements between all JAMM/MPN DUBs, suggesting a common mechanism for JAMM domain substrate cleavage (**Supplementary Fig. 8b-d**).

Next, we examined the mode of BRCC36 engagement with Ub (P1) and Ub (P1’), and compared it to the structure of AMSH-LP bound to K63-Ub2^18^. While the models showed high structural similarities, notable differences in the orientation of Ub (P1) and, to a greater extent, of Ub (P1’) were apparent (**Fig. 3a**). In the AMSH-LP:K63-Ub2 structure, α1 of Ub (P1) is shifted of ∼ 6.5 Å towards the AMSH-LP catalytic domain compared to the corresponding molecule in our ubiquitin-bound BRCC36 model (**Fig. 3a**). Re-positioning of these secondary structural elements results in a ∼ 2.7 Å dip of the Ub (P1) tail in the AMSH-LP model relative to our structure (**Fig. 3a**), causing the C-terminus of Ub (P1) to be nearer the AMSH-LP catalytic domain compared to BRCC36. In addition, the orientation of Ub (P1’) is strikingly different between the AMSH-LP and BRCC36 models, with a measured ∼ 25° rotation and ∼ 14 Å shift of α1 from the Ub (P1’)-bound AMSH-LP molecule relative to the corresponding structural elements of Ub (P1’) in BRCC36 (**Fig. 3a**). Consequently, Ub (P1’) is positioned ∼ 12.7 Å higher in the AMSH-LP:K63-Ub2 structure than the corresponding molecule in our ARISC(E33A)–RAP80:K63-Ub7 model (**Fig. 3a**). These analyses strongly underline the inherent plasticity of ubiquitin molecules even when connected with the same linkage type^11^.

**Fig. 3:**
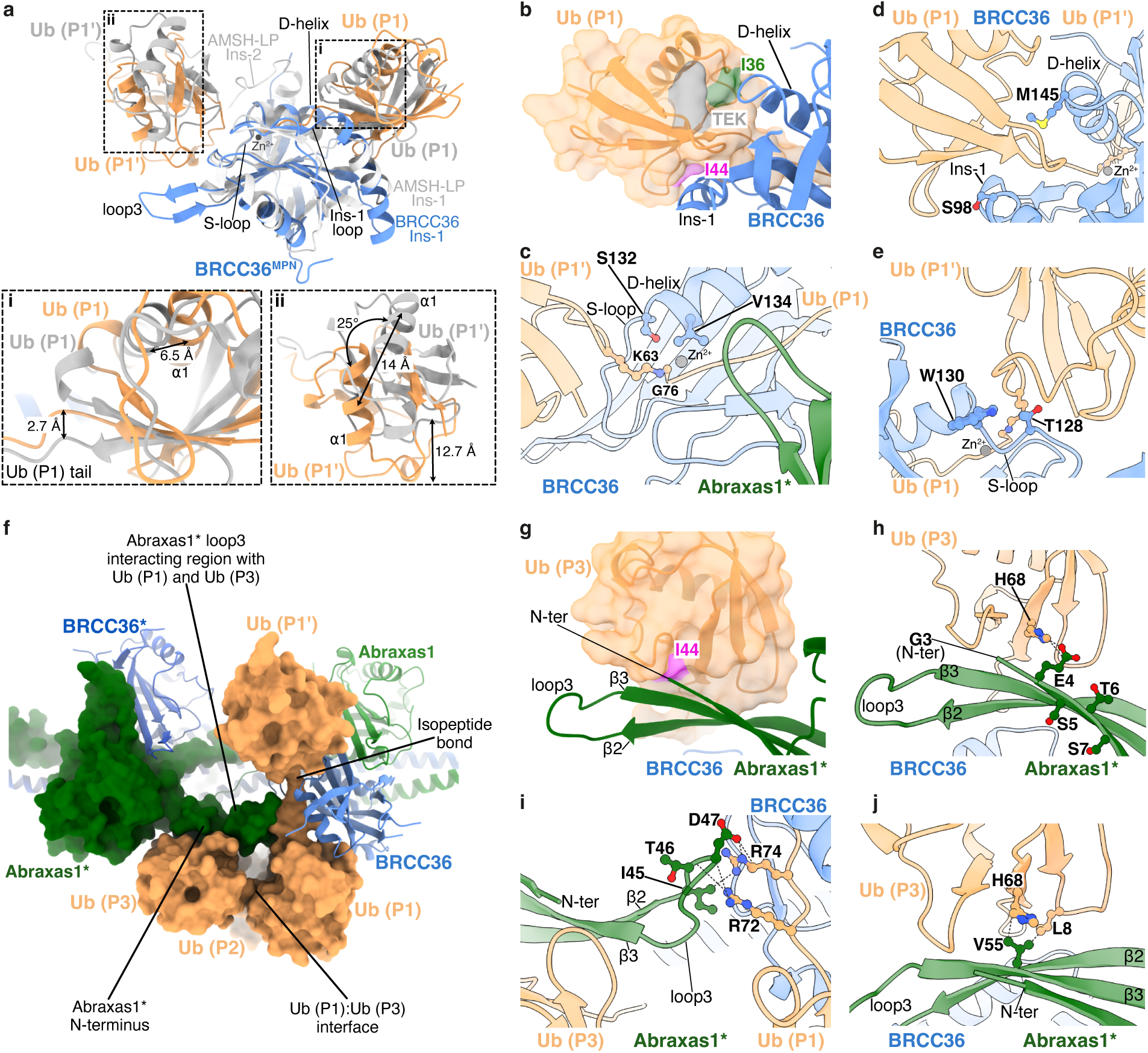
Structural analyses of the BRCC36 and Abraxas1 ubiquitin binding sites. **a**, Overlays between the structures of BRCC36^MPN^:K63-Ub2 (from the ARISC(E33A)–RAP80:K63-Ub7 complex; light blue and orange) and AMSH-LP^MPN^:K63-Ub2 (PDBid = 2ZNV; light grey). Structures are shown as cartoon, with a representative Zn^2+^ atom from ARISC(E33A)–RAP80:K63-Ub7 depicted as a grey sphere. Dashed black rectangles highlight divergent regions between the two models (*top*). Close-up views and structural details of Ub (P1) (i) and Ub (P1’) (ii). Measured distances highlight differences in the positioning of the ubiquitin moieties between the two structures (*bottom*). MPN, Mpr1, Pad1 N-terminal; Ins, insertion. **b,** Close-up view of the BRCC36:Ub (P1) interface. BRCC36 and Ub (P1) are shown as cartoon, and colored as in Fig. 1a and Fig. 2a. The I36 and I44 patches, and the TEK box are mapped onto the Ub (P1) surface and colored as indicated. **c-e,** Close-up views and structural details of the BRCC36:Ub (P1):Ub (P1’) (**c**), BRCC36:Ub (P1) (**d**), and BRCC36:Ub (P1’) (**e**) interfaces. Interacting residues and the Ub (P1):Ub (P1’) isopeptide bond are shown as ball & sticks and Zn^2+^ as a grey sphere. **f,** Close-up view of the Abraxas1*:Ub (P3):Ub (P1) interfaces. Abraxas1* and all ubiquitin molecules are shown as surface representation, while Abraxas1 and BRCC36/BRCC36* are depicted as cartoon. **g,** Close-up view of the Abraxas1*:Ub (P3) interface. The I44 hydrophobic patch is mapped onto the Ub (P3) surface and colored as indicated. **h-j,** Close-up views and structural details of the Abraxas1* N-terminus:Ub (P3) (**h**), Abraxas1* loop3:Ub (P1) (**i**), and Abraxas1* β2-loop3-β3:Ub (P3) (**j**) interfaces. Interacting residues are shown as ball & sticks; direct contacts between amino acids are indicated with dashed black lines. N-ter, N-terminus.

By contrast, Ins-1 and Ins-2, which play a prominent role in modulating ubiquitin binding^18^ and in coordinating the isopeptide bond, are similar in both AMSH-LP and BRCC36 as is the corresponding D-helix (**Fig. 3a,c,d** and **Supplementary Fig. 8b**). Notably, productive positioning of Ub (P1) into the BRCC36 active site relies on interactions between BRCC36 Ins-1 and the I44 hydrophobic patch in Ub (P1) as well as on the BRCC36 D-helix making extensive contacts with Ub (P1) I36 patch and TEK box^11^ regions (**Fig. 3b**). In contrast to the Ub (P1) interface, Ins-2 of AMSH-LP makes smaller contacts with Ub (P1’) (**Fig. 3a** and **Supplementary Fig. 8b**). The corresponding region in BRCC36 is not conserved across species and, in humans, forms an extended loop of 25 amino acids (residues 184-208) (**Supplementary Fig. 8a**, *left panel*)^41^. We could not observe the full BRCC36 Ins-2 in the ARISC(E33A)–RAP80:K63-Ub7 structure, and our XL-MS analyses identified few BRCC36 cross-links only with cyclical and linear K63-Ub7 substrates (**Fig. 2a,d-h**, **Supplementary Fig. 5c,h**, and **Supplementary Fig. 8a**, *left panel*). These observations therefore suggest that, in contrast to AMSH-LP, BRCC36 Ins-2 does not play a prominent role in ubiquitin binding. Despite these differences, BRCC36 engagement with Ub (P1’) also occurs via a smaller interface largely consisting of the BRCC36 S-loop and loop3 regions contacting Ub (P1’) loop2 and loop5 (**Fig. 3a,e**). Interestingly, interactions between loop3 and Ub (P1’) appear to be specific for BRCC36, as this region (residues 155-167) is shorter and not as extended in the other JAMM/MPN DUBs (**Fig. 3a** and **Supplementary Fig. 8e,f**). Our analyses therefore highlight how different JAMM MPN DUBs use diverse secondary structure elements to promote ubiquitin chain binding in their catalytic core.

### BRCC36:ubiquitin interactions are critical for DUB function

In our ARISC(E33A)–RAP80:K63-Ub7 cryo-EM structure, the local resolution at the BRCC36 interface with Ub (P1) and Ub (P1’) is ∼ 2.0 Å (**Supplementary Fig. 6b,c**), which is sufficient to visualise amino acid side chains in BRCC36 that contribute to ubiquitin binding in the active site. BRCC36 residues S132 and V134, localised in the S-loop and at the D-helix N-terminus, coordinate the isopeptide bond between Ub (P1) and Ub (P1’) (**Fig. 3c**).

By contrast, amino acids S98 and M145, which localise in Ins-1 and at the D-helix C-terminus, interact with Ub (P1) while S-loop residues T128 and W130 engage with Ub (P1’) (**Fig. 3d,e**). To validate our model and determine the contribution of each interacting residue to ARISC DUB activity, we mutated selected amino acids in BRCC36 and purified five ARISC variant complexes from insect cells (**Supplementary Fig. 9a**). We then assayed the resulting mutant proteins, together with ARISC WT, against K63-linked Ub2, Ub4, and Ub7 chains. Consistent with the function of each interacting residue in coordinating ubiquitin binding in the BRCC36 active site, all mutants tested abolished or strongly decreased ARISC enzymatic activity (**Fig. 4a** and **Supplementary Fig. 9b**). Next, we sought to determine the contribution of BRCC36 to ubiquitin chain binding. We introduced the S98K mutation in our inactive ARISC(E33A)–RAP80 complex to prevent cleavage of ubiquitin chains, and to discriminate between enzymatic activity and substrate binding. We then tested the resulting variant (here referred to as ARISC(E33A)^BRCC36(S98K)^–RAP80), alongside ARISC(E33A)–RAP80, in spectral shift assays to measure interactions with K63-linked Ub6 chains (**Fig. 4b**). ARISC(E33A)–RAP80 bound to K63-Ub6 with a K_d_ of ∼ 0.2 µM, which was strongly reduced in the presence of the ARISC(E33A)^BRCC36(S98K)^–RAP80 mutant (**Fig. 4c**). These data are consistent with our earlier enzymatic assays (**Fig. 4a** and **Supplementary Fig. 9b**), and indicate a prominent role for BRCC36 in supporting ARISC–RAP80 binding to ubiquitin chains.

**Fig. 4:**
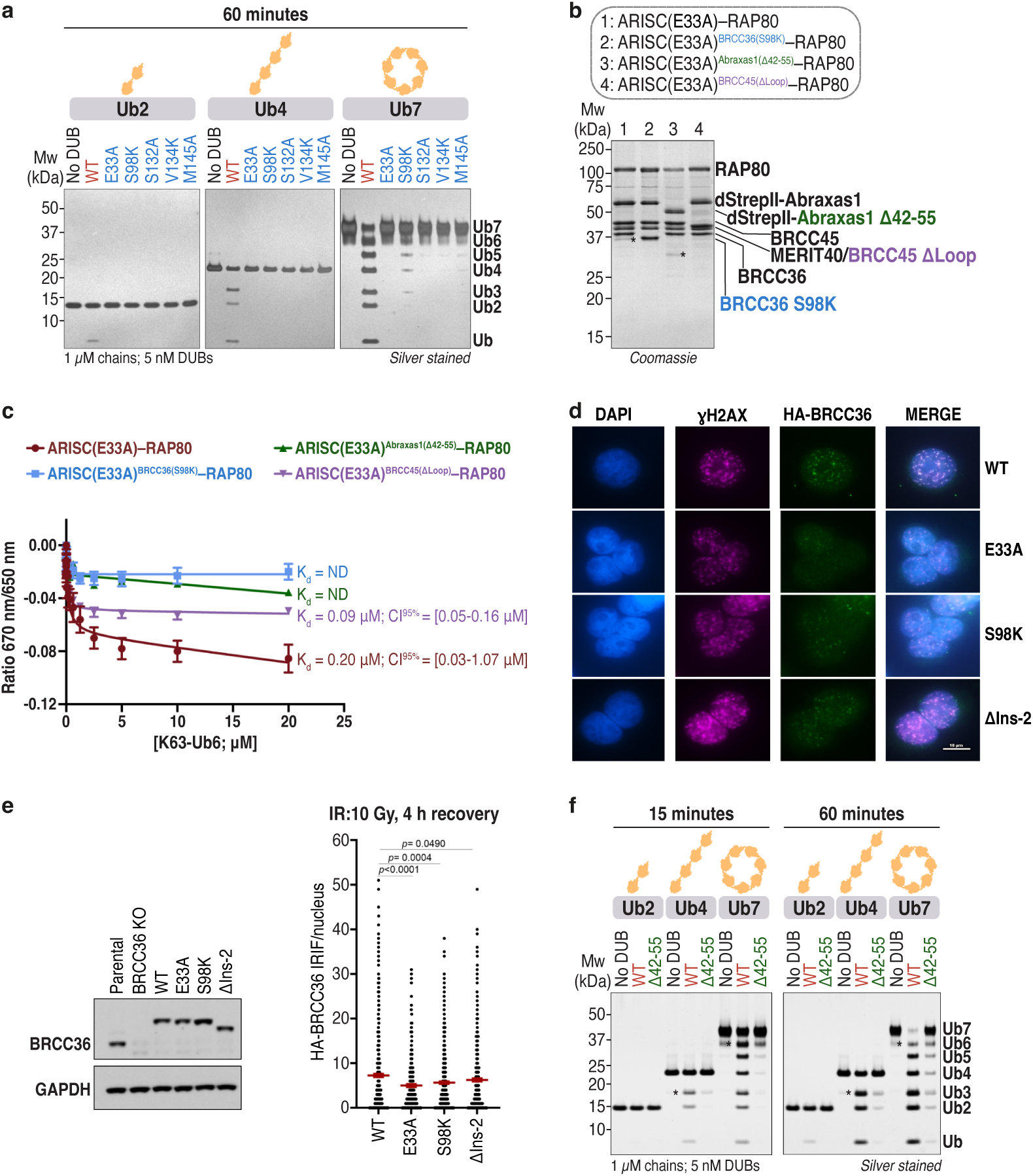
BRCC36 and Abraxas1 support ARISC interaction with ubiquitin chains. **a**, K63-Ub2, -Ub4, and -Ub7 chains (1 µM) were incubated with ARISC WT or the indicated ARISC variants (5 nM) for 60 minutes. Cleavage activity was analysed by SDS-PAGE and silver staining. Data are representative of two independent experiments. **b,** SDS-PAGE analysis of ARISC(E33A)–RAP80, ARISC(E33A)^BRCC36(S98K)^–RAP80, ARISC(E33A)^Abraxas1(Δ42-55)^–RAP80, and ARISC(E33A)^BRCC45(ΔLoop)^–RAP80. dStrepII, double StrepII tag. * indicates Abraxas1 degradation product. **c,** Spectral shift assays measuring binding of labelled ARISC(E33A)–RAP80 or the indicated mutant complexes (40 nM) to cyclical K63-Ub6 chains (20 µM-0 µM). Data points are mean ± SEM of two independent experiments carried out in technical duplicates. Dissociation constants (K_d_) are indicated; CI, confidence interval. **d,** Representative images of WT or mutants BRCC36 IRIF in HT-29 cells 4 h post irradiation (10 Gy). Scale bar is 10 µm. **e,** Western blots showing BRCC36 protein levels in HT-29 cells reconstituted with WT or mutants BRCC36 as indicated (l*eft*). Scatter plot showing quantification of the BRCC36 IRIF described in **d**. Data represent mean ± SEM derived from n ≥ 300 nuclei examined over two independent experiments; *p* values are indicated, unpaired two-tailed t test (*right*). **f,** K63-Ub2, -Ub4, and -Ub7 chains (1 µM) were incubated with ARISC WT or ARISC Δ42-55 (Abraxas1 Δ42-55) (5 nM) for up to 60 minutes. Cleavage activity was analysed as in **a**. Data are representative of two independent experiments. DUB, deubiquitylating enzyme; WT, wild type; Ub, ubiquitin.

To assess the importance of BRCC36 interaction with ubiquitin chains in cells, we tested a subset of our BRCC36 mutants for their ability to form ionising radiation-induced foci (IRIF) relative to WT protein. In agreement with our *in vitro* assays, the Ins-1 mutant S98K demonstrated a reduction of BRCC36 IRIF levels that was comparable to the inactive E33A variant (**Fig. 4d,e**). Conversely, deletion of the human-specific BRCC36 Ins-2 loop did not cause considerable reduction in IRIF levels when compared to WT protein (**Fig. 4d,e**). These data indicate that BRCC36-mediated recruitment to ubiquitylated chromatin is lost in the DUB-inactive E33A variant^40^, and in the BRCC36 S98K mutant where engagement with ubiquitin chains in its active site is impaired.

### Abraxas1* is essential for polyubiquitin chain engagement

The role of BRCC36 in promoting ubiquitin chain engagement within ARISC extends beyond the active site-bound ubiquitin molecules. In our ARISC(E33A)–RAP80:K63-Ub7 structure, the C-terminus of BRCC36 Ins-1 and α4 of BRCC36 coiled coil1 (CC1) contact the TEK box and F4 patches of Ub (P2). In addition, the loop connecting BRCC36 CC1 and CC2 is in proximity of the Ub (P2):Ub (P3) isopeptide bond (**Supplementary Fig. 9c,d**). Engagement of ARISC with Ub (P2) is also supported by the Abraxas1 protomer, which positions its CC loop in closer proximity to the Ub (P2) N-terminus (**Supplementary Fig. 9d**).

Similar to the ancestral CfAbraxas and HsAbraxas2^35,38^, human Abraxas1 lack catalytic function (**Supplementary Fig. 8a**, *right panel*). However, this subunit plays key structural roles within ARISC and ARISC–RAP80, where it uses its MPN^-^ and C-terminal regions to stabilise interactions with the BRCC36 MPN^+^ domain and the BRCC45-MERIT40-RAP80 arm^37,40^. In our cryo-EM structure, the Abraxas1* protomer is essential for ARISC–RAP80 engagement with the ubiquitin moieties at the distal end of the K63 chain. Specifically, the N-terminal (residues 1-10) and β3 regions of Abraxas1* bind Ub (P3) while loop3 coordinates the Ub (P1) tail into the active site of the adjacent BRCC36 protomer (**Fig. 3f**). These contacts are also supported by the I44 hydrophobic patch of Ub (P3), which grasps the N-terminus of Abraxas1 β3 in a manner reminiscent of the BRCC36 Ins-1:Ub (P1) interface (**Fig. 3b,g**). AlphaFold 3 modelling of the Abraxas1:ubiquitin sub-complex predicted ubiquitin to be positioned on the same Abraxas1 interface as seen in the cryo-EM structure, thus further validating our findings (**Supplementary Fig. 9e**). Sequence analyses revealed that both the N-terminal and β2-loop3-β3 regions share a high degree of sequence conservation across Abraxas1 species, with only some divergencies in the fish (*Salmon salar*), zebrafish (*Danio rerio*), and ant (*Camponotus floridanus*) orthologues (**Supplementary Fig. 9f**). By contrast, the analogous subunit of the cytoplasmic BRISC complex, Abraxas2, has a shorter N-terminus compared to human Abraxas1 and the two proteins display poor sequence conservation around the β2-loop3-β3 region (**Supplementary Fig. 9f**). These observations suggest that Abraxas1 has evolved specific ubiquitin-interacting surfaces required for ARISC-mediated engagement with K63-linked ubiquitin chains. Our analyses also indicate that BRCC36 and Abraxas1* loop3 regions share similar ubiquitin-binding properties (**Fig. 3a,f,h-j**, **Supplementary Fig. 8a**, *right panel*, and **Supplementary Fig. 9e**), allowing ARISC to stably interact with the proximal and distal end of a ubiquitin chain from adjacent interfaces within its BRCC36-Abraxas1 catalytic core.

The local resolution of our ARISC(E33A)–RAP80:K63-Ub7 cryo-EM structure around the Abraxas1* N-terminal and loop3 regions is ∼ 2.0-2.5 Å (**Supplementary Fig. 6b,c**), allowing the identification of residues that contribute to ubiquitin chain binding. In our model, E4 of Abraxas1* interacts with H68 of Ub (P3) (**Fig. 3f,h**) while amino acids 42-55 make extensive contacts with both Ub (P3) and residues located at the adjacent Ub (P1) tail (**Fig. 3f,i**). In particular, Abraxas1* I45, T46, and D47 make main- and side-chain contacts with R72 and R74 of Ub (P1) (**Fig. 3i**). Furthermore, V55, which is localised in Abraxas1* β3, interacts with L8 and H68 of Ub (P3) (**Fig. 3f,j**). To validate our structural model and assess the contribution of Abraxas1 β2-loop3-β3 residues to ARISC DUB activity, we deleted amino acids 42-55 and purified the resulting ARISC variant complex from insect cells (**Supplementary Fig. 9a**). Strikingly, Abraxas1* Δ42-55 displayed a strong reduction in ARISC DUB activity when compared to WT complex (**Fig. 4f**). To directly assess the importance of Abraxas1 β2-loop3-β3 region to ubiquitin chain binding, we introduced the same Abraxas1 Δ42-55 variant in the inactive ARISC(E33A)–RAP80 complex (here referred to as ARISC(E33A)^Abraxas1(^Δ42–55^)^–RAP80). We then tested the resulting mutant, alongside ARISC(E33A)–RAP80, in spectral shift assays against K63-Ub6 chains (**Fig. 4b**). In agreement with our earlier experiments, deletion of Abraxas1 residues 42-55 strongly reduced ARISC–RAP80 binding to ubiquitin chains *in vitro* (**Fig. 4c,f**).

To assess the relevance of the Abraxas1*:ubiquitin interface for ARISC–RAP80 function in cells, we generated a short and long deletion of the Abraxas1 β2-loop3-β3 region (Δ42-55 and Δ46-50, respectively) and assessed the ability of these variants to form Abraxas1 IRIF. We also deleted Abraxas1 residues 274-325, which are localised at the Abraxas1:RAP80 AIR interface and are critical for ARISC binding to RAP80^37,40^. Consistent with the established role of the Abraxas1 C-terminus in mediating RAP80 AIR interactions with ARISC, deletion of amino acids 274-325 strongly reduced Abraxas1 IRIF (**Supplementary Fig. 9g,h**). By contrast, we did not detect any considerable difference in IRIF levels between Abraxas1 WT and Abraxas1 Δ42-55 or Δ46-50 variants (**Supplementary Fig. 9g,h**). These data suggest that interactions between Abraxas1 and ubiquitin near the catalytic centre, although necessary for enzyme activity and substrate binding *in vitro*, they are not on their own sufficient for ARISC–RAP80 recruitment to DNA damage sites.

### BRCC45* provides a minor contribution to ubiquitin chain binding

In the structure of the ARISC(E33A)–RAP80:K63-Ub7 complex, the BRCC45* protomer does not make extensive contacts with the ubiquitin moieties bound to the catalytic site (**Supplementary Fig. 5c**). The only putative BRCC45* interface closer to the K63-Ub chain consists of BRCC45* F140 and residues 161-170 of UEV-M loop3, which are likely to be positioned in proximity to Ub (P4) (**Supplementary Fig. 5c,g** and **Supplementary Fig. 10a**). By contrast, amino acids Y263 and R320 in the BRCC45 UEV-C domain are predominantly involved in protein:protein interactions and positioned closer to the Abraxas1 C-termini (**Supplementary Fig. 10a**). To validate our model, and assess BRCC45* contribution to ubiquitin chain binding, we mutated or deleted selected BRCC45 residues and purified four ARISC variant complexes from insect cells (**Supplementary Fig. 9a**). We then assayed the resulting mutant proteins, together with ARISC WT, against K63-linked Ub2, Ub4, and Ub7 chains (**Supplementary Fig. 1a**, *left panel*). Consistent with our structural model, ARISC enzymatic activity was marginally decreased against Ub2 and Ub4 chains, particularly for the F140A, Δ161-170 (here referred to as ΔLoop^BRCC45^), and W165A mutants (**Supplementary Fig. 5c** and **Supplementary Fig. 10**). The ability of ARISC to hydrolyse K63-Ub7 was instead reduced in the presence of all BRCC45 variants (**Supplementary Fig. 10b**). To directly determine BRCC45 contribution to ubiquitin chain binding, we introduced the ΔLoop^BRCC45^ variant in our inactive ARISC(E33A)–RAP80 complex (here referred to as ARISC(E33A)^BRCC45(ΔLoop)^–RAP80). We then assayed the resulting mutant, together with ARISC(E33A)–RAP80, in spectral shift assays against K63-Ub6 chains (**Fig. 4b**). Deletion of residues 161-170 in BRCC45 reduced ARISC–RAP80 binding to ubiquitin chains, albeit to a lesser extent compared to the BRCC36 and Abraxas1 variants (**Fig. 4c**). These data are consistent with our positioning of Ub (P4) in the low-resolution ab-initio map and earlier XL-MS analyses, and indicate that BRCC45* F140 and the UEV-N loop region likely mediate some contacts with Ub (P4) (**Supplementary Fig. 5c,g** and **Supplementary Fig. 10a**). In our model, BRCC45* residues Y263 and R320 are positioned in proximity to the Abraxas1 C-termini, but do not mediate ubiquitin binding (**Supplementary Fig. 10a**). This indicates that the moderate reduction in ubiquitin chain hydrolysis observed for the Y263A+R320A double mutant is likely due to decreased ARISC complex assembly as a result of destabilised BRCC45*:Abraxas1* and, to a lesser extent, BRCC45*:Abraxas1*:RAP80 AIR* interfaces (**Supplementary Fig. 10**).

### Conserved mechanisms promote ARISC:polyubiquitin interaction

Our biochemical analyses demonstrate that RAP80 enhances ARISC activity, particularly in the presence of Ub6 and longer chains (**Fig. 1c** and **Supplementary Fig. 1b,c**), but do not provide information on which regions within ARISC are required for sustained DUB function. To understand the molecular determinants of ARISC enzymatic activity, we purified stoichiometric amounts of three ARISC variants, in which the N-terminus of MERIT40 (residues 1-71) and/or the C-terminal region of Abraxas1 (residues 266-409) were deleted (here indicated as ARISCΔN, ARISCΔC, and ARISCΔNΔC, respectively) (**Supplementary Fig. 11a**). We then compared the enzymatic activities of these ARISC variants, and of FL ARISC, against purified K63-Ub2, -Ub4, and -Ub7 chains (**Supplementary Fig. 1a**, *left panel*). We detected comparable levels of DUB activity for FL, ΔC, and ΔNΔC ARISC variants in the presence of all chain lengths while the activity of ARISCΔN was marginally reduced (**Supplementary Fig. 11b**). These results indicate that all ARISC constructs are active DUBs and suggest that the N-terminus of MERIT40 may play a potential role in modulating ARISC enzymatic activity. In addition, our biochemical analyses identify ARISCΔC as the minimally deleted complex with comparable levels of DUB activity as FL ARISC.

To determine how ARISC interacts with cyclical and linear ubiquitin chains and compare the mechanisms of ARISC-mediated ubiquitin binding with the ones observed for ARISC–RAP80, we performed single-particle cryo-EM analyses. We used our ARISCΔC(E33A) variant and set up cryo-EM grids with linear K63-Ub4 and cyclic K63-Ub7 chains as described for ARISC(E33A)–RAP80 (**Supplementary Fig. 5b**, *middle and right panels*)^46^. We collected two large datasets and identified structures containing the same U-shaped architecture previously reported^36,37^, as well as ubiquitin-like densities similar to the ones observed for ARISC–RAP80 (**Supplementary Fig. 12a,b** and **Supplementary Fig. 13a,b**). Following 3D refinements and postprocessing, we obtained consensus cryo-EM maps at overall resolutions of 3.2 Å for both complexes (**Supplementary Fig. 12b,c**, **Supplementary Fig. 13b,c**, and **Supplementary Table 1**). To improve particle distribution and attenuate preferential orientations (**Supplementary Fig. 7h,m**), we performed focused masking and local refinements on different regions of both maps. This allowed the generation of final composite maps with local resolutions between 3.41-3.70 Å for ARISCΔC(E33A):K63-Ub7 and 3.53-3.88 Å for ARISCΔC(E33A):K63-Ub4 (**Supplementary Fig. 7i-l** and **n-p, Supplementary Fig. 12b,d-h, Supplementary Fig. 13b,d-f**, and **Supplementary Table 1**).

We observe the highest resolution in the core of the ARISCΔC(E33A):K63-Ub7 and ARISCΔC(E33A):K63-Ub4 maps, consisting of the BRCC36-Abraxas1ΔC superdimer (**Fig. 5a,b**, *left panels*, **Supplementary Fig. 12b,c**, and **Supplementary Fig. 13b,c**). The resolution is lower around the BRCC45 UEV-N domain and, in agreement with the intrinsic flexibility observed for the ARISC and BRISC arm regions^37,38,40,41^, we visualise no density corresponding to BRCC45 UEV-M and UEV-C nor to MERIT40 (**Fig. 5a,b**, *left panels*, **Supplementary Fig. 12b,c**, and **Supplementary Fig. 13b,c**). Local refinements using masks comprising the BRCC45 UEV-M domain partially improved the densities of this region in both maps, allowing us to model build BRCC45 UEV-N and rigid-body fit the BRCC45 UEV-M domain from our model of the ARISC(E33A)–RAP80:K63-Ub7 complex (**Fig. 5a,b**, **Supplementary Fig. 12b,d,e**, and **Supplementary Fig. 13b,d,e**).

**Fig. 5:**
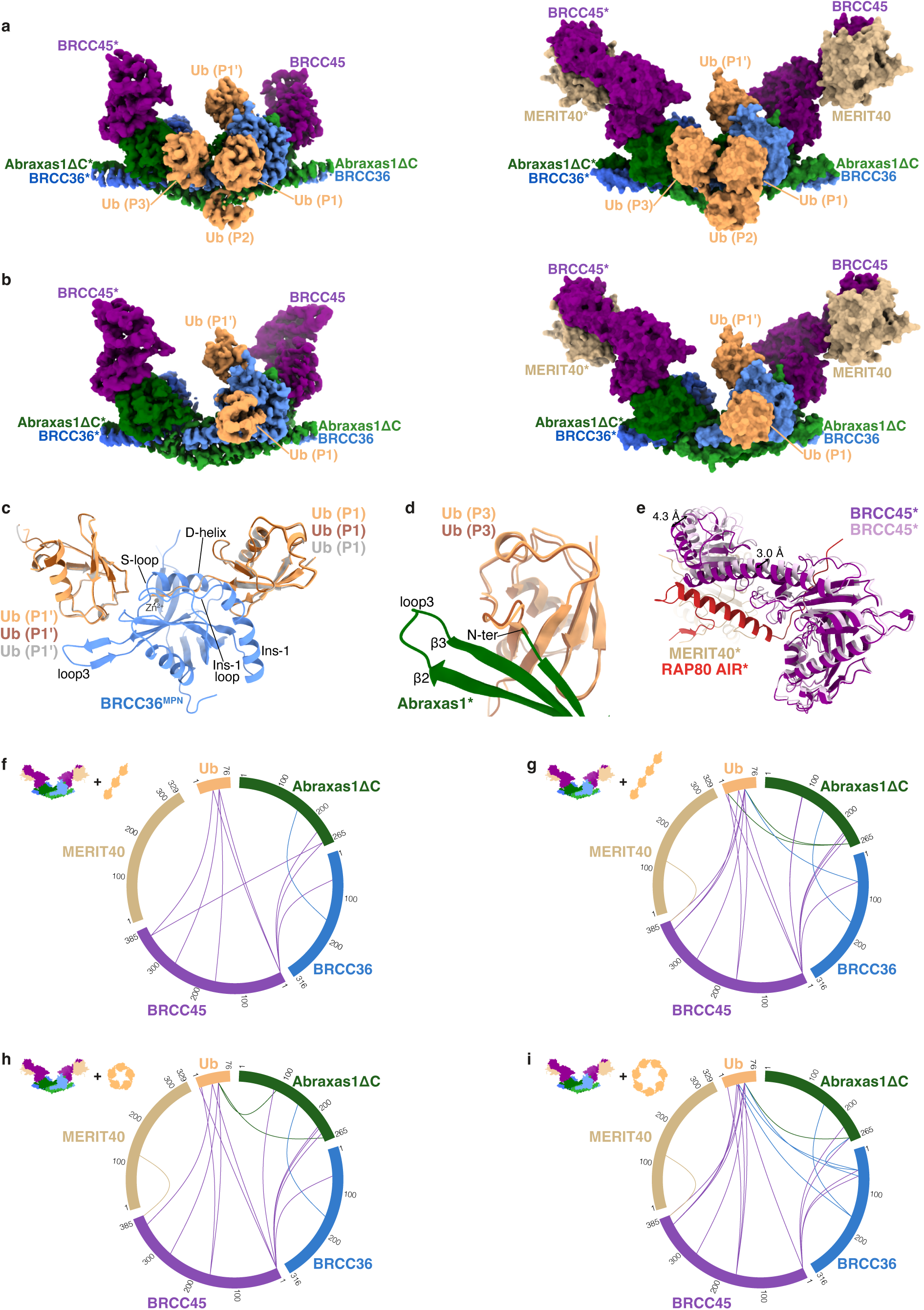
Cryo-EM structures of the ARISCΔC(E33A):K63-Ub7 and ARISCΔC(E33A):K63-Ub4 complexes. **a,b**, Cryo-EM composite maps of ARISCΔC(E33A):K63-Ub7 (**a**) and ARISCΔC(E33A):K63-Ub4 (**b**) at contour levels of 0.0315 and 0.0372 (*left*). The same structures are shown as surface representation (*right*). Owing to low resolution in the extended arm regions, overlayed BRCC45 UEV-C and MERIT40 models are shown for size comparison only. ARISC subunits and ubiquitin chains are colored as in Fig. 1a and Fig. 2a. **c,** Overlays between the cryo-EM structures of BRCC36^MPN^:K63-Ub2 (from ARISC(E33A)–RAP80:K63-Ub7; light blue and orange), ARISCΔC(E33A):K63-Ub7 (light brown), and ARISCΔC(E33A):K63-Ub4 (grey). The BRCC36 MPN domains from the ARISCΔC(E33A):K63-Ub7 and ARISCΔC(E33A):K63-Ub4 structures were omitted for clarity. Structures are shown as cartoon, with a modelled Zn^2+^ atom depicted as a grey sphere. Ins-1, insertion-1; MPN, Mpr1, Pad1 N-terminal. **d,** Close-up view of the Abraxas1*:Ub (P3) interface obtained by overlaying the cryo-EM structures of ARISC(E33A)–RAP80:K63-Ub7 (green and orange) and ARISCΔC(E33A):K63-Ub7 (light brown). The Abraxas1* model from the ARISCΔC(E33A):K63-Ub7 structure was omitted for clarity; structures are shown as cartoon. N-ter, N-terminus. **e,** Overlays between the BRCC45 molecules from ARISC(E33A)–RAP80:K63-Ub7 (purple, with MERIT40 and RAP80 AIR colored as in Fig. 1a) and ARISCΔC(E33A):K63-Ub7 (light pink). Owing to low resolution in the extended arm regions of the ARISCΔC(E33A):K63-Ub7 structure, the BRCC45 UEV-C model is shown for overlay purposes only. Structures are shown as cartoon. AIR, Abraxas1-interacting region. **f-i,** Chemical cross-linking and mass spectrometry analyses of ARISCΔC(E33A) in complex with K63-linked Ub3 (**f**), Ub4 (**g**), Ub5 (**h**), and Ub7 (**i**) chains. Proteins are labelled and their corresponding inter-subunit cross-links color-coded accordingly. Communal cross-links obtained from two independent experiments are shown. Crosslink sites involving ubiquitin were searched and mapped against a single ubiquitin sequence. Ub, ubiquitin.

Next, we sought to identify the position of the ubiquitin chains in the ARISCΔC(E33A):K63-Ub7 and ARISCΔC(E33A):K63-Ub4 consensus maps. Supported by well-defined densities on either side of each BRCC36 molecule, we rigid-body fitted two K63-linked ubiquitin moieties into the Ub (P1) and Ub (P1’) sites of both maps (**Fig. 5a,b**, **Supplementary Fig. 12b,c,f**, and **Supplementary Fig. 13b,c,f**). Further visualisation of the ARISCΔC(E33A):K63-Ub7 consensus map and of the best map obtained from heterogeneous refinement revealed additional densities extending from Ub (P1) towards Abraxas1* (**Fig. 5a**, **Supplementary Fig. 11c**, and **Supplementary Fig. 12b,c**), and we were able to confidently place two K63-linked ubiquitin moieties in these Ub (P2) and Ub (P3) sites (**Fig. 5a** and **Supplementary Fig. 12b,g**). Subsequent visualisation of the same heterogeneous refinement map revealed a ubiquitin-like density extending from Ub (P3) towards BRCC45* (**Supplementary Fig. 11c**), which we could however not improve following local refinement. We therefore rigid-body fitted a fifth ubiquitin molecule in this putative Ub (P4) site of the best heterogeneous refinement map, as previously described for ARISC–RAP80 (**Fig. 5a** and **Supplementary Fig. 11c**), and hypothesised that this ubiquitin moiety, alongside two additional missing ubiquitin molecules in the K63-Ub7 chain, will likely extend from Ub (P3) towards Ub (P1’) (**Supplementary Fig. 11d**). Taken together, our ensemble of structures demonstrates that, like ARISC–RAP80, ARISC employs multi-subunit interactions to engage with K63-linked ubiquitin chains (**Fig. 2a** and **Fig. 5a**). Comparisons of our structural models revealed only small differences in BRCC36 engagement of Ub (P1) and Ub (P1’) between ARISCΔC and full-length ARISC–RAP80 complexes, indicating that cyclical and linear K63-linked ubiquitin chains contact the BRCC36 active site in a similar manner (**Fig. 5c**). Likewise, overlaying of ARISCΔC(E33A):K63-Ub7 and ARISC(E33A)–RAP80:K63-Ub7 models revealed only minor differences (less than 1.7 Å) between their respective Ub (P3) moieties, confirming a shared mode of Abraxas1-mediated ubiquitin binding between ARISC and ARISC–RAP80 (**Fig. 2a**, **Fig. 3f-j**, and **Fig. 5a,d**). Interestingly, we observed a ∼ 3.5 Å downward shift of the BRCC45 UEV-M and UEV-C domains in the ARISC(E33A)–RAP80:K63-Ub7 model compared to the corresponding regions in the ARISCΔC(E33A):K63-Ub7 structure (**Fig. 5e**). This is consistent with the intrinsic flexibility previously reported for the ARISC arms^37,40^, and suggests that ARISC–RAP80 may sample more “open” BRCC45-MERIT40 arm conformations compared to ARISC.

To independently validate our structural models, we performed XL-MS analysis of ARISCΔC(E33A) in the absence and presence of K63-linked Ub3, Ub4, Ub5, and Ub7 chains (**Fig. 5f-i**, and **Supplementary Fig. 11e**). Taking advantage of the XL-MS-cleavable cross-linker DSBU, we identified intermolecular cross-links between all ARISC subunits as well as several cross-links between ubiquitin and BRCC36, Abraxas1ΔC, and BRCC45 (**Fig. 5f-i** and **Supplementary Fig. 11e,f**). This allowed us to confirm the positioning of BRCC36 Ins-2, which is not fully visible in our model, at the interface between Ub (P1’) and Ub (P4), Abraxas1ΔC in proximity to Ub (P2) and Ub (P3), and BRCC45* UEV-M loop3 near Ub (P4) in ARISC. These analyses are therefore consistent with our cryo-EM data and with our earlier observations on the ARISC–RAP80 complex (**Fig. 2a**, **Fig. 5a,b**, **Supplementary Fig. 5c,g**, and **Supplementary Fig. 11c,f**).

Collectively, these data describe structures of ARISC bound to linear and cyclical K63-linked ubiquitin chains. Our analyses indicate a shared mode of substrate recognition by the catalytic BRCC36-Abraxas1 superdimer, and a supportive role of the BRCC45* subunit to engage with polyUb chains in both ARISC and ARISC–RAP80 assemblies.

### ARISC and ARISC–RAP80 favour distal ubiquitin hydrolysis

Our cryo-EM structures of ARISC and ARISC–RAP80 in complex with ubiquitin chains (**Fig. 2a** and **Fig. 5a,b**) answer outstanding questions on how substrate recognition is regulated by these large enzymatic assemblies, but do not clarify if the complexes prefer the distal or proximal end. A previous study showed that the mouse ARISC–RAP80 AIR complex preferentially hydrolyses K63-Ub4 from the distal end of the chain^37^.

To further assess how human ARISC and ARISC–RAP80 complexes orient ubiquitin chains, and compare the mode of substrate hydrolysis between the two enzymatic assemblies, we introduced a fluorescent label at the proximal end of a K63-Ub4 chain using the same strategy as for our distally labelled K63-Ub4 substrate (**Supplementary Fig. 2a-c**). ARISC–RAP80 displayed comparable activities towards proximally and distally labelled K63-Ub4 chains, while ARISC showed a partial reduction in ubiquitin chain hydrolysis when assayed against proximally labelled K63-Ub4 compared to the distally labelled substrate (**Fig. 6a,b**). In addition, we observed a time-dependent appearance of fluorescent Ub3, Ub2, and Ub cleavage products when incubating ARISC–RAP80 with the proximally labelled chain (**Fig. 6a**, *left panel*). By contrast, incubation of the same DUB complex with the distally labelled K63-Ub4 substrate resulted in the accumulation of fluorescent Ub as the predominant cleavage product, with only small sub-populations of Ub3 and Ub2 chains appearing over time (**Fig. 6a**, *right panel*). Similar results were also observed for ARISC, with cleavage pattern differences less apparent but consistent with the ARISC–RAP80 results (**Fig. 6b**). These data indicate that ARISC and ARISC–RAP80 preferentially cleave ubiquitin chains from their distal end, in agreement with previous findings (**Fig. 6c**)^37^. Given the appearance of Ub3 and Ub2 cleavage products in the presence of the proximally labelled chain, our data also indicate that both ARISC and ARISC–RAP80 are capable of ubiquitin hydrolysis in the middle of the chain albeit with reduced preference (**Fig. 6**). Therefore, ARISC and ARISC–RAP80 complexes can be classified as endo-DUBs but with a preference for the distal end of a ubiquitin chain.

**Fig. 6:**
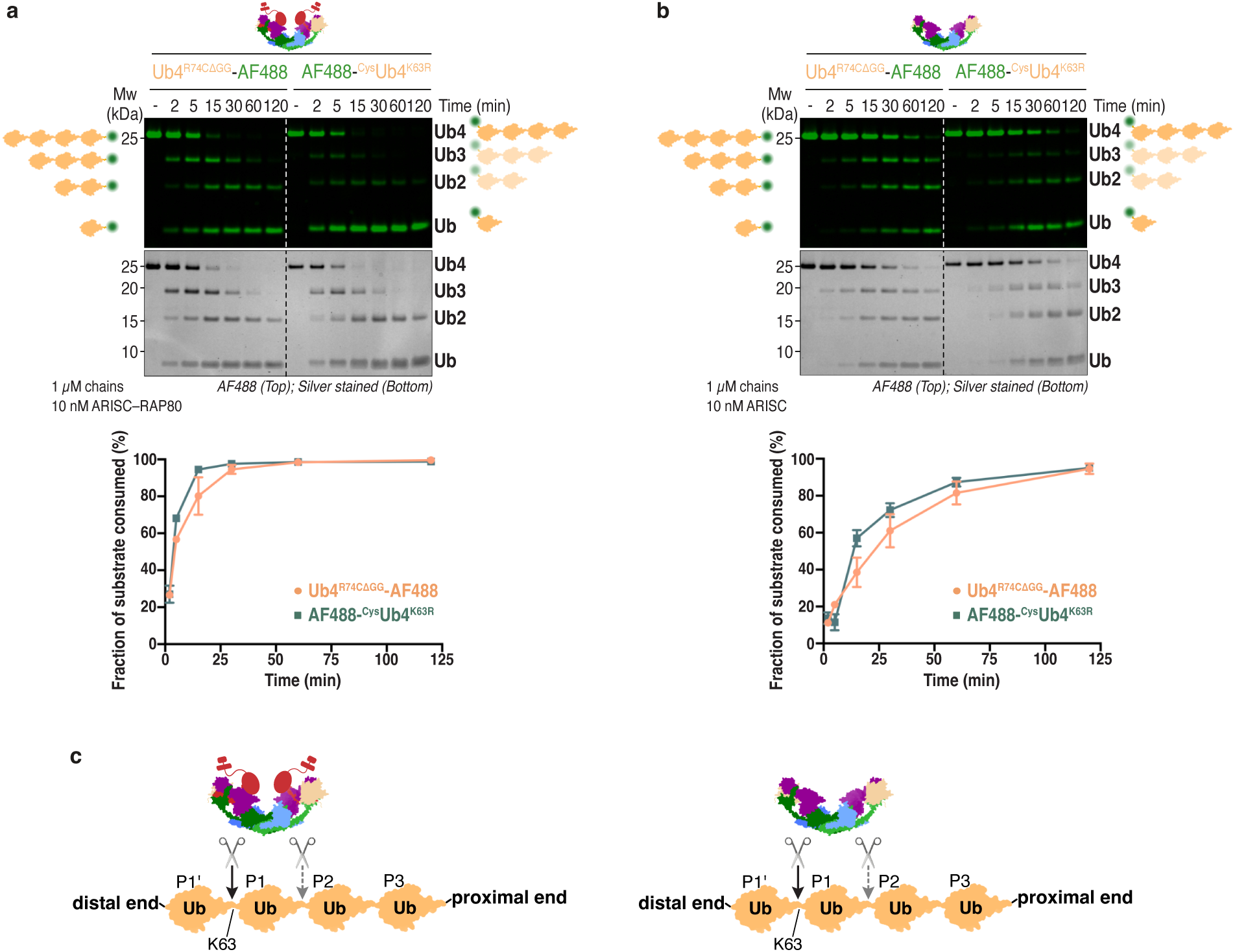
ARISC and ARISC–RAP80 are endo DUBs that preferentially cleave ubiquitin chains from the distal end. **a,b**, Gel-based ubiquitin chain cleavage assays. Alexa-Fluor 488 (AF488) labelled proximally (Ub4^R74CΔGG^-AF488) and distally (AF488-^Cys^Ub4^K63R^) blocked K63-Ub4 chains (1 µM) were incubated with ARISC–RAP80 (**a**) or ARISC (**b**) (10 nM) for up to 120 minutes. Cleavage activity was analysed as described in Fig. 1f (*top*). The disappearance of the K63-Ub4 parent band for each substrate was quantified using densitometry, and plotted as fraction of substrate consumed (%). Data points are mean ± SEM of two independent experiments (*bottom*). **c,** Schematics summarising the data shown in **a,b**. ARISC and ARISC–RAP80 complexes preferentially cleave ubiquitin chains from the distal end (black arrow), but also display activity towards the middle of the chain (dashed grey arrow).

### ARISC interacts with ubiquitin via non-catalytic sites

Our ubiquitin-bound ARISC–RAP80 structure shows four ubiquitin moieties surrounding the BRCC36 catalytic site (**Fig. 2a**). Upon further inspection of the consensus cryo-EM map, we noticed low-resolution globular densities on the top of each BRCC45 UEV-C domain that could not be attributed to BRCC45, MERIT40, the Abraxas1 C-termini, or RAP80 AIR (**Supplementary Fig. 14a,b**). Local refinements using masks focused on these regions modestly improved ARISC arm densities (**Supplementary Fig. 6b,h,i**), with the resulting globular features resembling a ubiquitin fold (**Supplementary Fig. 14a,b**). Given the ubiquitin-binding properties of the BRCC45-MERIT40 sub-complex^36,37^, we hypothesised that these densities represent ubiquitin moieties. Indeed, we could rigid-body fit ubiquitin monomers above each BRCC45 UEV-C domain in our ARISC(E33A)–RAP80:K63-Ub7 map (**Fig. 7a,b**). In agreement with these findings, XL-MS analyses of ARISC(E33A)–RAP80 and ARISCΔC(E33A) in the presence of all chain lengths and types revealed contacts between residues located in the BRCC45 UEV-C domain and ubiquitin (**Fig. 2d-h** and **Fig. 5f-i**). The majority of interactions centre around BRCC45 residues K270, K360, and K345, which appear to be positioned in proximity to M1, K6, K11, K29, K33, and K48 of ubiquitin (**Fig. 7c**). Strikingly, AlphaFold 3 modelling of the BRCC45:ubiquitin sub-complex predicted four of the five models to be characterised by an interface that closely matched the one observed in our structure (**Fig. 7d** and **Supplementary Fig. 14c**). In the fifth model, ubiquitin occupies an interface overlapping with BRCC45:Abraxas1, rendering it incompatible with ARISC binding (**Fig. 2a**, **Fig. 7a**, and **Supplementary Fig. 14c**). Our two-pronged approach using experimental evidence and AlphaFold predictions therefore strongly suggests that ARISC–RAP80 contacts ubiquitin via the BRCC45 UEV-C domains, which we refer to as non-catalytic ubiquitin sites (**Fig. 7a,b**).

**Fig. 7:**
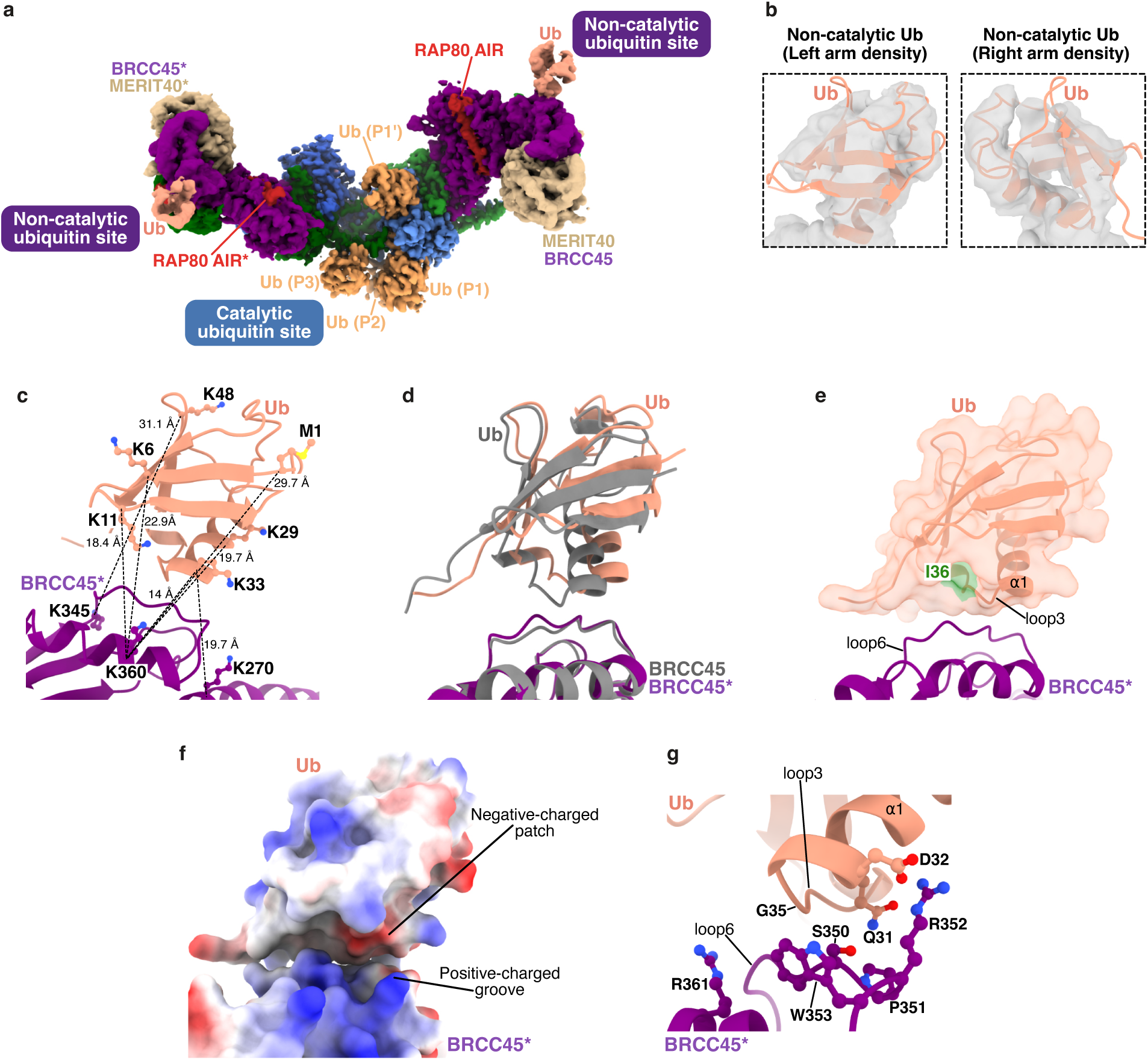
ARISC–RAP80 interacts with ubiquitin via catalytic and non-catalytic sites. **a**, Cryo-EM composite map of ARISC(E33A)–RAP80:K63-Ub7 at contour level of 0.0432, with ARISC–RAP80 subunits colored as in Fig. 1a. Ubiquitin molecules bound to catalytic (BRCC36 active site) and non-catalytic (BRCC45 UEV-C) sites are colored as in Fig. 2a and salmon. AIR, Abraxas1-interacting region. **b,** Close-up view of the densities corresponding to the non-catalytic ubiquitin sites on the left and right ARISC arms (see **Supplementary Fig. 14a,b**). Cartoon models of ubiquitin were rigid-body fitted into the cryo-EM map and colored as in **a**. **c**, Close-up views and structural details of the non-catalytic BRCC45*:Ub interface. Residues involved in cross-links are shown as ball & sticks. Distances are indicated with dashed black lines. **d,** Structural overlays between the non-catalytic BRCC45*:Ub interface as seen in the ARISC(E33A)–RAP80:K63-Ub7 structure (purple and salmon) and the corresponding interface predicted in AlphaFold 3 model 1 (grey). Models are shown as cartoon, and colored as in Fig. 1a and a. **e,** Close-up view of the non-catalytic BRCC45*:Ub interface. The hydrophobic I36 patch is mapped onto the ubiquitin surface and colored as indicated. **f,** Electrostatic potential molecular surfaces of the non-catalytic BRCC45*:Ub interface. The BRCC45* and ubiquitin models are oriented as in **e**, and the solvent-accessible surfaces displayed with a contour of ± 10.0 kT/e. **g,** Close-up views and structural details of the non-catalytic BRCC45*:Ub interface. Putative interacting residues are shown as ball & sticks. Ub, ubiquitin.

In our cryo-EM structure, ubiquitin utilises the C-terminus of α1 and the adjacent loop3 region, including its hydrophobic I36 patch, to contact loop6 of each BRCC45 UEV-C domain (**Fig. 7a,d,e**). In E2∼ubiquitin conjugates (E2∼Ub), E2 enzymes employ the same interface to interact with the bound ubiquitin (**Supplementary Fig. 14d**), indicating a conserved ubiquitin-interacting function of this region across UEV-containing proteins. Interestingly, residues 345-368 of BRCC45 form a positively-charged groove that sustains the complementary negatively-charged region (comprising amino acids 32-36) in ubiquitin (**Fig. 7f**). Detailed analyses of the BRCC45:ubiquitin interface showed that BRCC45 residues S350, P351, R352, W353, and R361 are located in closed proximity to amino acids Q31, D32, and G35 of ubiquitin (**Fig. 7a,g**).

To assess the importance of these non-catalytic ubiquitin sites for ARISC–RAP80 function in cells, we mutated selective amino acids at the BRCC45*:ubiquitin interface (**Fig. 7g**) and tested the resulting variants (S350Y+P351N+R352D and W353A+R361A) for their ability to form BRCC45 IRIF. Similarly to the results obtained for Abraxas1 Δ42-55 and Δ46-50, we did not detect any difference in IRIF levels between BRCC45 WT and BRCC45S^350Y+P351N+R352D^ or BRCC45^W353A+R361A^ variants (**Supplementary Fig. 9g,h** and **Supplementary Fig. 14e,f**). These results indicate that interactions between BRCC45 UEV-C and ubiquitin, although clearly present in our cryo-EM structure and XL-MS analyses, are not necessary for ARISC–RAP80 recruitment to damaged chromatin. It remains possible that the BRCC45 UEV-C domains provide low-affinity surfaces for mono and/or polyubiquitin chains, fine-tuning chain engagement or avidity in contexts not captured by our recruitment assays.

## Discussion

Deubiquitylating enzymes control a variety of cellular processes by hydrolysing ubiquitin chains or removing ubiquitin from substrate proteins^1^. The ARISC–RAP80 complex, which regulates DNA damage repair via homologous recombination (HR), acts selectively towards K63-linked ubiquitin polymers^27,28^. However, the molecular mechanisms underpinning ARISC–RAP80 recognition and cleavage of K63-linked chains remain elusive. By employing a combination of structural, biochemical, and cell-based approaches, we have determined how ARISC and ARISC–RAP80 decode K63-linked ubiquitin signalling, with important implications on BRCA1-A DUB function in DNA damage repair.

ARISC and ARISC–RAP80 engage with ubiquitin chains via multi-subunit interactions within the ARISC catalytic site. We verified substrate engagement by designing structure-guided mutants that failed to interact with K63-linked chains (**Fig. 3c,d,i,j**, **Fig. 4a-c,f**, **Supplementary Fig. 9a,b**, and **Supplementary Fig. 10**). Cell-based analyses confirmed the observed mode of BRCC36-mediated ubiquitin chain binding is important for ARISC–RAP80 recruitment to DNA damage sites (**Fig. 4d,e**). Substrate recognition and enzymatic activity are also supported by multiple domains in RAP80 (**Fig. 2d-h**), which preferentially interact with long, over short, K63-linked ubiquitin chains. A key finding is that, *in vitro*, ARISC–RAP80 mediates ubiquitin recognition via non-catalytic sites located on the BRCC45 UEV-C domains (**Fig. 7a-e,g**). This suggests that both RAP80 and BRCC45 contribute to ARISC–RAP80 recruitment to ubiquitylated chromatin, thereby ensuring timely and coordinated recognition of signals for DSB repair. A clear rationale for two recognition modules remains to be established, however, one possibility is a division of labour in which RAP80 engages branched SUMO-ubiquitin chains via its tandem SIM-UIM domains, while BRCC45 provide low-affinity binding surfaces for recognition of homotypic chains or monoubiquitin modifications.

Based on published works implicating ubiquitylated H2A and phosphorylated H2AX (ɣH2AX) as substrates for ARISC–RAP80/BRCA1-A^29–31^, we propose that targeting of the BRCA1-A complex to polyubiquitylated chromatin requires different domains in RAP80 and low-affinity, non-catalytic ubiquitin sites in the BRCC45 arms (**Fig. 8**). This mechanism would promote BRCA1-A recruitment to chromatin in the presence of DNA damage, while allowing continuous ubiquitin chain cleavage by the ARISC catalytic core. Our findings therefore describe how BRCA1-A can fine-tune ubiquitin-dependent signalling in the DNA damage response (DDR) by combining its multiple substrate recognition modules with its ubiquitin chain cleavage activity.

**Fig. 8:**
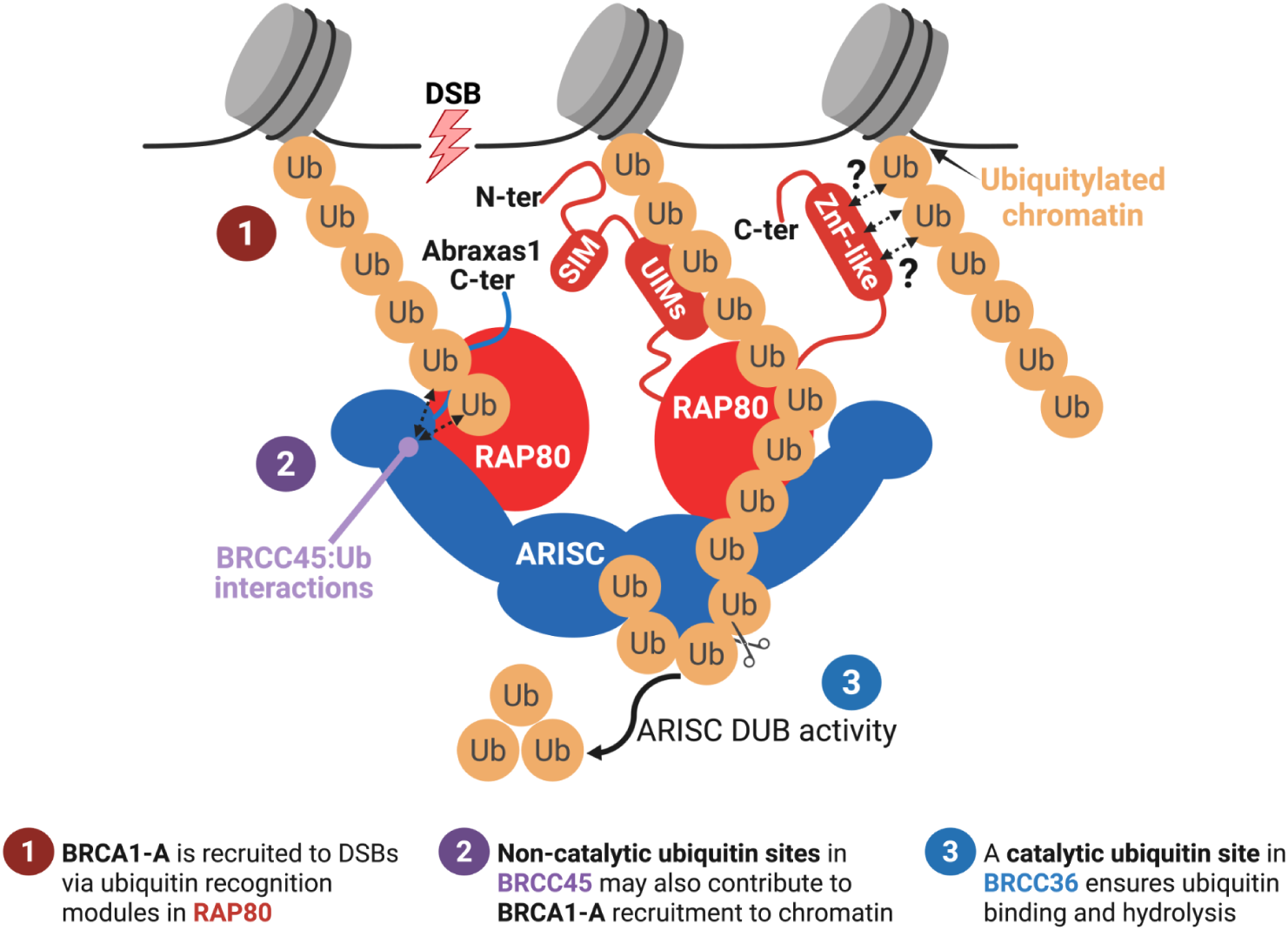
“Read-and-Erase” model for BRCA1-A recognition and cleavage of ubiquitin chains on chromatin. Recruitment of the BRCA1-A complex to DNA damaged sites requires recognition of K63-linked ubiquitin chains on chromatin by different RAP80 regions (**1**) and low-affinity, non-catalytic ubiquitin sites on the BRCC45 arms (**2**). Substrate cleavage is dependent on ubiquitin chain interaction with the catalytic ubiquitin site in BRCC36 and Abraxas1* from the adjacent hetero-tetramer (**3**). This mechanism promotes BRCA1-A recruitment to chromatin after DNA damage, while allowing continuous ubiquitin chain cleavage by the ARISC catalytic core.

Ubiquitin chains adopt a diverse range of conformations, with K63-linked polymers forming “open” and extended structures that lack intra-chain ubiquitin interactions^11^. In the ARISC and ARISC–RAP80 cryo-EM structures described herein, cyclic K63-linked chains adopt a previously-undescribed conformation characterised by extensive contact surfaces between its ubiquitin moieties (**Fig. 2a,c** and **Fig. 5a**). This differs considerably from available crystal structures of the isolated RAP80 UIMs, TAB2 ZnF, and RNF168 UDM1/2 bound to K63-Ub2^19–21^. These data suggest that the inherent flexibility of K63-linked ubiquitin chains may allow macromolecular complexes containing multiple ubiquitin-binding domains to recognise and remodel K63-linked chains, thereby facilitating different cell signalling outcomes. In addition, our work raises questions on whether cyclical polyubiquitin chains play a functionally relevant role in the DDR. Homotypical K63-linked and hybrid SUMO-ubiquitin chains have also been reported at DSBs^54,55^, and RAP80 has been shown to interact with branched K11/K63-linked polymers *in vitro*^56^. However, cyclical K63-linked chains have yet to be identified in the same cellular context. While we acknowledge that the cyclical K63-linked ubiquitin chains reported in this work may be an artifact of our *in vitro* system, we do not exclude the possibility that such chains may exist at the cellular level. It may be possible that, in cells, cyclical chains represent a short-lived ubiquitin pool “poised” to become available to the ubiquitylation machinery only upon cleavage by specific DUBs, which would then turn these chains into linear polymers. Elucidating the functional relevance of cyclical ubiquitin chains in the DDR is an important endeavour for future studies.

ARISC and ARISC–RAP80 complexes stably interact with polyubiquitin chains via a three-subunit interface, comprising BRCC36, Abraxas1* and, to a lesser extent, BRCC45* (**Fig. 2a**, **Fig. 5a**, **Extended Data Fig. 5c,g**, and **Supplementary Fig. 11c,f**). BRCC36 shares structural similarities with AMSH-LP, and both enzymes use their Ins-1 helix and Ins-1 loop to engage with ubiquitin chains in their active sites (**Fig. 3a**). However, BRCC36 interacts with Ub (P1’) via its S-loop and loop3 regions and does not use Ins-2 like AMSH-LP (**Fig. 3a**). Consistent with these observations, our cell-based studies, together with the lack of sequence conservation for Ins-2 across BRCC36 species and the absence of clear density for this region in our structural models (**Fig. 4d,e**, **Supplementary Fig. 5c**, and **Supplementary Fig. 11c**)^41^, confirmed a non-prominent role for BRCC36 Ins-2 in mediating ubiquitin binding. Beyond the BRCC36 active site, the pseudo-DUB Abraxas1 plays a pivotal role in mediating ARISC and ARISC–RAP80 engagement with polyubiquitin chains and in regulating DUB activity (**Fig. 3f,h,i** and **Fig. 4c,f**). The mode of Abraxas1*-mediated interaction with ubiquitin chains is unique to ARISC and ARISC–RAP80 assemblies, as the corresponding β2-loop3-β3 regions in CSN6 and RPN8 are not as extended as in Abraxas1*^53,57^. Our analyses therefore highlight how JAMM/MPN DUBs have evolved to use different structural elements, and interact with protein partners to maximise substrate binding and enzymatic activity even when recognising the same ubiquitin linkage type. Curiously, ubiquitin chains interact with only one BRCC36 protomer in our structures while the corresponding BRCC36* interface appears to be unoccupied (**Fig. 2a** and **Fig. 5a,b**). Our analyses indicate that structural re-arrangements or re-positioning of the ubiquitin molecules within ARISC may allow both BRCC36 active sites to concomitantly engage with their ubiquitin substrates (**Supplementary Fig. 5e**). Alternatively, it is possible that ARISC–RAP80 binds to ubiquitin chains on both of its active sites simultaneously, but only cleaves one ubiquitin substrate at a time. Further studies are required to understand the precise contribution of each BRCC36 protomer to ARISC–RAP80 driven enzymatic activity.

Ubiquitin binding domains (UBDs) are small protein modules that non-covalently bind to mono or polyubiquitin moieties^1,58^. In the ARISC–RAP80 complex, known RAP80 UBDs, including its N-terminus and UIMs, as well as the SUMO-UIMs linker, AIR, and ZnF-like domain, are positioned in proximity to K63-linked ubiquitin molecules distal from the BRCC36 catalytic site (**Fig. 8**). We have previously identified several residues in the RAP80 N-terminus, UIMs and AIR regions, alongside BRCC36 K65 and K204, to be ubiquitylated in the inactive ARISC(E33A)–RAP80 complex, and proposed a model whereby BRCC36 functions to relieve ARISC–RAP80 auto-inhibition and promote recognition of DNA damaged sites^34,40^. The RAP80:ubiquitin interactions described here are consistent with our previous findings, and further highlight the importance of concerted ubiquitin binding and hydrolysis actions for sustained ARISC–RAP80 recruitment to DSBs.

Our biochemical analyses indicate that ARISC and ARISC–RAP80 complexes are endo-DUBs, although they preferentially initiate ubiquitin hydrolysis from the distal end of the chain (**Fig. 6**). This mechanism is reminiscent of MINDY1 and MINDY2, which endo- or exo-DUB activities are dependent on ubiquitin chain length^17^, but differs from RPN11, a JAMM/MPN DUB that has evolved to perform *en bloc* removal of ubiquitin chains from proteasome-associated substrates^59^. More recently, USP53, another K63-specific DUB, was also shown to prefer en bloc removal of polyubiquitin substrates over endo-DUB activity^22^. As such, the catalytic mechanism reported for ARISC and ARISC–RAP80 complexes adds to the diversity of DUB activities within and outside the JAMM/MPN family.

Chromatin-associated proteins and protein complexes interact with nucleosomes via direct binding to the histone octamer and/or nucleosomal DNA, or by recognising specific post-translational marks^9,60^. Here we propose that, in addition to RAP80, ARISC may utilise its non-catalytic ubiquitin sites in BRCC45 to interact with mono and/or polyubiquityled chromatin (**Fig. 8**). BRCA1-BARD1 interacts directly with nucleosomes and recognizes monoubiquitylated histone H2A on lysine 15 (H2A-K15Ub) on damaged chromatin^46,61–64^. It is possible that ARISC–RAP80 may also directly bind to the nucleosome surface or engage with BRCA1-BARD1 to mediate multivalent interactions with several modified nucleosomes. How these modular complexes navigate chromatin engagement to direct DNA repair represents an important question for future research that will shed light on links between damage recognition and genome stability.

## Methods

### Bacterial and insect cell cultures

Bacterial cells were grown in Luria-Bertani (LB) broth (Fisher Chemical) or Terrific Broth (TB; Fisher Chemical) media supplemented with appropriate antibiotics [100 µg/mL ampicillin (Merk Life Science), 34 µg/mL chloramphenicol (Amresco), 10 µg/mL gentamycin (VWR), 50 µg/mL kanamycin (Thermo Fisher Scientific), 10 µg/mL tetracycline (Sigma)], and incubated at 37°C and 18°C as per requirements. For preparation of bacmids DNA, x-gal (Thermo Fisher Scientific) and isopropyl-β-D-1-thiogalactopyranoside (IPTG; Fluorochem) were also added to bacteria cultures at final concentrations of 15 µg/mL and 40 µg/mL, respectively. *Spodoptera frugiperda 9* (*Sf9*) insect cells (Thermo Fisher Scientific) were grown in SF 900 II SFM media (Thermo Fisher Scientific) supplemented with 1x antibiotic-antimycotic mix (Thermo Fisher Scientific), and maintained at 27°C.

### Mammalian cell culture

HT-29 and HEK293T cell lines were purchased from ATCC and cultured in 1x Dulbecco’s modified Eagle’s medium (Gibco) supplemented with 10% bovine calf serum (Corning) and 1% Pen Strep (Gibco) with 5% CO_2_ at 37°C.

### Generation of plasmid constructs

#### Cloning and expression of protein complexes in insect cells

To generate constructs for insect cells expression, genes for the four-subunits human full-length (FL) ARISC (BRCC36, Abraxas1, BRCC45, and MERIT40) and BRISC (BRCC36, Abraxas2, BRCC45, and MERIT40) wild-type (WT) complexes were cloned into pFL (BRCC36-Abraxas1 or BRCC36-Abraxas2) and pUCDM (BRCC45-MERIT40) vectors, and assembled into single multigene cassettes via Cre-mediated recombination^65^. A 6xHis and double (d)StrepII purification tags, followed by either a Tobacco Etch Virus (TEV) or PreScission protease sites, were engineered at the N-termini of BRCC45 and Abraxas1 or Abraxas2, respectively. FL ARISC(E33A) (BRCC36 E33A) and ARISCΔC(E33A) (BRCC36 E33A, Abraxas1ΔC) complexes as well as all ARISC and ARISC(E33A) variants, bearing mutations or deletions in BRCC36, Abraxas1 or BRCC45, were cloned and assembled into multigene constructs as described above. ARISCΔN (MERIT40ΔN), ARISCΔC (Abraxas1ΔC) and ARISCΔNΔC (Abraxas1ΔC, MERIT40ΔN) WT complexes were cloned and assembled as described above, with the exception that these expression constructs only bear a 6xHis tag and TEV protease site at the N-terminus of BRCC45. The DNA sequences for human FL RAP80 WT, RAP80 ΔUIMs, RAP80 ΔZnF, and RAP80 AIR were cloned into pFastBac-HTB using standard molecular biology techniques. All RAP80 constructs are characterised by a 6xHis purification tag and TEV protease site at their N-termini. A list of expression constructs and primers used in this study is summarised in **Supplementary Tables 2** and **3**.

Bacmid DNA was generated in DH10 MultiBac*^Turbo^* cells (Geneva Biotech) following manufacturer’s protocol, and virus amplification in *Sf9* cells (Thermo Fisher Scientific) was performed using standard procedures and as previously described^35,40^. For recombinant expression of ARISC and BRISC variants, *Sf9* cells were infected with baculoviruses encoding each construct. To express ARISC–RAP80 and ARISC(E33A)–RAP80 variants, bearing mutations or deletions in BRCC36, Abraxas1 or BRCC45, as well as ARISC–RAP80 ΔUIMs, ARISC–RAP80 ΔZnF, and ARISC–RAP80 AIR complexes, *Sf9* cells were co-infected with baculoviruses encoding ARISC or ARISC(E33A) variants and each RAP80 variant in a 1:1 ratio. Following 48 h after infection, cells were harvested by centrifugation at 500 g for 15 min and protein purification was carried out as outlined below.

#### Cloning and expression of proteins in bacteria

Expression constructs for UBA1, Ubc13, Mms2, and ubiquitin WT were a gift from Professor Frank Sicheri. Constructs expressing pOPINB-AMSH* (Addgene plasmid number: 66712) and pOPINB-OTULIN (Addgene plasmid number: 61464) were a gift from Professor David Komander^15,43^. To allow site-specific labelling of ubiquitin WT cloned into pProEx-HTB with Alexa-Fluor 488 C_5_ maleimide dye (Thermo Fisher Scientific), a cysteine residue was introduced between the N-terminal 6xHis tag/TEV cleavage site and the first methionine of ubiquitin by site-directed mutagenesis. The resulting construct, referred to as Cys0-ubiquitin, was used as template for a second round of site-directed mutagenesis where lysine 63 was mutated to arginine. This allowed the generation of a ubiquitin variant blocked at the distal end, here referred to as Cys0-ubiquitin K63R. In parallel, the gene encoding ubiquitin 1-74 (referred to as ubiquitin ΔGG) was cloned into pET-3a using standard techniques. To allow site-specific labelling of ubiquitin ΔGG as indicated above and generate a variant blocked at the proximal end, arginine 74 was mutated to cysteine. This approach resulted in an additional expression construct, here referred to as ubiquitin R74C ΔGG. To ensure correct assembly of linear ubiquitin chains, ubiquitin K63R was cloned into pET-3a with an N-terminal 8xHis purification tag and TEV protease site. A sequence containing tryptophan, glycine, and cysteine residues (referred to as “WGC”) was also introduced after the TEV cleavage site to allow for precise protein quantification. All mutations were confirmed by DNA sequencing. A list of expression constructs and primers used in this study is summarised in **Supplementary Tables 2** and **3**.

UBA1, Mms2, Cys0-ubiquitin K63R, and ubiquitin K63R were expressed with an N-terminal 6xHis-glutathione S-transferase (GST) (UBA1), maltose binding protein (MBP)-6xHis (Mms2), 6xHis (Cys0-ubiquitin K63R) or 8xHis (ubiquitin K63R) purification tags, each followed by a TEV protease site, in *Escherichia coli* (*E. coli*) BL21 (DE3) RIL cells (Thermo Fisher Scientific). Ubc13, AMSH*, and OTULIN were expressed with an N-terminal GST tag and thrombin site (Ubc13) or with an N-terminal 6xHis tag and PreScission protease site (AMSH* and OTULIN) in the same cell line. Ubiquitin WT and R74C ΔGG variants were expressed as untagged construct in *E. coli* BL21 (DE3) RIL and Rosetta (Novagen) cells, respectively. Cells were grown at 37°C in LB or TB medium to an OD_600_ between 0.5-1.5, then induced with 0.5 mM-1 mM IPTG and grown overnight (O/N) at 18°C. Cells were harvested by centrifugation at 6,000 g for 20 min, and protein purification was carried out as outlined below.

### Generation of knock out cell lines using CRISPR-Cas9

CRISPR-based gene knockout was carried out using Streptococcus pyogenes Cas9 (SpCas9) system. HT-29 or HEK293T cells stably expressing SpCas9 were generated by lentiviral transduction of lenti-SpCas9 hygro (Addgene plasmid number: 104995) followed by hygromycin (500 µg/mL) selection. sgRNA guides for Abraxas1 (GCGGCGGTAGCATGGA), BRCC36 (TCTAGTTGAACGATGATACA) or BRCC45 (ctctgttagttttgaggaca) were cloned into lentiGuide-Puro (Addgene, plasmid number: 52963) sgRNA expression vector digested with BsmBI-v2 (New England Biolabs). Plasmids (2.7 µg) carrying individual sgRNAs were packaged into lentivirus particles by co-transfection with psPAX2 (2 µg; Addgene plasmid number: 12260) and pMD2.G (1.3 µg; Addgene plasmid number: 12259) packaging plasmids into HEK293T cells using lipofectamine 2000 (Invitrogen). Lentivirus supernatant was collected 48 h post-transfection and filtered through a 0.45 µm filter (Millipore). Cas9 expressing HT-29 or HEK293T cells were transduced using polybrene (8 µg/mL; Sigma-Aldrich) by spin infection at 2,000 rpm for 60 min. Transduced cells were selected by puromycin (4 µg/mL) and knockout efficiencies were determined by Western Blotting.

### Site-directed mutagenesis and cell lines reconstitution

Abraxas1 and BRCC36 mutants were generated by site-directed mutagenesis of pOZ-N-FH Abraxas (Addgene plasmid number: 27495) and pOZ-N-FH BRCC36 (Addgene plasmid number: 27496)^34^,respectively using Q5 Site-Directed Mutagenesis kit (New England Biolabs; catalogue number: E0554S) following manufacturer’s protocol. BRCC45 mutants were generated by site-directed mutagenesis of FLAG tagged pCMV6-BRCC45 plasmid (RC220494, OriGene Technologies, Inc.).For BRCC36 reconstitution and BRCC45 transient overexpression, sgRNA-resistant plasmids were generated via site-directed mutagenesis and were subsequently used for additional mutagenesis. Phoenix cells were used to generate retrovirus particles following co-transfection of the pOZ-expression plasmids and pCL-Ampho packaging vector with 2:1 ratio using lipofectamine 2000 (Invitrogen). Virus supernatant was collected 48 h post-transfection and filtered through a 0.45 µm filter. Abraxas1 or BRCC36 KO HT-29 cells were transduced with virus particles carrying respective WT or mutant pOZ-expression plasmids using polybrene (8 µg/mL; Sigma-Aldrich) by spin infection at 2,000 rpm for 90 min. Transduced cells were selected using anti-IL2-receptor antibody 48 h post-transduction and overexpression was determined by Western Blotting.

### Protein purification

#### Purification of ARISC, ARISC–RAP80, and BRISC complexes

Purifications of 6xHis tagged ARISCΔN, ARISCΔC, and ARISCΔNΔC WT complexes were carried out as previously described^35,40^, with the exception that proteins were resolved on a Superose 6 10/300 Increase column (Cytiva) pre-equilibrated in 25 mM HEPES-NaOH pH 7.3, 150 mM NaCl, 10% (v/v) glycerol, and 1 mM tris(2-carboxyethyl)phosphine (TCEP). Eluted peaks were analysed by SDS-PAGE, and fractions containing >95% pure ARISC variants were combined, concentrated to 5-10 mg/mL, snap-frozen in liquid nitrogen, and stored at -80°C.

Purifications of 6xHis and (d)StrepII tagged ARISC and ARISC–RAP80 variants as well as BRISC were performed as previously described^35,38,40,41^, with minor modifications. Briefly, eluted fractions from nickel-affinity chromatography containing ARISC, ARISC–RAP80, and BRISC complexes were pooled, and 0.3 mg of 6xHis-TEV protease was added prior to O/N dialysis against 4 L of dialysis buffer [50 mM Tris-HCl pH 7.5, 150 mM NaCl, 5% (v/v) glycerol, and 1 mM TCEP] at 4°C. The dialysed samples were subsequently incubated with 2 mL Strep-Tactin Sepharose resin (IBA Lifesciences GmbH), pre-equilibrated in binding buffer [50 mM Tris-HCl pH 7.5, 300 mM NaCl, 5% (v/v) glycerol, and 1 mM TCEP], for 1 h at 4°C in rotation. After washing the resins four times with 50 mL of the same buffer, ARISC, ARISC–RAP80, and BRISC complexes were eluted by subsequent washes (1 mL each) with elution buffer (binding buffer supplemented with 5 mM desthiobiotin). Fractions containing ARISC and ARISC–RAP80 variants as well as BRISC were pooled, concentrated and resolved on a Superose 6 10/300 Increase column (Cytiva) pre-equilibrated in 25 mM HEPES-NaOH pH 7.3, 150 mM NaCl, 10% (v/v) glycerol, and 1 mM TCEP. ARISC(E33A)–RAP80 and ARISCΔC(E33A) complexes used for structural studies were purified on the same size-exclusion chromatography column, with the exception that 25 mM HEPES-NaOH pH 7.3, 150 mM NaCl, and 1 mM TCEP was used as storage buffer. Eluted peaks were analysed by SDS-PAGE, and fractions containing >95% pure ARISC, ARISC–RAP80, and BRISC proteins were combined, concentrated to 2-10 mg/mL, snap-frozen in liquid nitrogen, and stored at -80°C. Purifications of all ARISC WT and ARISC(E33A)–RAP80 variants, bearing mutations or deletions in BRCC36, Abraxas1 or BRCC45, were carried out in an identical manner.

#### Purification of AMSH* and OTULIN

Purifications of 6xHis tagged AMSH* and OTULIN were carried out as previously described^35,40^, with the exception that 0.3 mg/mL lysozyme was added to the lysis buffer. Following nickel-affinity chromatography, the recovered proteins were pooled and 0.3 mg of 6xHis-PreScission protease added prior to O/N dialysis against 4 L of dialysis buffer [50 mM Tris-HCl pH 7.6, 150 mM NaCl, 20 mM imidazole, 5% (v/v) glycerol, 0.075% (v/v) β-mercaptoethanol, and 1 mM benzamidine] at 4°C. After removal of the 6xHis-PreScission protease and uncleaved 6xHis tagged AMSH* or OTULIN by subtraction using 1 mL HisPur^TM^ Ni-NTA resin (Thermo Fisher Scientific), the cleaved proteins were concentrated and resolved on a Superdex 75 10/300 GL column (GE Healthcare) pre-equilibrated in 50 mM HEPES-NaOH pH 7.5, 150 mM NaCl, and 1 mM TCEP. Eluted peaks were analysed by SDS-PAGE, and fractions containing >95% pure AMSH* or OTULIN were combined, concentrated to 2 mg/mL, snap-frozen in liquid nitrogen, and stored at -80°C.

#### Purification of UBA1, Ubc13, Mms2, and ubiquitin variants

Purifications of 6xHis-GST tagged UBA1, GST-tagged Ubc13, and MBP-6xHis tagged Mms2 were performed as previously described^64^. Purifications of untagged ubiquitin WT and R74C ΔGG, and of 6xHis tagged Cys0-ubiquitin K63R and 8xHis tagged ubiquitin K63R, were carried out as described elsewhere^18,45,66^.

#### Preparation of wild type and distally blocked chains

Wild type (WT) lysine 63 (K63)-linked ubiquitin chains were assembled as previously described^67^, with minor modifications. Briefly, purified UBA1 (at 1 μM), Ubc13 and Mms2 (at 8 μM), and ubiquitin WT (at 1 mM) were incubated for 6 h at 37°C in a buffer containing 40 mM Tris-HCl pH 8.5, 10 mM MgCl_2_, 10 mM adenosine triphosphate (ATP), and 0.5 mM dithiothreitol (DTT). The reaction was stopped by the addition of 10 mM DTT, centrifuged at 15,000 g for 5 min to remove any precipitated proteins, and then diluted ten-fold in buffer A (50 mM sodium acetate pH 4.5). Ubiquitin chains were separated using a Resource S column 6 mL (Cytiva) over a linear gradient from 0%-50% buffer B (buffer A supplemented with 1 M NaCl) in 58 CV. A final concentration of 100 mM HEPES-NaOH pH 7.5 was added to all eluted fractions in order to bring the pH up to ∼ 7.0, and eluted peaks were analysed by SDS-PAGE. Fractions containing >95% pure ubiquitin chains of each length were combined, concentrated to 1-3 mg/mL, snap-frozen in liquid nitrogen, and stored at -80°C.

Distally blocked K63-linked ubiquitin chains were assembled as previously described^45^, with minor modifications. Briefly, purified UBA1 (at 0.6 μM), Ubc13 and Mms2 (at 8 μM), ubiquitin WT (at 1 mM), and 8xHis-TEV ubiquitin K63R (at 200 μM) were incubated for O/N at 37°C in a buffer containing 50 mM Tris-HCl pH 7.6, 5 mM MgCl_2_, 2.5 mM ATP, 10 mM creatine phosphate, and 0.6 U/mL creatine phosphokinase. The reaction was stopped and centrifuged as described above, and then diluted ten-fold in low salt buffer [50 mM Tris-HCl pH 7.5, 300 mM NaCl, 20 mM Imidazole, 5% (v/v) glycerol, and 1 mM DTT]. The sample was then incubated with 2.5 mL HisPur^TM^ Ni-NTA resin (Thermo Fisher Scientific), pre-equilibrated in low salt buffer, for 1h at 4°C in rotation. After washing the resin twice with 10 CV low salt buffer, 8xHis tagged ubiquitin chains were eluted by subsequent washes (6 mL each) with elution buffer (low salt buffer containing 200 mM Imidazole). Fractions containing purified 8xHis ubiquitin chains were pooled, incubated with 0.3 mg of 6xHis-TEV protease, and dialysed O/N against 4 L of dialysis buffer (low salt buffer containing 150 mM NaCl). Following removal of the TEV protease by subtraction using 1.5 mL HisPur^TM^ Ni-NTA resin (Thermo Fisher Scientific), the cleaved distally blocked ubiquitin chains were diluted ten-fold in buffer A (50 mM sodium acetate pH 4.5). Ubiquitin chains were then purified as described above, with the exception that a Resource S column 1 mL (Cytiva) was used and chains were separated over a 100 CV linear gradient. A final concentration of 100 mM HEPES-NaOH pH 7.5 was added to all eluted fractions in order to bring the pH up to ∼ 7.0, and eluted peaks were analysed by SDS-PAGE. Fractions containing >95% pure ubiquitin chains of each length were combined, concentrated to 1 mg/mL, snap-frozen in liquid nitrogen, and stored at -80°C.

#### Preparation of proximally and distally blocked chains

Synthesis and purification of proximally and distally blocked K63-linked tetraUb (Ub4) chains was carried out as described for the WT ubiquitin chain variants, with the exception that purified triUb (Ub3) chain was reacted in a 1:3 molar ratio with ubiquitin R74C ΔGG (to assemble proximally blocked chains) or Cys0-ubiquitin K63R (to assemble distally blocked chains) and each reaction mixture was incubated for 3 h at 37°C. Following purification, fractions containing >95% pure proximally or distally blocked Ub4 variants were combined, concentrated to 1.5 mg/mL, snap-frozen in liquid nitrogen, and stored at -80°C.

#### Labelling of proximally and distally blocked chains

Purified proximally and distally blocked tetraUb (Ub4) chains were each mixed with Alexa-Fluor 488 C_5_ maleimide dye (Thermo Fisher Scientific) and TCEP in a 1:10:20 molar ratio in a buffer containing 20 mM HEPES-NaOH pH 7.5 and 100 mM NaCl. The reactions were then incubated at room temperature for O/N in rotation and protected from light. Labelling reactions were subsequently stopped by the addition of 5 mM DTT and 2 mM L-Cysteine, and further incubated at room temperature for 30 min in rotation. Each sample was then passed through a 0.5 mL Zeba^TM^ spin desalting column (Thermo Fisher Scientific), pre-equilibrated in 25 mM HEPES-NaOH pH 7.5, 150 mM NaCl, and 1 mM TCEP, to remove excess dye, concentrated to 0.3-0.8 mg/mL, snap-frozen in liquid nitrogen, and stored at - 80°C.

### AlphaFold 3D structure prediction

A model for the human BRCC36 protein was retrieved from the AlphaFold Protein Structure Database (https://alphafold.ebi.ac.uk/) under accession code AF-P46736-F1-v4. To generate models for BRCC45:ubiquitin and Abraxas1:ubiquitin sub-complexes, the corresponding protein sequences were used as inputs in AlphaFold 3^68^ assuming a 1:1 stoichiometry. Structural predictions were run using the default settings on the AlphaFold 3 webserver, asking for five models as outputs, and manually inspected in UCSF Chimera X v1.6.1^69,70^.

### Grids preparation and data collection

Freshly purified ARISCΔC(E33A) and ARISC(E33A)–RAP80 (at 0.8 mg/mL and 0.6 mg/mL, respectively) were mixed with 1.5-fold molar excess of K63-linked tetraUb (Ub4) and/or heptaUb (Ub7) chains, and incubated in the presence of 0.025% (v/v) glutaraldehyde (Sigma-Aldrich) at room temperature for 4 min. Cross-linking reactions were then quenched by the addition of 100 mM Tris-HCl pH 7.5 prior to cryo-EM grids preparation. UltrAuFoil R1.2/1.3 300-mesh grids (Quantifoil Micro Tools GmbH) were glow-discharged for 1 min at 12 mA and 0.38 mBar pressure using a PELCO easiGlow system (Ted Pella). Cryo-EM grids were prepared by applying 3 μL of each cross-linked sample onto the glow-discharged UltrAuFoil grids, followed by immediate blotting (blot force = 1 N, blot time = 6 s, waiting time = 27 s) and plunge-freezing in liquid ethane cooled by liquid nitrogen, using a FEI Vitrobot IV (Thermo Fisher Scientific) at 100% relative humidity and with a chamber temperature set at 4°C.

All datasets (four for ARISCΔC(E33A):K63-Ub7, two for ARISCΔC(E33A):K63-Ub4, and one for ARISC(E33A)–RAP80:K63-Ub7) were collected on a FEI Titan KRIOS transmission electron microscope (Thermo Fisher Scientific) operating in counting mode at 300 keV, using a magnification of 165,000x and a pixel size of 0.74 Å. For ARISCΔC(E33A):K63-Ub7 complex, a total of 55,093 movies were recorded using the EPU automated acquisition software (v3.8) on a FEI Falcon 4i direct electron detector^71^ with an energy filter of 10 eV. A dose per physical pixel/s between 6.65-7.42 was used for each exposure, resulting in a total electron dose of 45.5 e^-^/Å^2^. Frames were then grouped into 43 fractions, resulting in an electron dose between 1.06-1.17 e^-^/Å^2^ per frame/fraction. Four exposures per hole were collected with an exposure time around 3.36-3.75 s each, defocus values ranging from -0.9 µm to -3.0 µm, and an aberration free image shift (AFIS) range of 10 µm. For ARISCΔC(E33A):K63-Ub4 complex, a total of 58,037 movies were recorded as outlined above. A dose per physical pixel/s between 7.35-7.39 was used for each exposure, resulting in a total electron dose around 45.6-45.9 e^-^/Å^2^. Frames were then grouped into 39 fractions, resulting in an electron dose of 1.17 e^-^/Å^2^ per frame/fraction. Four exposures per hole were collected as described above, using an exposure time of 3.4 s. For ARISC(E33A)–RAP80:K63-Ub7 complex, a total of 10,649 movies were recorded as indicated above. A dose per physical pixel/s of 7.33 was used for each exposure, resulting in a total electron dose of 45.5 e^-^/Å^2^. Frames were then grouped into 39 fractions, resulting in an electron dose of 1.17 e^-^/Å^2^ per frame/fraction. Four exposures per hole were collected as described above, using an exposure time of 3.4 s.

Detailed information on data collection, refinement, and validation statistics is shown in Supplementary Table 1.

### Data processing

Schematics of the data processing pipelines are shown in **Supplementary Fig. 6b** (for ARISC(E33A)–RAP80:K63-Ub7), **Supplementary Fig. 12b** (for ARISCΔC(E33A):K63-Ub7) and **Supplementary Fig. 13b** (for ARISCΔC(E33A):K63-Ub4), and further details on the reported maps and models are available in **Supplementary Table 1**. Image processing was carried out using a combination of RELION v4.0^72^ and cryoSPARC v4.5.3^73^. Drift-corrected averages of each movie were created using cryoSPARC patch motion correction^73,74^, and real-time contrast transfer function (CTF) parameters of each determined using cryoSPARC patch CTF estimation^73^. Motion correction and CTF estimation were carried out using cryoSPARC Live^TM^ (https://cryosparc.com/live).

For ARISCΔC(E33A):K63-Ub7, 1,049 particles were manually picked from dataset I and used to train crYOLO v1.6.1^75^. This trained model was used for picking on all datasets, resulting in 2,206,570 total particles (231,627 particles from dataset I; 869,099 particles from dataset II; 637,740 particles from dataset III; 468,104 particles from dataset IV). Particles from each dataset were imported in cryoSPARC, extracted using a box size of 400 x 400 pixels and a binning factor of four, and independently subjected to iterative rounds of reference-free 2D classification with a mask diameter of 240 Å. After visual inspection, high quality 2D classes obtained from each dataset (97,221 particles from dataset I; 281,336 particles from dataset II; 173,070 particles from dataset III; 237,597 particles from dataset IV) were combined, re-extracted using a box size of 400 x 400 pixels without binning, and subjected to further rounds of reference-free 2D classification as described above. Following 2D classification, a total of 402,933 particles were retained and used to generate seven initial 3D volumes. These models were subsequently subjected to 3D classification using heterogeneous refinement with C1 symmetry, which yielded two well-resolved maps (183,040 total particles) representing ARISCΔC bound to ubiquitin chains. These particles and the best-resolved model were then used for homogeneous and non-uniform 3D refinements with C1 symmetry, generating an initial map with a global resolution of 3.17 Å. To eliminate alignment artefacts, a mask comprising the densities belonging to ARISCΔC and ubiquitin chains was created using UCSF ChimeraX v1.6.1^69,70^. This mask, together with the particles and map obtained from non-uniform 3D refinement, were imported in RELION and subjected to focused alignment-free 3D classification with C1 symmetry and four classes as outputs. One of the classes obtained from this 3D classification step was characterised by clear densities surrounding ARISCΔC and ubiquitin moieties, and comprised a total of 100,036 particles. These particles and the best-resolved map were then imported in cryoSPARC and subjected to non-uniform 3D refinement, yielding a final map with a global resolution of 3.20 Å. To improve the density of the map around the BRCC45 arms and ubiquitin moieties, focused masks comprising these regions were created using UCSF ChimeraX v1.6.1^69,70^. In parallel, masks comprising the remaining portions of the map outside of these regions were also created in the same manner. These masks, together with the particles and map obtained from non-uniform 3D refinement, were initially used for particle subtraction. The resulting particles, focused masks, and map obtained from non-uniform 3D refinement were then used for local refinements with C1 symmetry. This generated improved maps with final resolutions of 3.44 Å (Right Arm), 3.57 Å (Left Arm), 3.7 Å (Ub (P2) & Ub (P3)), and 3.41 Å (Ub (P1’)), which were combined with the map obtained from non-uniform 3D refinement to assemble a composite map for the ARISCΔC(E33A):K63-Ub7 complex.

For ARISCΔC(E33A):K63-Ub4, the same crYOLO model described above was used for picking on all datasets. The resulting 2,194,750 total particles (1,383,733 particles from dataset I; 811,017 particles from dataset II) were then imported in cryoSPARC, extracted and independently subjected to iterative rounds of reference-free 2D classification as described for ARISCΔC(E33A):K63-Ub7. After visual inspection, high quality 2D classes obtained from each dataset (538,732 particles from dataset I; 295,266 particles from datasets II) were re-extracted, combined, and subjected to further rounds of reference-free 2D classification as described above. Following 2D classification, a total of 399,099 particles were retained and used to generate six initial 3D volumes. To separate the density corresponding to ARISCΔC in complex with ubiquitin chains from a low-occupancy ARISCΔC dimer, the best-resolved initial model (100,118 particles) was subsequently subjected to 3D classification using heterogeneous refinement with C1 symmetry. This yielded a well-resolved map (54,452 particles) that was used as a template for homogeneous and non-uniform 3D refinements with C1 symmetry, generating a final map with a global resolution of 3.21 Å. To improve the density of the map around the BRCC45 arms and ubiquitin moieties, focused masks comprising these regions were created using UCSF ChimeraX v1.6.1^69,70^. In parallel, masks comprising the remaining portions of the map outside of these regions were also created in the same manner. These masks, together with the particles and map obtained from non-uniform 3D refinement, were initially used for particle subtraction. The resulting particles, focused masks, and map obtained from non-uniform 3D refinement were then used for local refinements with C1 symmetry. This generated improved maps with final resolutions of 3.88 Å (Right Arm), 3.76 Å (Left Arm), and 3.53 Å (Ub (P1’)), which were combined with the map obtained from non-uniform 3D refinement to assemble a composite map for the ARISCΔC(E33A):K63-Ub4 complex.

For ARISC(E33A)–RAP80:K63-Ub7, 1,000 particles were manually picked and used to train crYOLO v1.6.1^75^. This trained model was used for picking on all movies, resulting in 789,840 total particles. Particles were then imported in cryoSPARC, extracted using a box size of 460 x 460 pixels and a binning factor of five, and subsequently subjected to iterative rounds of reference-free 2D classification with a mask diameter of 280 Å. After visual inspection, high quality 2D classes (532,328 particles) were re-extracted using a box size of 460 x 460 pixels without binning and subjected to further rounds of reference-free 2D classification as described above. Following 2D classification, a total of 349,497 particles were retained and used to generate three initial 3D volumes. The best-resolved model (214,650 particles) was subsequently used as a template for homogeneous and non-uniform 3D refinements with C1 symmetry, generating a final map with a global resolution of 2.92 Å. To improve the densities of the map around the BRCC45-MERIT40 arms and ubiquitin moieties, focused masks comprising these regions were created using UCSF ChimeraX v1.6.1^69,70^. In parallel, masks comprising the remaining portions of the map outside of these regions were also created in the same manner. These masks, together with the particles and map obtained from non-uniform 3D refinement, were initially used for particle subtraction. The resulting particles, focused masks, and map obtained from non-uniform 3D refinement were then used for local refinements with C1 symmetry. This generated improved maps with final resolutions of 4.09 Å (Right Arm), 4.02 Å (Left Arm), 3.17 Å (Ub (P2) & Ub (P3)), 3.16 Å (Ub (P1’)), 3.99 Å (Non-catalytic Ub, Right Arm), and 3.91 Å (Non-catalytic Ub, Left Arm), which were combined with the map obtained from non-uniform 3D refinement to assemble a composite map for the ARISC(E33A)–RAP80:K63-Ub7 complex.

The final resolutions of all maps were determined using the gold-standard Fourier shell correlation criterion (FSC = 0.143); local resolutions were determined using the local resolution implementation in cryoSPARC v4.5.3^73^. DeepEMhancer^76^ was employed using default settings (i.e. “tightTarget” model) for local sharpening.

### Model building and refinement

Atomic models for ARISC(E33A)–RAP80:K63-Ub7, ARISCΔC(E33A):K63-Ub7, and ARISCΔC(E33A):K63-Ub4 complexes were built by using high-resolution cryo-EM maps. A preliminary model of the human ARISC–RAP80 AIR complex^40^ was rigid-body fitted into the ARISC(E33A)–RAP80:K63-Ub7 composite map using UCSF Chimera X v1.6.1^69,70^. The same model with BRCC45 UEV-C, MERIT40, and RAP80 AIR removed was also rigid-body fitted into the ARISCΔC(E33A):K63-Ub7 and ARISCΔC(E33A):K63-Ub4 composite maps in the same manner. The crystal structure of human ubiquitin (PDBid = 1UBQ)^77^ was used as a starting model for the K63-linked ubiquitin chains and rigid-body fitted four (for ARISC(E33A)–RAP80:K63-Ub7 and ARISCΔC(E33A):K63-Ub7) or two (for ARISCΔC(E33A):K63-Ub4) times into all composite maps as outlined above. The resulting models were then manually inspected and rebuilt using Coot v0.9.8.1^78^, and iterative rounds of real-space refinement were performed in Coot v0.9.8.1 and PHENIX v1.17.1^79^ using default parameters, and secondary structure and bonds geometry (to ensure formation of the K63-linked isopeptide bond between the different ubiquitin moieties) restraints. The Abraxas1 C-termini (amino acids 258-331), different regions in BRCC45 (amino acids 161-169, 300-306, 316-346, and 353-382), MERIT40, RAP80 AIR regions (amino acids 287-292 and 325-329), Ub (P2), and the non-catalytic ubiquitin molecules from the ARISC(E33A)–RAP80:K63-Ub7 model displayed poor side chain density, and therefore the side chains of these regions were set to an occupancy of 0. Different BRCC45 regions (amino acids 131-177 and 222-256) and Ub (P2) from the ARISCΔC(E33A):K63-Ub7 model as well as various BRCC45 regions (amino acids 25-34, 131-180, and 202-266) from the ARISCΔC(E33A):K63-Ub4 model also displayed poor side chain density, and therefore the side chains of these regions were also set to an occupancy of 0. Gaps were left where direct connectivity between secondary structure elements could not be determined. The overall quality of the models was assessed using MolProbity^80^.

### DUB activity assays (IQF)

To assess enzymatic activity, purified ARISC WT and E33A complexes (at 10 nM) were incubated at 30°C in DUB reaction buffer containing 50 mM HEPES-NaOH pH 7.0, 100 mM NaCl, 0.1 mg/mL bovine serum albumin (BSA), 1 mM DTT, and 0.03% (v/v) Brji-35. Internally quenched fluorescent K63-linked diubiquitin substrate (K63-diUb IQF; Lifesensors, catalogue number: DU6303) at a concentration of 50 nM was used as a reporter for DUB activity. A total of 20 μL enzyme reactions were carried out in 384-well black flat-bottom low flange plates (Corning; catalogue number: 35373), and cleaved diUb was monitored by measuring fluorescent intensity (excitation: 544 nm, emission: 575 nm; dichroic mirror: 560 nm) every 30 s over 1 h in a Hidex Sense microplate reader. Following baseline correction of raw data values, the change in fluorescence over time was plotted using Prism 10 v10.1.0 (GraphPad Software).

### Gel-based ubiquitin chain cleavage assays

For experiments using WT and distally blocked K63-linked ubiquitin chains, purified FL ARISC, ARISCΔN, ARISCΔC, and ARISCΔNΔC variants, all ARISC mutants, ARISC–RAP80, ARISC–RAP80 AIR, and BRISC (at 5 nM or 10 nM, as described in the corresponding figure legends) were mixed with the indicated length and type of K63-linked ubiquitin chains (at 2 μM, 1 μM, or 0.5 μM, as described in the corresponding figure legends) in storage buffer containing 25 mM HEPES-NaOH pH 7.3, 150 mM NaCl, 10% (v/v) glycerol, and 1 mM TCEP. For experiments with fluorescently labelled ubiquitin substrates, ARISC, ARISC–RAP80, ARISC–RAP80 ΔUIMs, ARISC–RAP80 ΔZnF, and ARISC–RAP80 AIR (at 10 nM) were mixed with Alexa-Fluor 488 (AF488) labelled proximally or distally blocked K63-linked tetraUb (Ub4) chains (at 1.5 μM, 1 μM, or 0.3 μM, as described in the corresponding figure legends) in the same buffer indicated above. For experiments using WT K63-linked pentaUb (Ub5) and M1-linked tetraUb (Ub4) (MRC Protein Phosphorylation and Ubiquitylation Unit, DU20766) chains, ARISC (at 20 nM), ARISC–RAP80 (at 10 nM), AMSH* (at 100 nM), and OTULIN (at 10 nM) were mixed with the indicated length and type of ubiquitin chains (at 1 μM) in the same storage buffer. Enzyme reactions (30 μL final volume) were incubated at 30°C in a ProFlex PCR system (Life Technologies), and 8 μL samples were taken from each reaction at the indicated time points. Reactions were stopped with the addition of 3 μL 4x SDS-PAGE loading dye [240 mM Tris-HCl pH 6.8, 40% (v/v) glycerol, 8% (w/v) SDS, 0.04% (w/v) bromophenol blue, and 5% (v/v) β-Mercaptoethanol], and products were separated on 4-12% or 12% Nu-PAGE Bis-Tris gels (Invitrogen). Gels were run for 35-40 min at 200 V and room temperature in MES buffer, and stained using a silver stain kit (Bio-Rad) or an oriole stain kit (Bio-Rad) according to manufacturer’s instructions. Bands were visualised using an iBright FL 1500 gel imaging system (Thermo Fisher Scientific). For assays with AF488 labelled proximally and distally blocked K63-Ub4 variants, gels were scanned as described (excitation: 455-485 nm, emission: 508-557 nm) prior to being silver stained. The disappearance of each K63-Ub4 parent band was subsequently quantified using densitometry, and plotted as fraction of substrate consumed (%) using Prism 10 v10.1.0 (GraphPad Software).

### Spectral shift assays

Specific labelling of cysteine residues in ARISC(E33A)–RAP80, ARISC(E33A)^BRCC36(S98K)^–RAP80, ARISC(E33A)^Abraxas1(Δ42-55)^–RAP80, and ARISC(E33A)^BRCC45(ΔLoop)^–RAP80 was performed using a RED-MALEIMIDE 2^nd^ Generation labelling kit (NanoTemper Technologies). Briefly, each ARISC(E33A)–RAP80 complex (at 3 µM) was mixed with RED-MALEIMIDE 2^nd^ Generation dye (at 15 µM) in labelling buffer containing 17.7 mM sodium dihydrogen phosphate, 32.5 mM disodium phosphate, 100 mM NaCl, pH 7.0. Samples (50 μL final volume) were then incubated at room temperature for 45 min in the dark, and passed through a Zeba^TM^ Spin Desalting column (ThermoFisher Scientific) two times in order to remove excess dye. To measure the affinity of interaction between ARISC(E33A)–RAP80 variants and WT K63-linked hexaUb (Ub6) chains, each maleimide labelled ARISC(E33A)–RAP80 complex (at 40 nM) was mixed with K63-Ub6 in a 16-point, two-fold dilution series (20 µM-0 µM) in assay buffer containing 50 mM HEPES-NaOH pH 7.5, 150 mM NaCl, 0.5 mM DTT, 0.1 mg/mL BSA, and 0.005% (v/v) Tween-20. A total of 20 μL enzyme reactions were set up in 384-well Dianthus microplates (NanoTemper Technologies), and incubated for 30 min at room temperature. Measurements were performed in a Dianthus NT.23 instrument at 25°C, using the auto-excitation settings in a DI.Control software (v2.1.1; NanoTemper Technologies). Data were analysed using DI.Screening Analysis software (v2.1.1; NanoTemper Technologies). Following baseline correction of raw data values, binding affinity curves were generated by plotting the ratio of the fluorescence intensities at 670 nm and 650 nm against K63-Ub6 concentration, using the built-in equation for total binding (one site) in Prism 10 v10.1.0 (GraphPad Software).

### Intact mass spectrometry

Protein desalting and mass analysis was performed by liquid chromatography-mass spectrometry (LC-MS) using a Vanquish UPLC (Thermo Scientific) interfaced to a Orbitrap Exploris 240 mass spectrometer (Thermo Scientific). Purified WT and distally blocked K63-linked ubiquitin chains (20 μL at a concentration of 10 μM each) as well as AF488 labelled proximally and distally blocked tetraUb (Ub4) chains (15 μL at a concentration of 10 μM each) were diluted to 5 μM using 0.1% (v/v) formic acid, and 5 μL of each diluted sample was run on a MAbPac RP column (2.1 x 100 mm, Thermo Scientific). Buffer A was 0.1% (v/v) formic acid in water, buffer B was 0.1% (v/v) formic acid in acetonitrile, and the system flow rate was kept constant at 0.25 mL/min. The bound proteins were eluted by a gradient of 20%-45% buffer B in A over 1 min, and held at 45% buffer B in A for 1.5 min. The column was subsequently washed with a gradient of 45%-80% buffer B over 1.5 min, and held at 80% buffer B for 2 min before being re-equilibrated in 20% buffer B in A prior to the next injection. Data processing was performed using UniDec^81^.

### Cross-linking mass spectrometry

Purified ARISCΔC(E33A) and ARISC(E33A)–RAP80 complexes (at 5 μM) were mixed with 1.5-fold molar excess of WT K63-linked tri, tetra, penta, or heptaUb (Ub3, Ub4, Ub5, or Ub7) chains prior to cross-linking mass spectrometry experiments. In parallel, ARISC(E33A)–RAP80 was also mixed with distally blocked K63-Ub7 chains at the same concentrations outlined above. The mixtures were then incubated with 0.4 mM disuccinimidyl dibutyric urea (DSBU; Thermo Fisher Scientific) at room temperature for 45 min, and the reactions were quenched by adding 50 mM Tris-HCl pH 7.5. Protein digestion was carried out as previously described^82^.

The peptides were then analysed by liquid chromatography-tandem mass spectrometry (LC-MS/MS) on an Orbitrap Eclipse Tribrid mass spectrometer interfaced with a Vanquish Neo liquid chromatography system (Thermo Fisher Scientific), which were operated using Xcalibur v4.7 and Tune v4.2. Prior to LC separation, peptides were desalted using a Trap column (300 µm x 5 mm, µPrecolumn, 5 µm particles, Acclaim^TM^ PepMap^TM^ 100 C18, Thermo Fisher Scientific) at room temperature. After washing the Trap column with 0.1% (v/v) formic acid, the dipeptides were eluted at a flow rate of 0.25 nL/min from the Trap column onto an analytical EASY-Spray column (75 µm x 500 mm, 2 µm particles, Acclaim^TM^ PepMap^TM^ 100 C18, Thermo Fisher Scientific) using a linear gradient of 2-50% (v/v) solvent B [0.1% (v/v) formic acid in 80% (v/v) acetonitrile] in solvent A [0.1% (v/v) formic acid in water] over approximately 90 min at 45°C. The mass spectrometer was operated in data-dependent acquisition mode. The survey scan range was set to m/z 380-1450 in profile mode, with a resolution of 120,000 (at m/z 200), standard target value, and maximum injection time of 50 ms. The cycle time was adjusted to 3 s. Precursor ions with charge states 3-8+ were isolated using a 1.4 m/z window and fragmented by HCD using optimized stepped normalized collision energies (30 ± 6). Fragment ion scans were acquired at a resolution of 60,000, using an AGC target of 1 x 10^5^, a maximum injection time of 120 ms, and by scanning from 200–2000 m/z. The dynamic exclusion was enabled for 30 s (including isotopes). Data analysis was performed in XlinkX (Proteome Discoverer v3.0, Thermo Scientific). RAW files were converted to mgf files using Proteome Discoverer (Thermo Scientific) and used in the same software to identify cross-linked peptides. Sequence-specific fasta files were used for database searches. xiView^83^ was used to visualise cross-linking results.

### Western blotting

Cells were lysed in 2x LDS sample buffer (Thermo Scientific; catalogue number: NP0007) supplemented with benzonase (Sigma-Aldrich; catalogue number: E1014) and MgCl_2_ (1.5 mM). Protein samples were run on NuPAGE™ 4-12% Bis-Tris gels (Thermo Fisher Scientific; catalogue number: NP0335BOX), and subsequently transferred onto 0.2 µm nitrocellulose blotting membrane (Cytiva; catalogue number: 10600004) using wet transfer at 100 V at 4 °C for 2 h. The membranes were blocked with 5% milk in 1x PBS for 1 h at RT followed by O/N incubation with primary antibodies. The following primary antibodies were used: Abraxas1 (Bethyl laboratories; catalogue number: A302-180A); BRCC36 (Abcam; catalogue number: ab108411); BRCC45 (Abcam; catalogue number: ab177960) GAPDH (Cell Signaling; catalogue number: 2118S). Following three washes with 1x PBST, membranes were incubated with HRP-conjugated anti-rabbit (Amersham; catalogue number: NA934; 1:2500 dilution) secondary antibody at RT for 1 h. Membranes were washed five times with 1x PBST, and protein bands were developed on autoradiogram using Western lightning plus-ECL substrate (Revvity Health Sciences Inc.; catalogue number: NEL105001EA).

### Immunofluorescence

Cells were seeded (0.5 x 10^6^) on coverslips or 8-well chamber slides (Celltreat; catalogue number: 229168) one day before exposure to ionizing radiation (IR). Cells were irradiated with 10 Gy using Gammacell 40 Exactor and recovered for 4 h before fixation. Cells were washed twice with ice-cold 1x PBS and pre-extracted for 10 min on ice using CSK buffer (10 mM PIPES pH 7.0, 100 mM NaCl, 300 mM Sucrose, 3 mM MgCl_2_, 1 mM EGTA, and 0.5% Triton X-100). Cells were washed twice with 1x PBS and fixed with 4% paraformaldehyde (Electron Microscopy Sciences; catalogue number: 50-980-495) diluted in 1x PBS for 20 min on ice. Following three washes with 1x PBS, cells were permeabilized with 0.5% Triton X-100 for 10 min on ice. Blocking was done using blocking buffer (5% BSA with 10% goat serum in 1x PBS) at 4°C for 1 h. Cells were incubated with primary antibodies O/N at 4°C. The following primary antibodies were used: anti-HA.11 Epitope Tag (BioLegend; catalogue number: 901516); Phospho-Histone H2A.X (Ser139) (Cell Signaling; catalogue number: 9718S). Cells were washed three times with 1x PBS and incubated with mouse AlexaFluor 488- (Thermo Scientific; catalogue number: A-11029) and rabbit AlexaFluor 647- (Thermo Scientific; catalogue number: A-21245) conjugated secondary antibodies for 1 h at RT. Immunofluorescence experiments using BRCC45 mutants were carried out in BRCC45 KO HEK293T cells transfected with sgRNA-resistant FLAG tagged WT or mutant pCMV6-BRCC45 expression plasmids. Cells were processed for IR and subsequent immunofluorescence experiments using anti FLAG antibody (Cell Signaling; catalogue number: 14793) 48 h after transfection. Cells were washed three times with 1x PBS followed by DAPI staining (Sigma-Aldrich; catalogue number: D9542; 4 µg/mL diluted in 1x PBS). Slides were mounted with ProLong Gold Antifade mountant (Thermo Scientific; catalogue number: P36930) and cured O/N at RT. Immunofluorescence images were acquired by Plan Apo VC 60× Oil DIC N2 objective lens of Nikon Eclipse 80i fluorescence microscope equipped with a CoolSnap MYO camera. IRIF were quantified using CellProfiler v4.2.5 image analysis software^84^ and graphs were generated in GraphPad Prism v10.

### Data analysis and figures generation

Structural representations shown in **Supplementary Fig. 8** were created in PyMOL (The PyMOL Molecular Graphic System, v3.1.3.1 Schrödinger, LLC). All other structural models, surface and electron density map representations, and schematics were obtained using UCSF Chimera X v1.6.1^69,70^. Structural overlays depicted in **Fig. 3a**, **Fig. 5c-e**, and **Supplementary Fig. 8** were performed using the program Superpose^85^, available within the CCP4 suite (Collaborative Computational Project, Number 4, 1994). Overlays of the Abraxas1:ubiquitin and BRCC45:ubiquitin AlphaFold 3 models as well as of BRCC45 UEV-C (from the ARISC(E33A)–RAP80:K63-Ub7 complex in this study) with UbcH5b∼Ub (PDBid = 3A33) shown in **Fig. 7d**, **Supplementary Fig. 9e**, and **Supplementary Fig. 14c,d** were obtained using UCSF Chimera X v1.6.1^69,70^. The European Bioinformatics Institute (EBI) web service of the PISA (Proteins, Interfaces, Structures and Assemblies) software (https://www.ebi.ac.uk/pdbe/pisa/)^86^ was used to analyse the BRCC36^MPN^:K63-Ub2 (from the ARISC(E33A)–RAP80:K63-Ub7 complex in this study) and AMSH-LP^MPN^:K63-Ub2 (PDBid = 2ZNV) structures. The electrostatic potential molecular surfaces shown in **Fig. 7f** were generated using the coulombic command in UCSF Chimera X v1.6.1^69,70^. Sequence alignment shown in **Supplementary Fig. 9f** was performed with Clustal Omega (https://www.ebi.ac.uk/jdispatcher/msa/clustalo)^87^, and edited and displayed using the Consurf web server (https://consurf.tau.ac.il/consurf_index.php)^88^. The final model shown in **Fig. 8** was created with BioRender (https://www.biorender.com/). All figure panels were assembled using Affinity Designer v2 (https://affinity.serif.com/en-gb/designer/).

### Quantification and statistical analysis

Unpaired two-tailed t test was performed to compute statistical significance in GraphPad Prism 10.

## Data availability

Cryo-EM maps and associated structural models have been deposited to the Electron Microscopy Data Bank and Protein Data Bank under accession codes EMD-55122, EMD-55090, EMD-55069, EMD-55070, EMD-55071, EMD-55072, EMD-55073, EMD-55074, and PDBid: 9SQY (for the ARISC(E33A)–RAP80:K63-Ub7 complex); EMD-55119, EMD-55079, EMD-55080, EMD-55081, EMD-55077, EMD-55082, and PDBid: 9SQW (for the ARISCΔC(E33A):K63-Ub7 complex); EMD-55118, EMD-55085, EMD-55086, EMD-55088, EMD-55089, and PDBid: 9SQV (for the ARISCΔC(E33A):K63-Ub4 complex). Raw cross-linking mass spectrometry data have been deposited to the ProteomeXchange Consortium via the PRIDE partner repository with the dataset identifier PXD068769. Reviewers can access the deposited dataset by using the following login details: username =; password = EYr4Tv9fayH8. All unique reagents are available upon request. Source Data are provided with this paper.

## Acknowledgments

We thank members of the Zeqiraj and Greenberg labs for critical discussion. This work was supported by a UK Research and Innovation (UKRI)-Biotechnology and Biological Sciences Research Council (BBSRC) Discovery Fellowship (BB/Z51522X/1) to M.F., a Wellcome Trust Senior Fellowship (222531/Z/21/Z) to E.Z., a UKRI-Medical Research Council (MRC) grant (MR/T029471/1) and a Basser External Research grant to E.Z. and M.F., National Institutes of Health (NIH) R01s (138835 and 1774904) and funds from the Basser Center for BRCA to R.A.G., and a Sir Henry Dale Fellowship (220628/Z/20/Z) jointly funded by the Wellcome Trust and Royal Society to A.N.C. The Astbury cryo-EM Facility is funded by a University of Leeds ABSL award and a Wellcome Trust grant (221524/Z/20/Z), and fundings from the National Institute for Health and Care Research (NIHR) Biomedical Research Center (NIHR200633) and the Wellcome Trust (223810/Z/21/Z) supported the purchase of mass spectrometry instrumentation. The Dianthus NT.23 instrument is funded by a UKRI-MRC World Class Labs award (MC_PC_MR/Y002482/1).

## Contributions

M.F. performed protein production, enzyme activity assays, spectral shift assays, and cryo-EM data collection, processing, and model building. A.D. performed genetic reconstitution experiments and immunofluorescence analyses. O.D. contributed to cryo-EM data collection and processing. H.P. carried out the preparation and labelling of proximally and distally blocked tetraubiquitin chains. U.M.S., J.L., and L.J.C. were involved with protein production. M.F., S.R.G., and A.N.C performed XL-MS analyses, while G.W. and A.N.C performed intact MS. F.C. was involved with mutants design. M.F., A.D., R.A.G., and E.Z. performed the majority of experimental design and data interpretation. M.F. wrote the first draft of the manuscript with input from E.Z., A.D., and R.A.G.

## Supplementary Information

**Supplementary Fig. 1:**
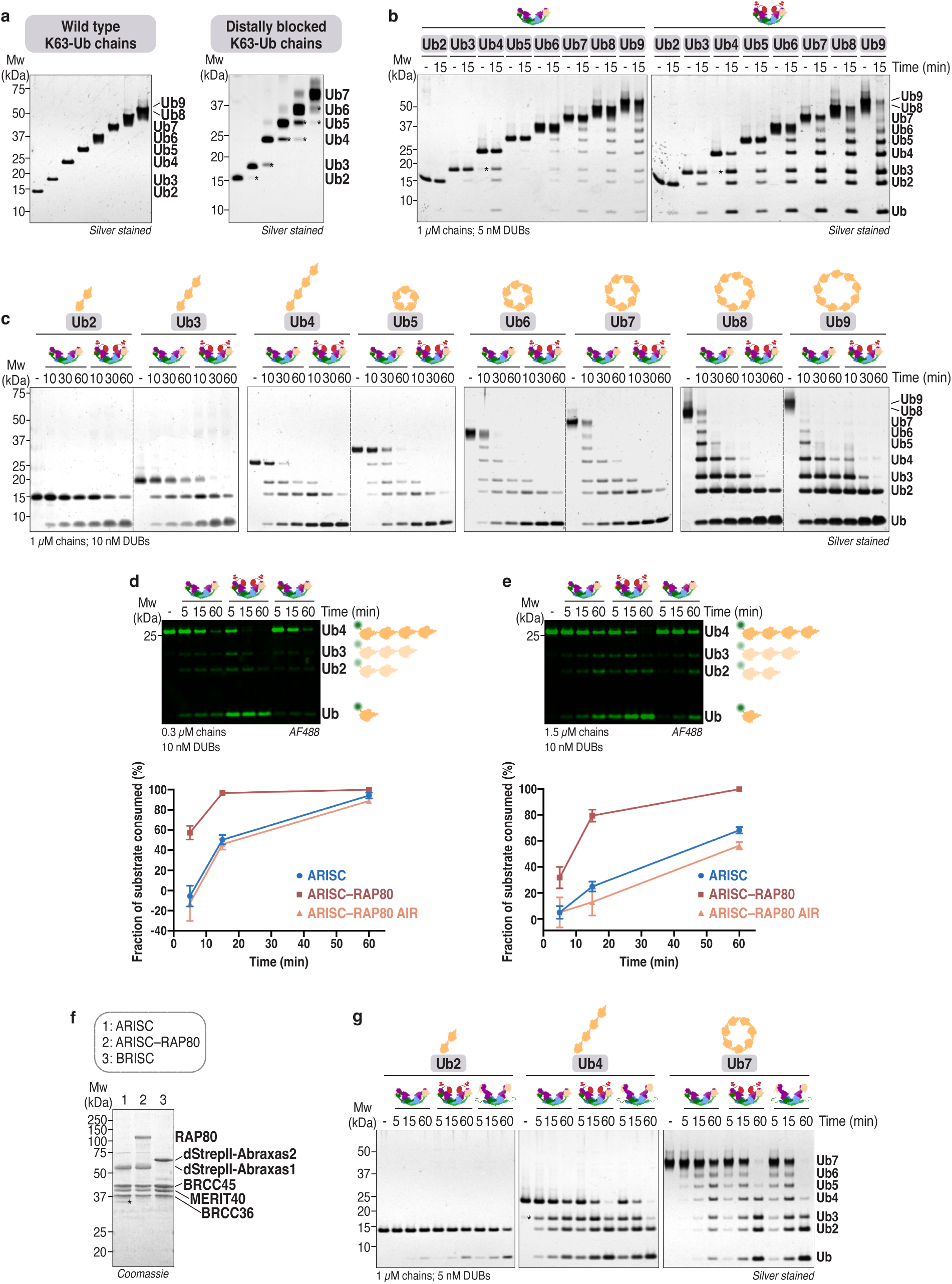
Preparation of protein complexes and ubiquitin chains, and assessment of DUB activities. **a**, SDS-PAGE analysis of wild type (WT) and distally blocked K63-linked ubiquitin chains. **b,** Reactions depicted in Fig. 1c (at 15 min time point) were re-loaded side-by-side on the same gels to better highlight differences in substrate cleavage between ARISC and ARISC–RAP80. The final concentration of K63-linked ubiquitin chains was 1 µM, while ARISC and ARISC–RAP80 were assayed at a final concentration of 5 nM. Cleavage activity was analysed by SDS-PAGE and silver staining. Data are representative of two independent experiments. **c,** K63-linked ubiquitin chains (1 µM) were incubated with ARISC or ARISC–RAP80 (10 nM) for the indicated time points. Cleavage activity was analysed as in **b**. Data are representative of three independent experiments. **d,e,** Alexa-Fluor 488 (AF488) labelled distally (AF488-^Cys^Ub4^K63R^) blocked K63-Ub4 chains (**d,** 0.3 µM; **e,** 1.5 µM) were incubated with ARISC, ARISC–RAP80, or ARISC–RAP80 AIR (10 nM) for the indicated time points. Cleavage activity was analysed by SDS-PAGE, and gels were subsequently scanned as described in Fig. 1f (*top*). The disappearance of the K63-Ub4 parent band was quantified using densitometry, and plotted as fraction of substrate consumed (%). Data points are mean ± SEM of two independent experiments (*bottom*). **f,** SDS-PAGE analysis of ARISC, ARISC–RAP80, and BRISC complexes. dStrepII, double StrepII tag. * indicates Abraxas1 degradation product. **g,** K63-Ub2, -Ub4, and -Ub7 chains (1 µM) were incubated with ARISC, ARISC–RAP80, or BRISC (5 nM) for the indicated time points. Cleavage activity was analysed as in **b**. Data are representative of two independent experiments. * indicates lower molecular weight ubiquitin species. Ub, ubiquitin; DUB, deubiquitylating enzyme.

**Supplementary Fig. 2:**
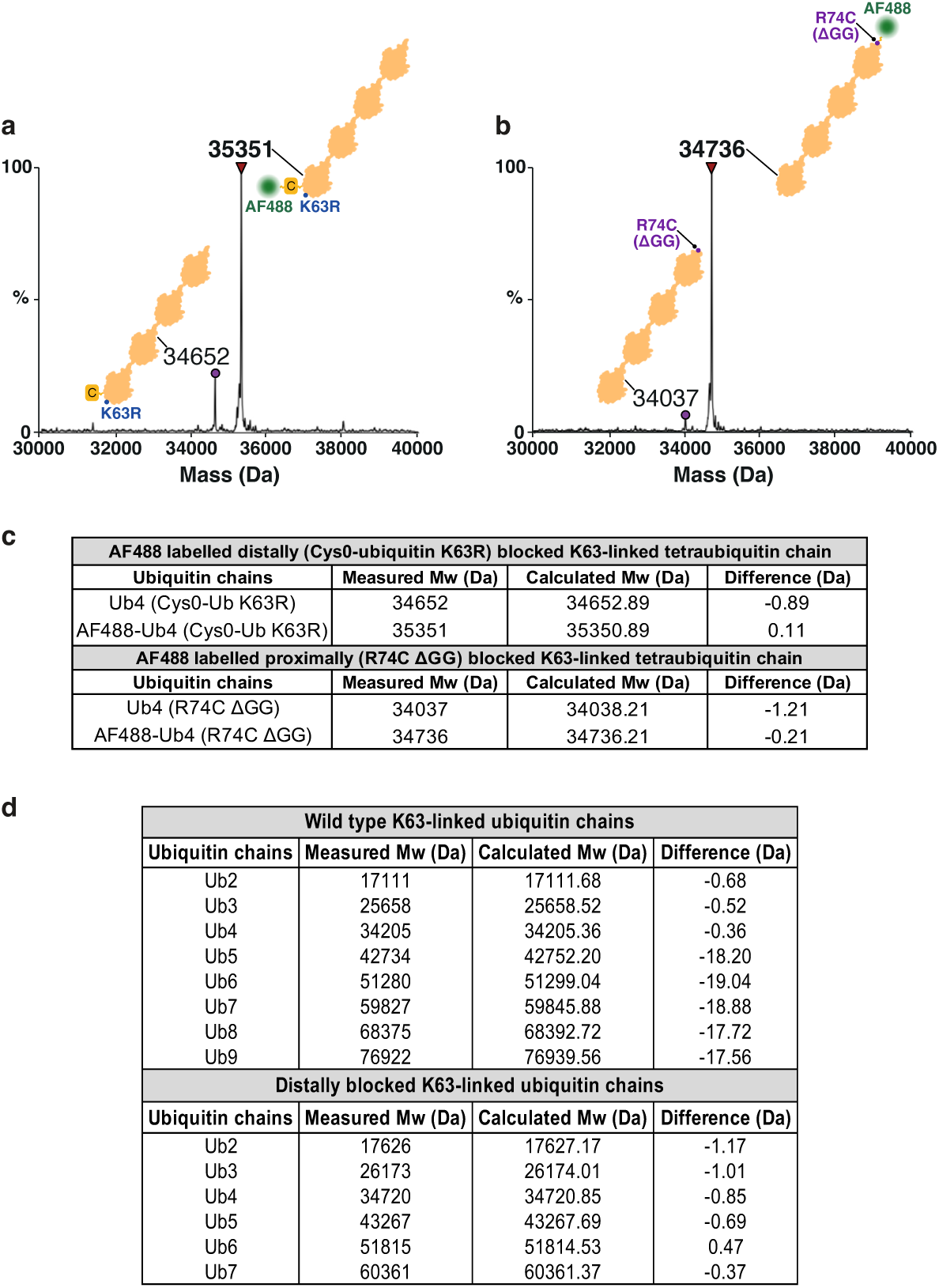
Validation of Alexa-Fluor 488 labelled distally and proximally blocked K63-linked tetraubiquitin chains by mass spectrometry analyses. **a,b**, Intact mass spectra of Alexa-Fluor 488 (AF488) labelled distally (**a**) and proximally (**b**) blocked K63-linked Ub4 chains. Observed masses and schematics for the different ubiquitin chains are shown. The positions of the K63R and R74C ΔGG mutants, of the N-terminal cysteine (Cys0), and of the AF488 fluorophore are indicated. Data are representative of a single experiment. **c,** Table summarising the measured and calculated masses from intact mass spectrometry analyses of AF488 labelled distally (AF488-^Cys^Ub4^K63R^) and proximally (Ub4^R74CΔGG^-AF488) blocked K63-Ub4 chains. Measured and calculated masses for the corresponding unlabelled species are also indicated. **d,** Table summarising the measured and calculated masses from intact mass spectrometry analyses of WT and distally blocked K63-linked ubiquitin chains (see **Supplementary Fig. 3**). A mass difference of ∼ 18 Da in WT Ub5-Ub9 chains indicates the lack of a water molecule.

**Supplementary Fig. 3:**
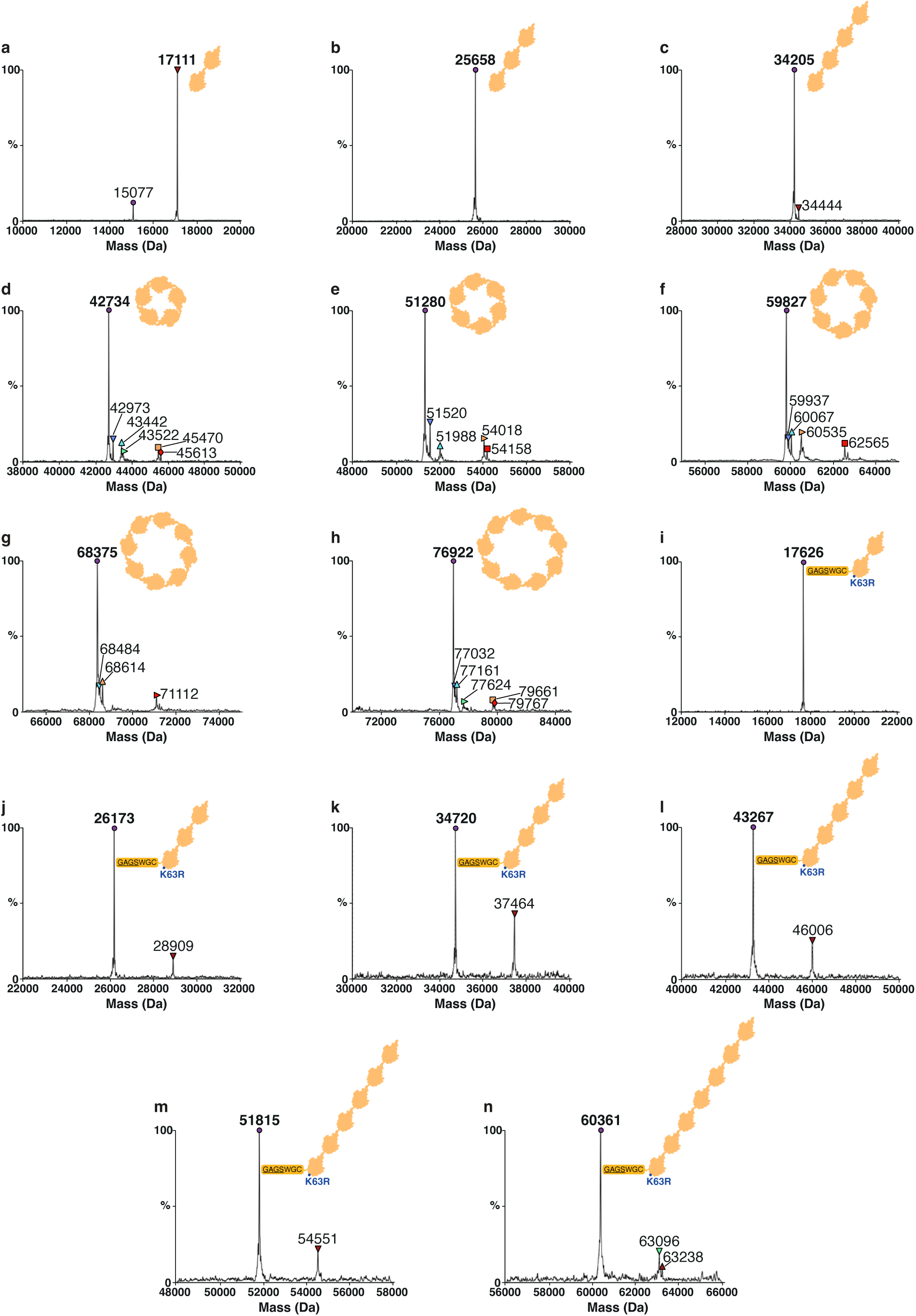
Validation of wild type and distally blocked K63-linked ubiquitin chains by mass spectrometry analyses. **a-h**, Intact mass spectrometry analyses of wild type K63-linked Ub2 (**a**), Ub3 (**b**), Ub4 (**c**), Ub5 (**d**), Ub6 (**e**), Ub7 (**f**), Ub8 (**g**), and Ub9 (**h**) chains. **i-n,** Intact mass spectra of distally blocked K63-linked Ub2 (**i**), Ub3 (**j**), Ub4 (**k**), Ub5 (**l**), Ub6 (**m**), and Ub7 (**n**) chains. Observed masses and schematics for the different types and lengths of ubiquitin chains are shown. Amino acids left at the N-terminus of ubiquitin K63R after cleavage of the 8xHis tag, and the position of the K63R mutant, are indicated. Data are representative of a single experiment (see **Supplementary Fig. 1a**).

**Supplementary Fig. 4:**
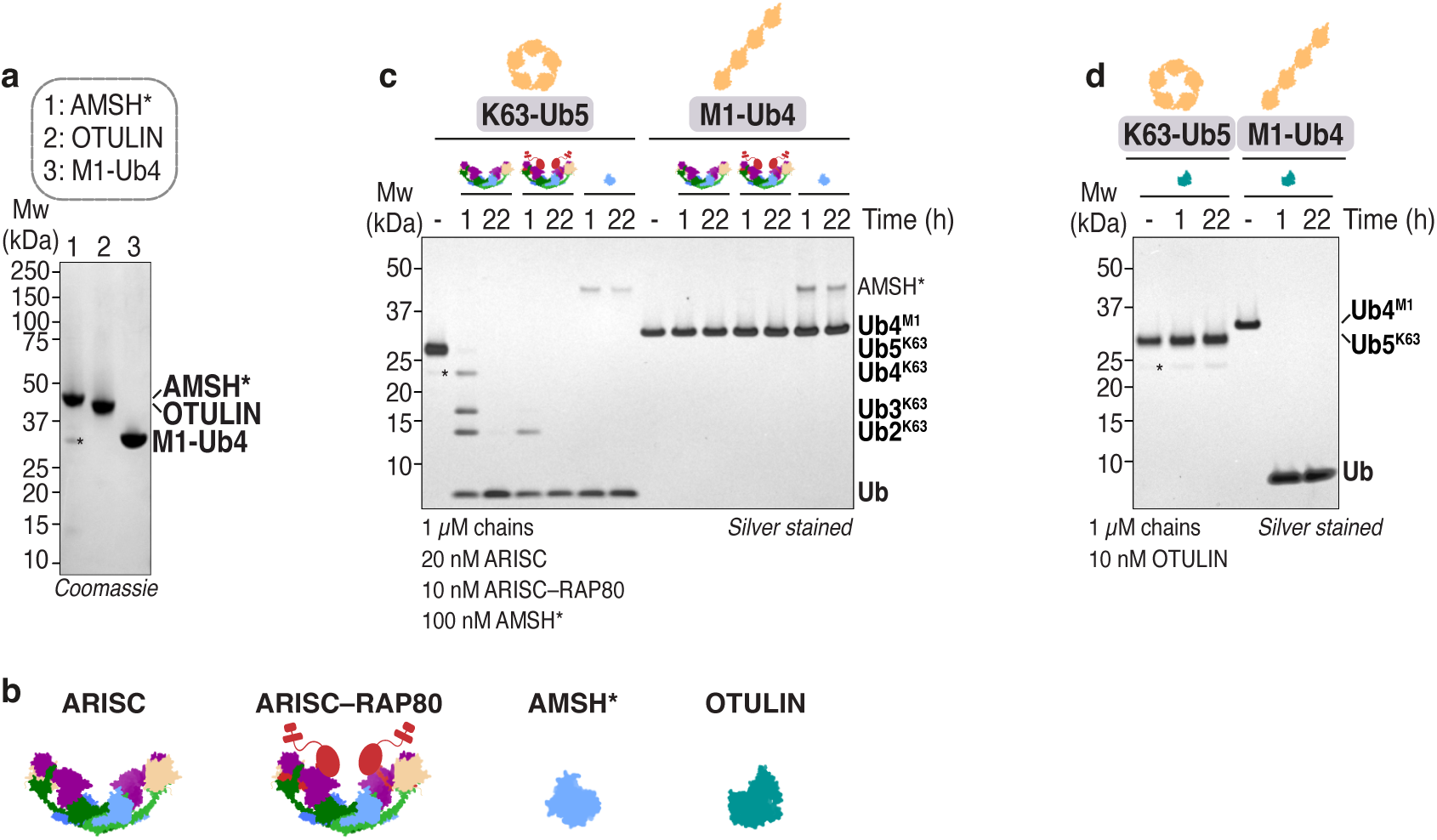
Biochemical analyses of wild type K63-linked pentaubiquitin chains. **a**, SDS-PAGE analysis of AMSH* (a STAM2-AMSH fusion), OTULIN, and M1-linked tetraUb (Ub4) chains. * indicates AMSH* contaminant or degradation product. **b,** Schematics of indicated proteins and protein complexes. **c,d,** Cyclical K63-Ub5 and linear M1-Ub4 chains (1 µM) were incubated with ARISC (20 nM), ARISC–RAP80 (10 nM), and AMSH* (100 nM) (**c**) or OTULIN (10 nM) (**d**) for up to 22 h. Cleavage activity was analysed by SDS-PAGE and silver staining. Data are representative of two independent experiments. * indicates lower molecular weight ubiquitin species. Ub, ubiquitin.

**Supplementary Fig. 4:**
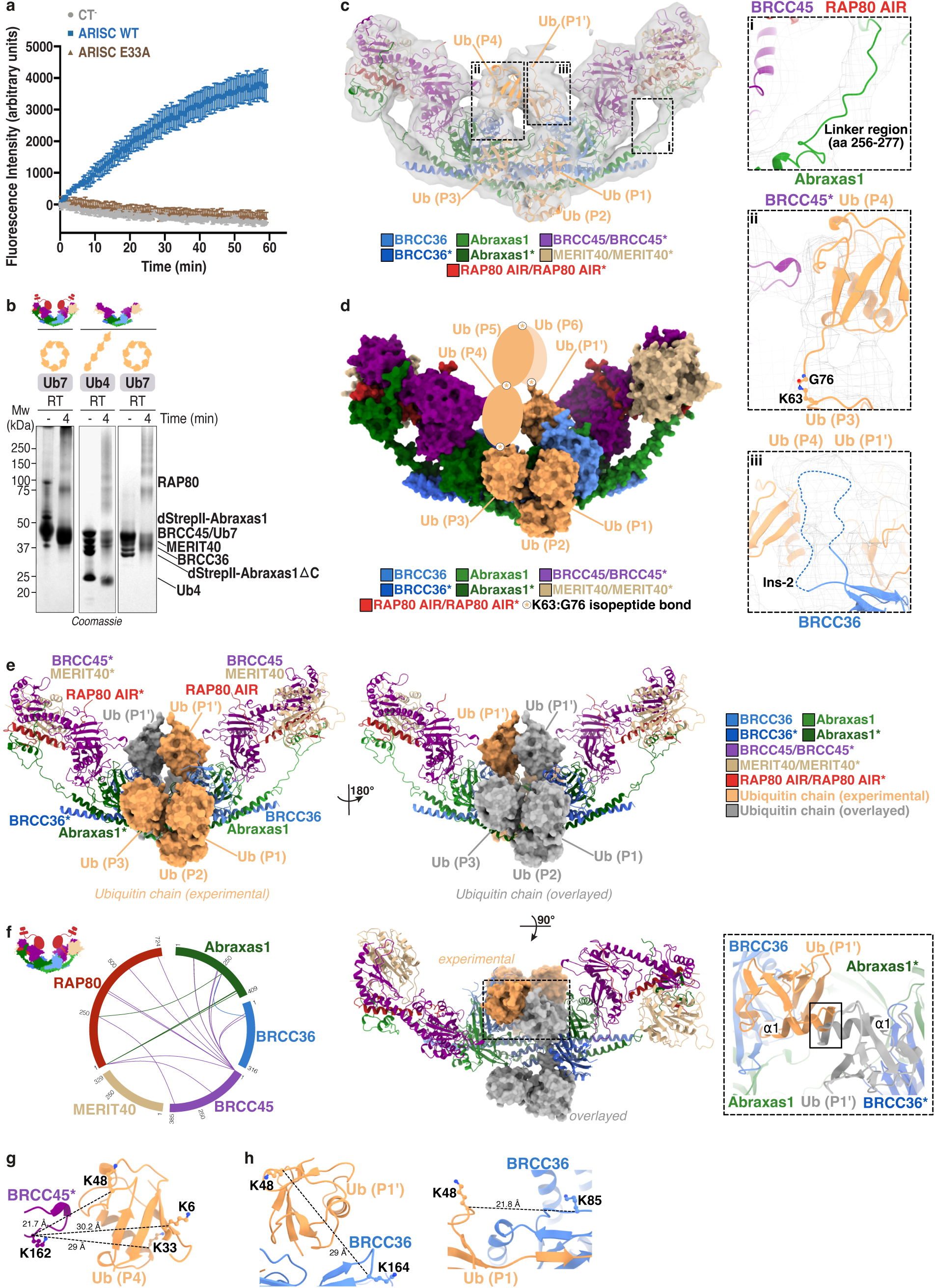
Preparation of ARISC and ARISC–RAP80 complexes for structural studies, and analysis of the ARISC(E33A)–RAP80:K63-Ub7 cryo-EM structure. **a**, Deubiquitylase activity of ARISC variants (10 nM) against K63-diUb IQF (50 nM). Data points are mean ± SEM of three independent experiments carried out in technical duplicates. CT^-^, negative control; WT, wild type. **b,** Glutaraldehyde cross-linking and SDS-PAGE analysis of ARISC(E33A)–RAP80 or ARISCΔC(E33A) and ubiquitin chains. RT, room temperature; dStrepII, double StrepII tag. **c,** The ARISC(E33A)–RAP80:K63-Ub7 ab-initio map (contour level = 0.105) is depicted in transparent surface. Cartoon models were rigid-body fitted into the density, and colored as in Fig. 1a and Fig. 2a. Dashed black rectangles highlight regions not well resolved in the final composite map (*left*). Close-up views and structural details of the Abraxas1 linker region (i), putative BRCC45*:Ub (P4) interface (ii), and BRCC36 Ins-2 (iii). The ab-initio map is depicted as mesh; Ub (P3) K63^NZ^ and Ub (P4) G76^C^ are positioned at isopeptide bond distance (1.3 Å), and missing residues are depicted with a dashed line (*right*). Ins-2, insertion-2. **d,** Surface representation of ARISC(E33A)–RAP80:K63-Ub7, with models colored as in **c**. Ubiquitin molecules not visualised in the cryo-EM map are depicted as orange ovals. AIR, Abraxas1-interacting region. **e,** The ARISC(E33A)–RAP80:K63-Ub7 structure is shown in three orientations, with a close-up view depicting the BRCC36 and BRCC36* active sites. The BRCC36 molecule and bound ubiquitin chains were overlayed onto the BRCC36* protomer. ARISC–RAP80 subunits and ubiquitin chains are colored as in **c**; the ubiquitin moieties modelled onto BRCC36* are colored grey. The black rectangle in the close-up view highlights a partial clash between ⍺1 of the two Ub (P1’) molecules. **f,** Chemical cross-linking and mass spectrometry analysis of ARISC(E33A)–RAP80. Proteins and cross-links are depicted as in Fig. 2d**-h**. Communal cross-links from two independent experiments are shown. **g,h,** Close-up views and structural details of BRCC45*:Ub (P4) (**g**), BRCC36:Ub (P1’), and BRCC36:Ub (P1) (**h**) interfaces. Structural models are colored as in **c**. Cross-linked residues are shown as ball & sticks. Distances are indicated with dashed black lines and measured in Ångstrom (Å). Ub, ubiquitin.

**Supplementary Fig. 6:**
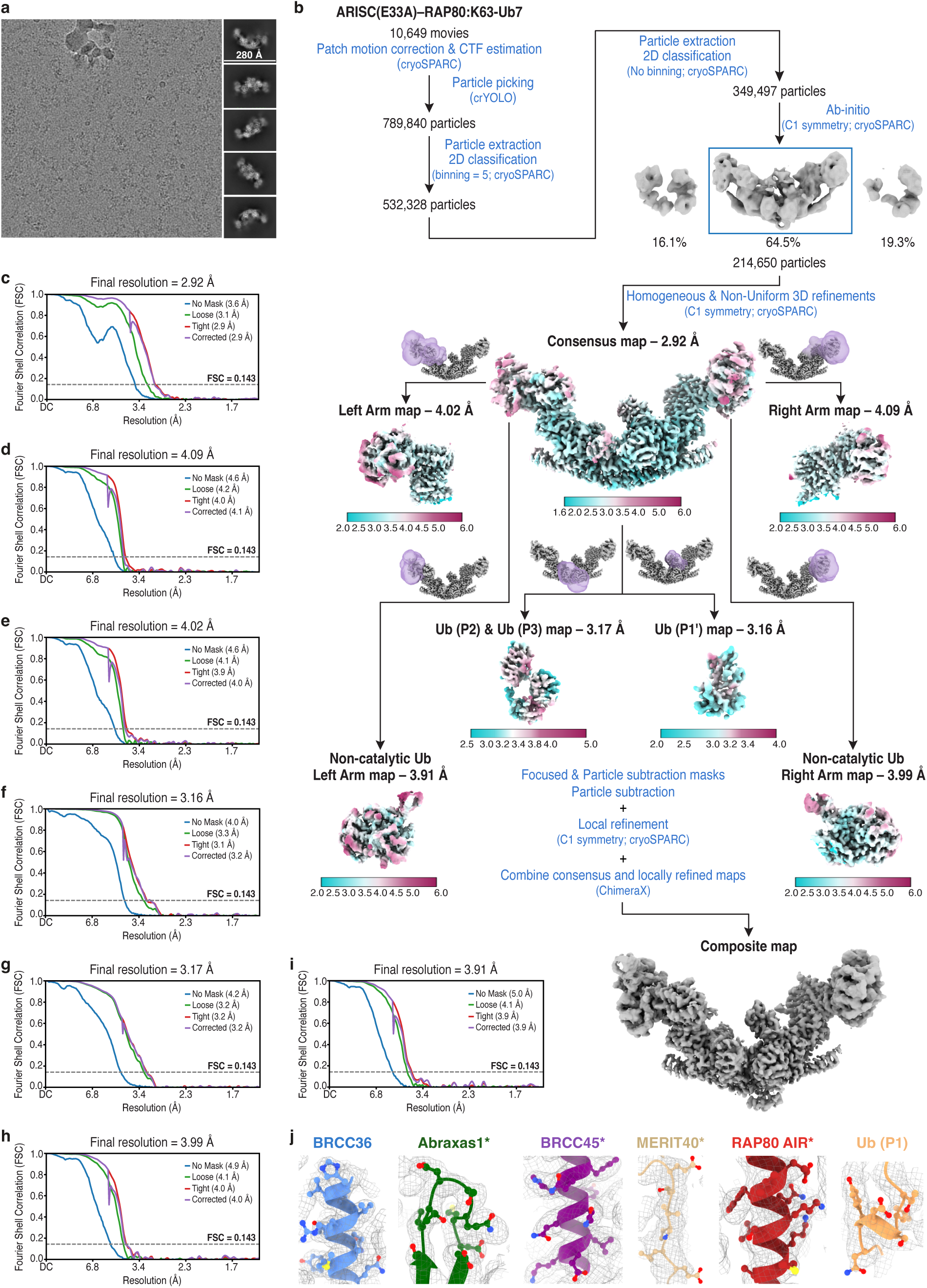
Cryo-EM analysis of the ARISC(E33A)–RAP80:K63-Ub7 complex. **a**, Representative micrograph (out of 10,649 collected movies) and 2D class averages (out of 349,497 selected particles) for the ARISC(E33A)–RAP80:K63-Ub7 dataset. **b,** Flowchart of data processing strategy. The boxed ab-initio model was selected for subsequent processing. The final consensus and locally refined maps are shown and colored by local resolution. See **Supplementary Table 1** for details. **c-i,** Fourier shell correlation (FSC) curves for the final ARISC(E33A)–RAP80:K63-Ub7 consensus map (**c**), and for the locally refined right (**d**) and left (**e**) arm, Ub (P1’) (**f**), Ub (P2) & Ub (P3) (**g**), and the non-catalytic ubiquitin right (**h**) and left (**i**) arm maps. Resolutions were calculated using the gold-standard FSC cutoff at 0.143 frequency. **j,** Representative density regions (at a contour level of 0.137) from the ARISC(E33A)–RAP80:K63-Ub7 composite map for the different components of the complex. Cartoon models are colored as in Fig. 1a and Fig. 2a, and residues are shown as ball & sticks. AIR, Abraxas1-interacting region; Ub, ubiquitin.

**Supplementary Fig. 7:**
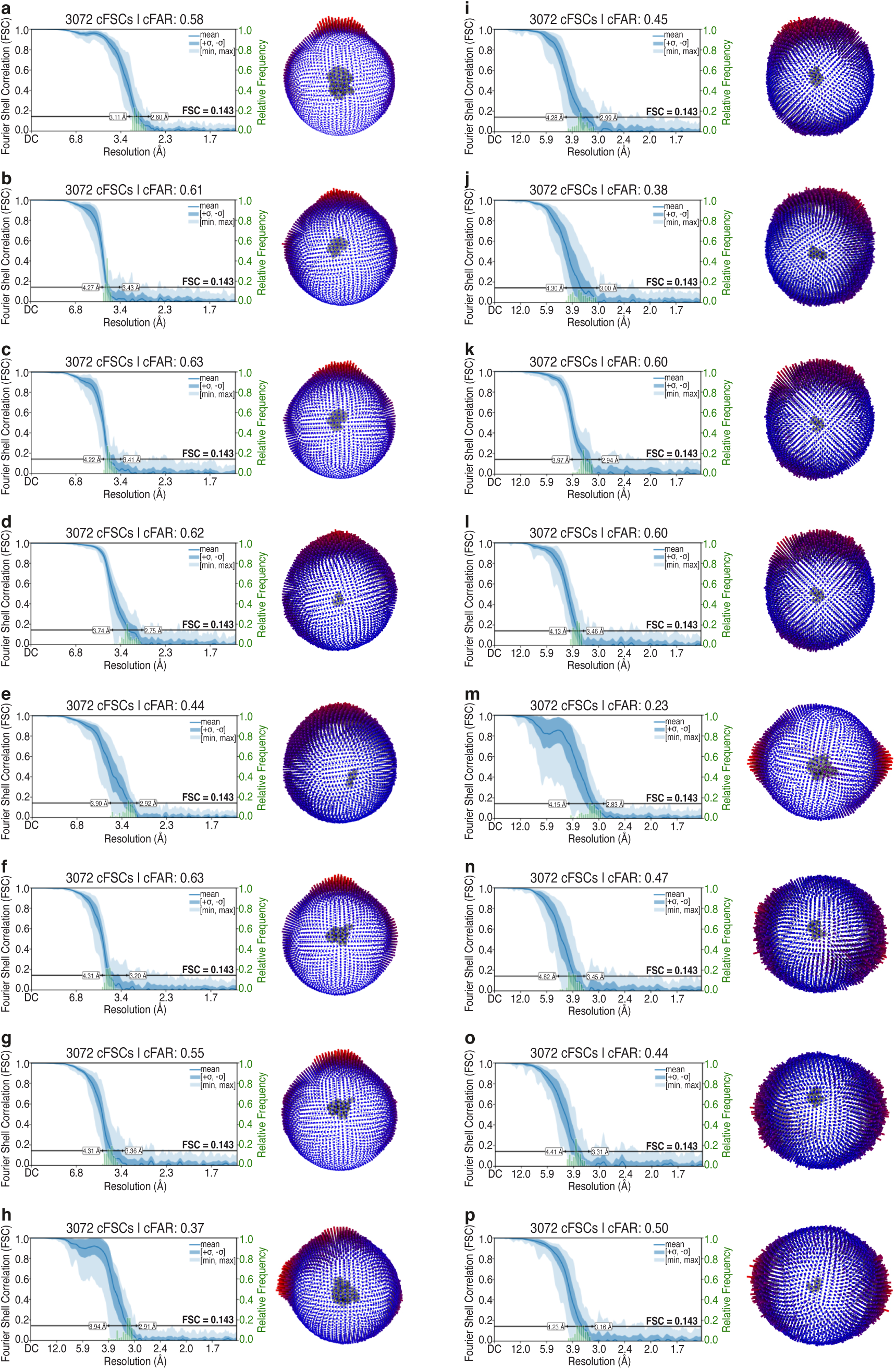
Cryo-electron microscopy validation of the ARISC(E33A)–RAP80:K63-Ub7, ARISCΔC(E33A):K63-Ub7, and ARISCΔC(E33A):K63-Ub4 datasets. **a-g**, Plots of conical Fourier shell correlation (FSC) Area Ratio (cFAR) (*left*) and Euler angle distribution of particles (*right*) in the ARISC(E33A)–RAP80:K63-Ub7 consensus map (**a**), and in the locally refined right (**b**) and left (**c**) arm, Ub (P1’) (**d**), Ub (P2) & Ub (P3) (**e**), and non-catalytic right (f) and left (**g**) arm maps (see **Supplementary Fig. 6**). **h-l,** Plots of cFAR (*left*) and Euler angle distribution of particles (*right*) in the ARISCΔC(E33A):K63-Ub7 consensus map (**h**), and in the locally refined right (**i**) and left (**j**) arm, Ub (P1’) (**k**), and Ub (P2) & Ub (P3) (**l**) maps (see **Supplementary Fig. 12**). **m-p**, Plots of cFAR (*left*) and Euler angle distribution of particles (*right*) in the ARISCΔC(E33A):K63-Ub4 consensus map (**m**), and in the locally refined right (**n**) and left (**o**) arm, and Ub (P1’) (**p**) maps (see **Supplementary Fig. 13**). Rod heights (*right*) are proportional to the number of particles in each direction. cFAR (*left*) quantifies the variance of directional half-map Fourier correlations across the viewing sphere ± 1 standard deviation from the mean (area within the light blue lines). Histograms of directional resolutions sampled over the cFAR curves and crossing at the gold-standard FSC cutoff of 0.143 frequency (green bars) are also indicated. cFAR values equal to or higher than 0.5 indicate no significant preferred particle orientation.

**Supplementary Fig. 8:**
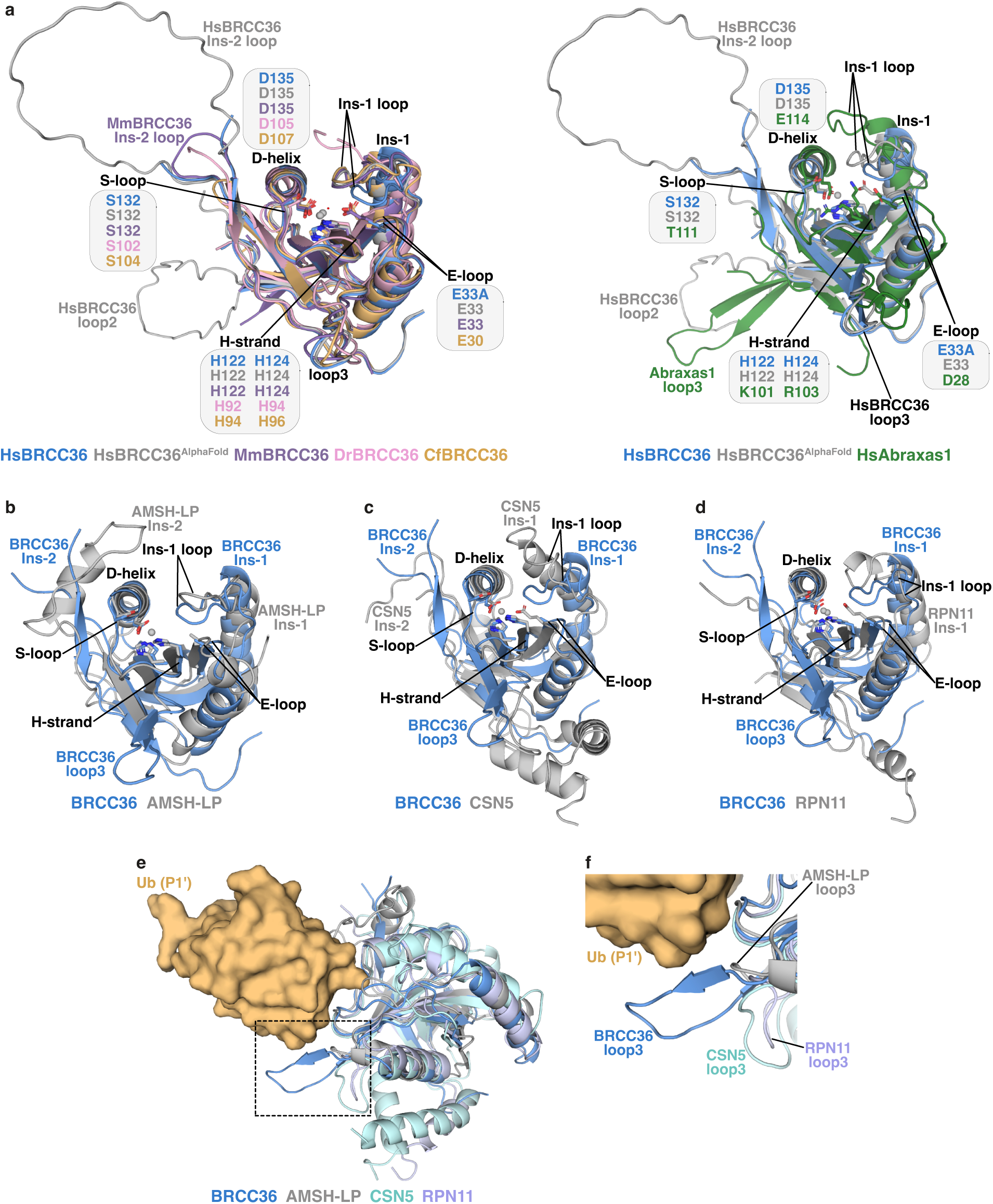
Structural analysis of BRCC36 and Abraxas1 subunits. **a**, Superimposition of the BRCC36 MPN domain from human (Hs; from the ARISC(E33A)–RAP80:K63-Ub7 complex and AlphaFold 3 model), mouse (Mm; PDBid = 6GVW), zebrafish (Dr; PDBid = 5CW6), and ant (Cf; PDBid = 5CW3) (*left*) or between the human BRCC36 MPN domain (Hs; from the ARISC(E33A)–RAP80:K63-Ub7 complex and AlphaFold 3 model) and the corresponding region of human Abraxas1 (Hs; from the ARISC(E33A)–RAP80:K63-Ub7 complex) (*right*). Structures are shown as cartoon and colored as indicated; active site residues are labelled and shown as sticks. Representative Zn^2+^ atoms obtained from the human (*left and right*), zebrafish, and ant (*left*) structures are depicted as grey spheres; the catalytic water molecule obtained from the mouse structure (*left*) is shown as a red sphere. Secondary structural elements are indicated. **b-d,** Overlays between the BRCC36 MPN domain (from the ARISC(E33A)–RAP80:K63-Ub7 complex) and the corresponding regions of AMSH-LP (PDBid = 2ZNV; RMSD = 2.4 Å) (**b**), CSN5 (PDBid = 4F7O; RMSD = 1.7 Å) (**c**), and RPN11 (PDBid = 4O8X; RMSD = 2.0 Å) (**d**). Structures are shown as cartoon and colored as indicated; active site residues are shown as sticks. Representative Zn^2+^ atoms obtained from the ARISC(E33A)–RAP80:K63-Ub7 (**b-d**) and RPN11 (**d**) structures are depicted as grey spheres. Secondary structural elements are indicated. Ins, insertion. **e,** Superimposition of the BRCC36 (from the ARISC(E33A)–RAP80:K63-Ub7 complex), AMSH-LP (PDBid = 2ZNV), CSN5 (PDBid = 4F7O), and RPN11 (PDBid = 4O8X) structures. Models are shown as cartoon and colored as indicated; the Ub (P1’) molecule (obtained from the ARISC(E33A)–RAP80:K63-Ub7 complex) is shown as surface and colored as in Fig. 2a. A dashed black rectangle highlights the position of loop3 relative to Ub (P1’). **f,** Close-up view of loop3 in the BRCC36, AMSH-LP, CSN5, and RPN11 structures shown in **e**. Models are depicted and colored as in **e**. Ub, ubiquitin.

**Supplementary Fig. 9:**
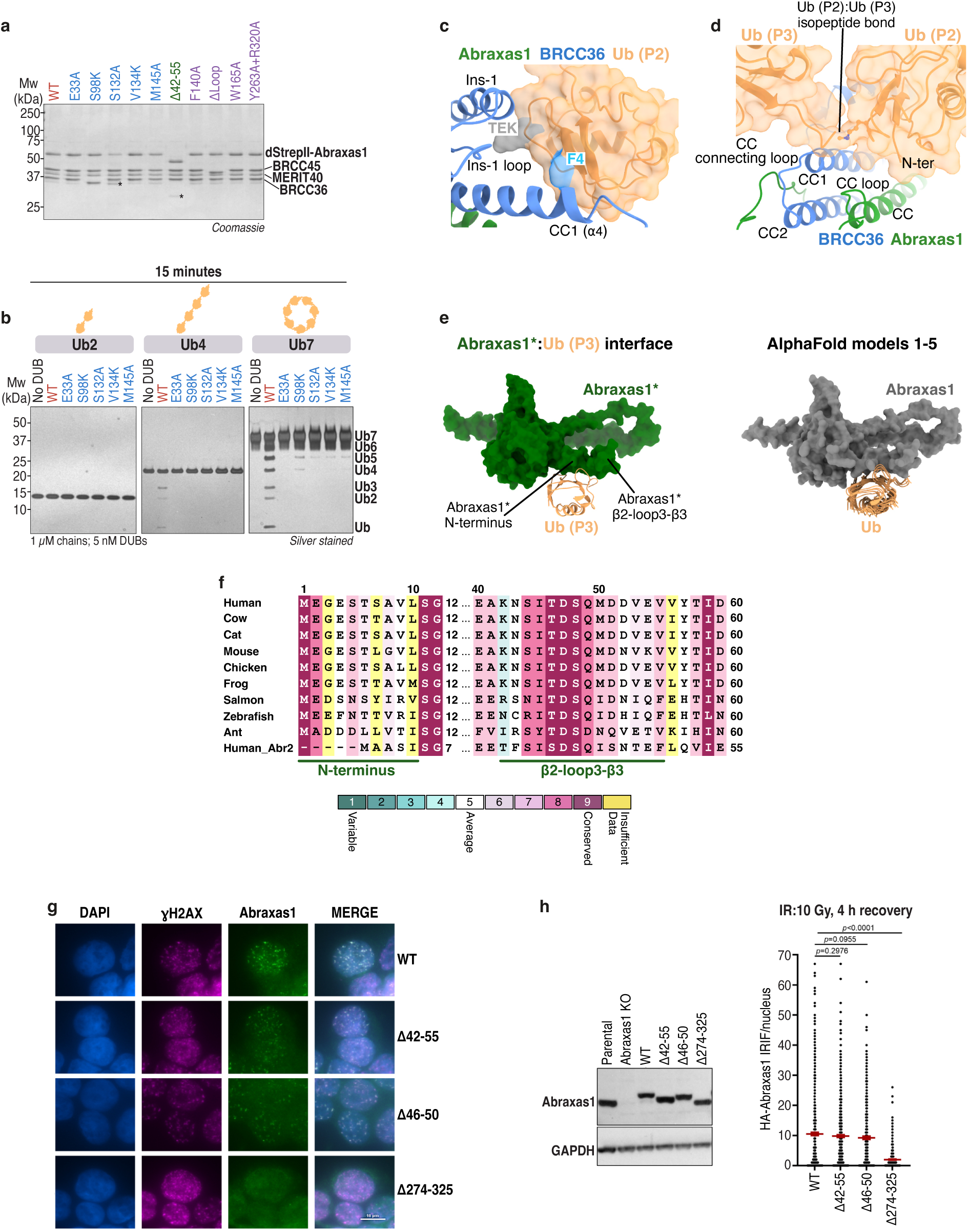
Structural, biochemical, and cell-based analyses and validation of the BRCC36 and Abraxas1 ubiquitin binding sites. **a**, SDS-PAGE analysis of ARISC WT (red) and variants bearing mutations or deletions in BRCC36 (blue), Abraxas1 (green), or BRCC45 (purple). BRCC45 ΔLoop refers to deletion of residues 161-170. dStrepII, double StrepII tag. * indicates Abraxas1 degradation product. **b,** K63-Ub2, -Ub4, and -Ub7 chains (1 µM) were incubated with ARISC variants (5 nM) for 15 minutes. Cleavage activity was analysed by SDS-PAGE and silver staining. Data are representative of two independent experiments. DUB, deubiquitylating enzyme. **c,** Close-up view of the BRCC36:Ub (P2) interface. Models are shown as cartoon, and colored as in Fig. 1a and Fig. 2a. The F4 patch and TEK box are indicated. Ins, insertion. **d,** Close-up view of the region near the Ub (P2):Ub (P3) isopeptide bond. Models are colored as in **c**. N-ter, N-terminus; CC, coiled coil. **e,** Structural comparisons between the Abraxas1*:Ub (P3) interface (from ARISC(E33A)–RAP80:K63-Ub7; green and orange) and the corresponding AlphaFold 3 models (grey and orange). Abraxas1 and ubiquitin are shown as surface and cartoon, and colored as in **c**. Ub, ubiquitin. **f,** Multiple sequence alignment of the Abraxas1 N-terminal and β2-loop3-β3 regions across nine species, with the corresponding residues in human Abraxas2 included. Sequence conservation was generated by the Consurf web server^1^, and indicated by a score between 1 and 9. The numbers at the top of the alignment refer to human Abraxas1. **g,** Representative images of Abraxas1 IRIF in HT-29 cells expressing WT or mutant proteins 4 h post irradiation (10 Gy). Scale bar is 10 µm. **h,** Western blots showing Abraxas1 protein levels in HT-29 cells reconstituted with WT or mutant proteins (l*eft*). Scatter plot showing quantification of the data described in **g**. Data represent mean ± SEM from n ≥ 350 nuclei examined over two independent experiments; p values are indicated, unpaired two-tailed t test (*right*). KO, knockout; WT, wild type.

**Supplementary Fig. 10:**
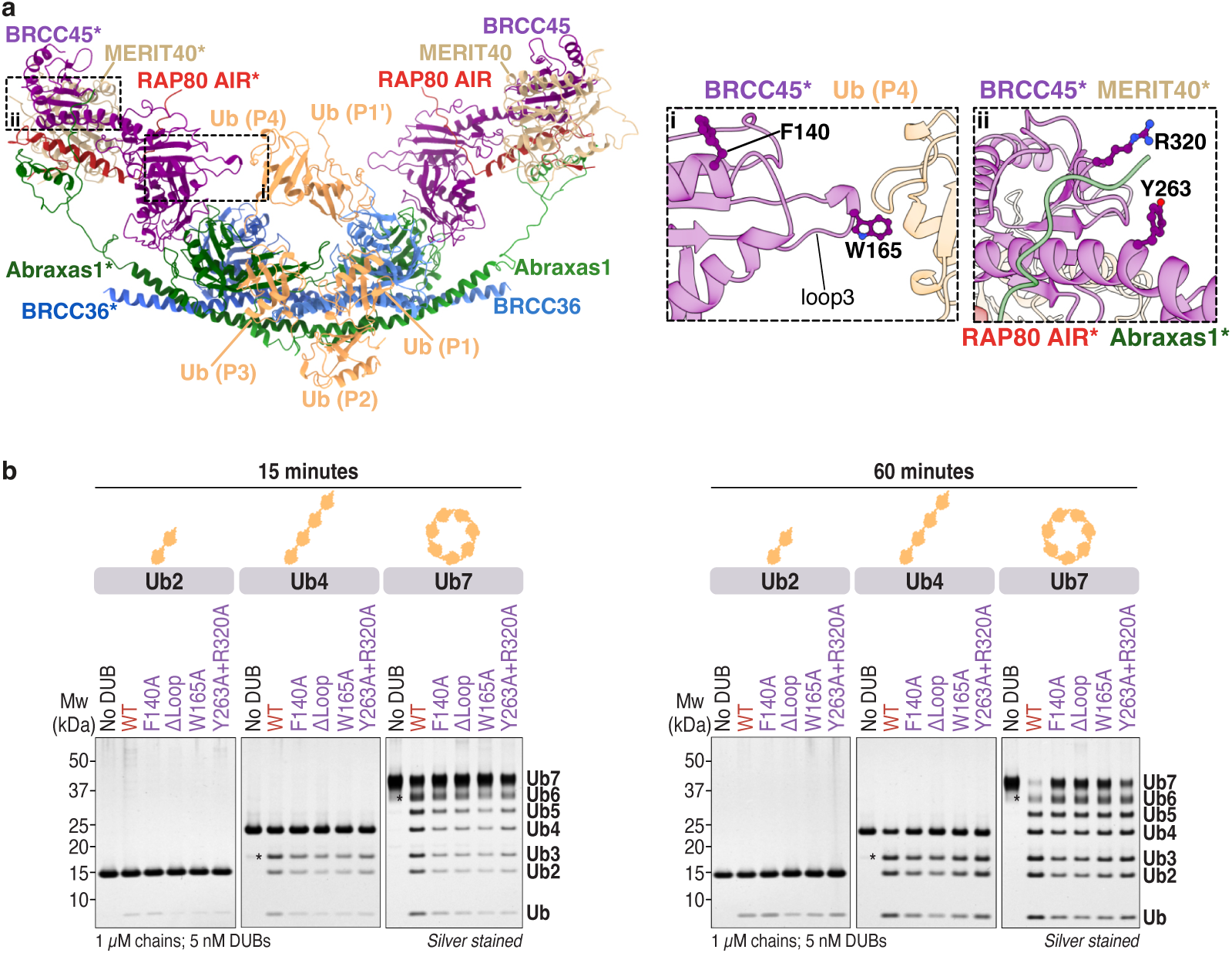
Determining the DUB activity of structure-guided BRCC45 mutants. **a**, The same model of the ARISC(E33A)–RAP80:K63-Ub7 complex shown in **Supplementary Fig. 5c** is depicted as cartoon representation, with ARISC–RAP80 subunits and ubiquitin chains colored as in Fig. 1a and Fig. 2a. Dashed black rectangles highlight regions analysed by mutagenesis (*left*). Close-up views and structural details of the predicted interacting residues at the putative BRCC45*:Ub (P4) (i) and BRCC45*:Abraxas1* (ii) interfaces. Residues are shown as ball & sticks (*right*). AIR, Abraxas1-interacting region. **b,** K63-Ub2, -Ub4, and -Ub7 chains (1 µM) were incubated with ARISC WT or the indicated ARISC variants (5 nM) for up to 60 minutes. Cleavage activity was analysed by SDS-PAGE and silver staining. Data are representative of two independent experiments. BRCC45 ΔLoop refers to deletion of amino acids 161-170. DUB, deubiquitylating enzyme; WT, wild type; Ub, ubiquitin.

**Supplementary Fig. 11:**
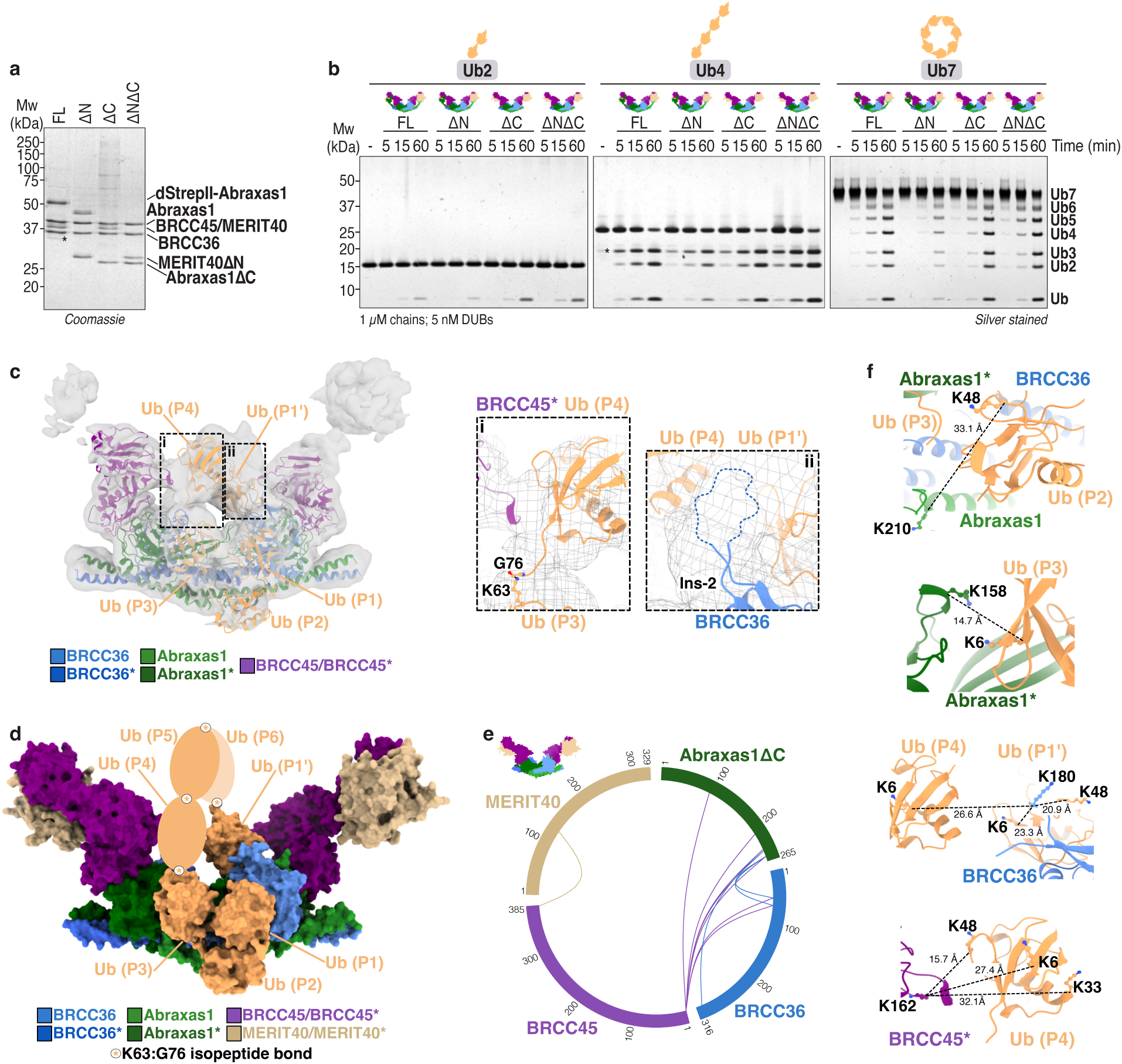
Biochemical analyses of ARISC variants, and validation of the ARISCΔC(E33A):K63-Ub7 and ARISCΔC(E33A):K63-Ub4 cryo-EM structures. **a**, SDS-PAGE analysis of FL ARISC, ARISCΔN, ARISCΔC, and ARISCΔNΔC. dStrepII, double StrepII tag. * indicates Abraxas1 degradation product. **b,** K63-Ub2, -Ub4, and -Ub7 chains (1 µM) were incubated with FL ARISC, ARISCΔN, ARISCΔC, or ARISCΔNΔC (5 nM) for the indicated time points. Cleavage activity was analysed by SDS-PAGE and silver staining. Data are representative of two independent experiments. FL, full-length; DUB, deubiquitylating enzyme. **c,** The ARISCΔN(E33A):K63-Ub7 heterogeneous refinement map (contour level = 0.118) is depicted in transparent surface. Cartoon models were rigid-body fitted into the cryo-EM density, and colored as in Fig. 1a and Fig. 2a. Dashed black rectangles highlight regions not well resolved in the final composite map (*left*). Close-up views and structural details of the putative BRCC45*:Ub (P4) (i) and BRCC36 Ins-2 (ii) interfaces. The cryo-EM heterogeneous refinement map is depicted as mesh; Ub (P3) K63^NZ^ and Ub (P4) G76^C^ are positioned at isopeptide bond distance (1.5 Å), and missing residues are depicted with a dashed line (*right*). Ins-2, insertion-2. **d,** The ARISCΔC(E33A):K63-Ub7 structure is shown as surface representation, with structural models colored as in **c**. Ubiquitin molecules not visualised in the cryo-EM map are depicted as orange ovals. Owing to low resolution in the extended arm regions, overlayed BRCC45 UEV-C and MERIT40 models are shown for size comparison only. **e,** Chemical cross-linking and mass spectrometry analyses of ARISCΔC. Proteins and cross-links are labelled and coloured as in Fig. 5f**-i**. Communal cross-links obtained from two independent experiments are shown. **f,** Close-up views and structural details of the Abraxas1:Ub (P2), Abraxas1*:Ub (P3), BRCC36:Ub (P1’):Ub (P4), and putative BRCC45*:Ub (P4) interfaces. Structural models are colored as in **c**. Residues involved in cross-links are shown as ball & sticks. Distances are indicated with dashed black lines and measured in Ångstrom (Å). Ub, ubiquitin.

**Supplementary Fig. 12:**
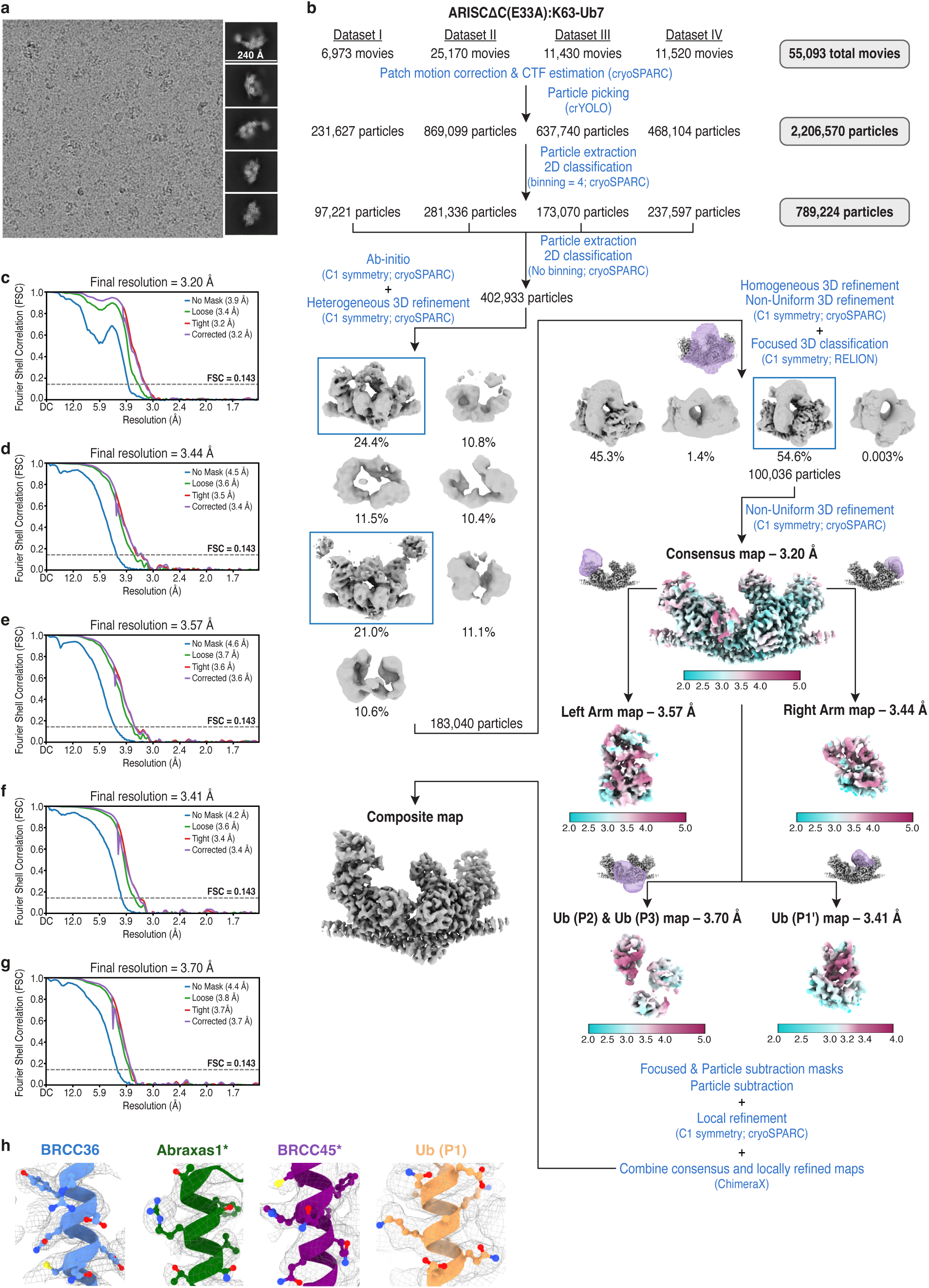
Cryo-EM analysis of the ARISCΔC(E33A):K63-Ub7 complex. **a**, Representative micrograph (out of 55,093 collected movies) and 2D class averages (out of 402,933 selected particles) for the ARISCΔC(E33A):K63-Ub7 dataset. **b,** Flowchart of data processing strategy. Boxed 3D classes were selected for subsequent processing. The final consensus and locally refined maps are shown and colored by local resolution. See **Supplementary Table 1** for details. **c-g,** Fourier shell correlation (FSC) curves for the final ARISCΔC(E33A):K63-Ub7 consensus map (**c**), and for the locally refined right (**d**) and left (**e**) arm, Ub (P1’) (**f**), and Ub (P2) & Ub (P3) (**g**) maps. Resolutions were calculated using the gold-standard FSC cutoff at 0.143 frequency. **h,** Representative density regions (at a contour level of 0.0514) from the ARISCΔC(E33A):K63-Ub7 composite map for the different components of the complex. Cartoon models are colored as in Fig. 1a and Fig. 2a, and residues are shown as ball & sticks. Ub, ubiquitin.

**Supplementary Fig. 13:**
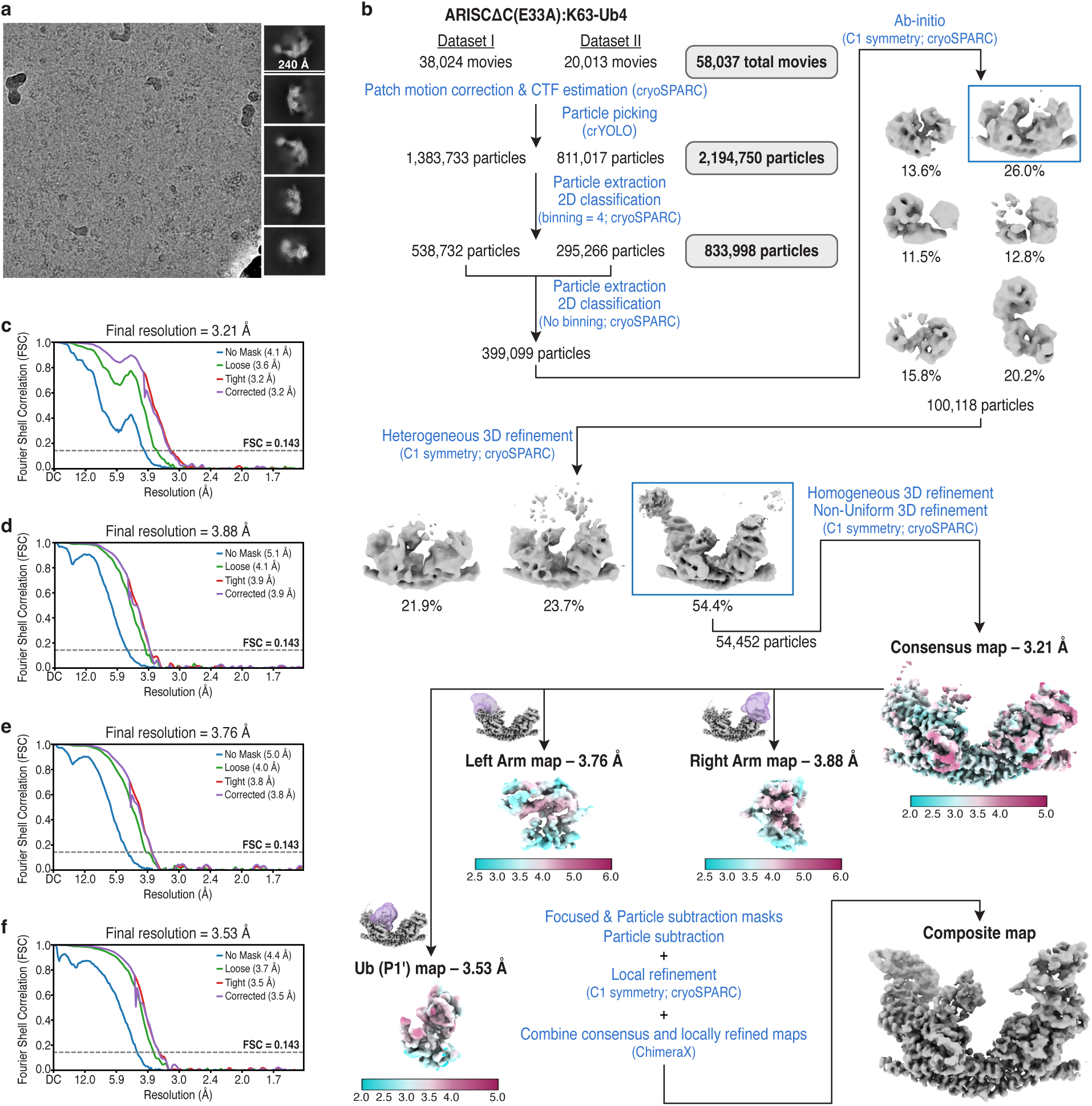
Cryo-EM analysis of the ARISCΔC(E33A):K63-Ub4 complex. **a**, Representative micrograph (out of 58,037 collected movies) and 2D class averages (out of 399,099 selected particles) for the ARISCΔC(E33A):K63-Ub4 dataset. **b,** Flowchart of data processing strategy. Boxed ab-initio and 3D classes were selected for subsequent processing. The final consensus and locally refined maps are shown and colored by local resolution. See **Supplementary Table 1** for details. **c-f,** Fourier shell correlation (FSC) curves for the final ARISCΔC(E33A):K63-Ub4 consensus map (**c**), and for the locally refined right (**d**) and left (**e**) arm, and Ub (P1’) (**f**) maps. Resolutions were calculated using the gold-standard FSC cutoff at 0.143 frequency.

**Supplementary Fig. 14:**
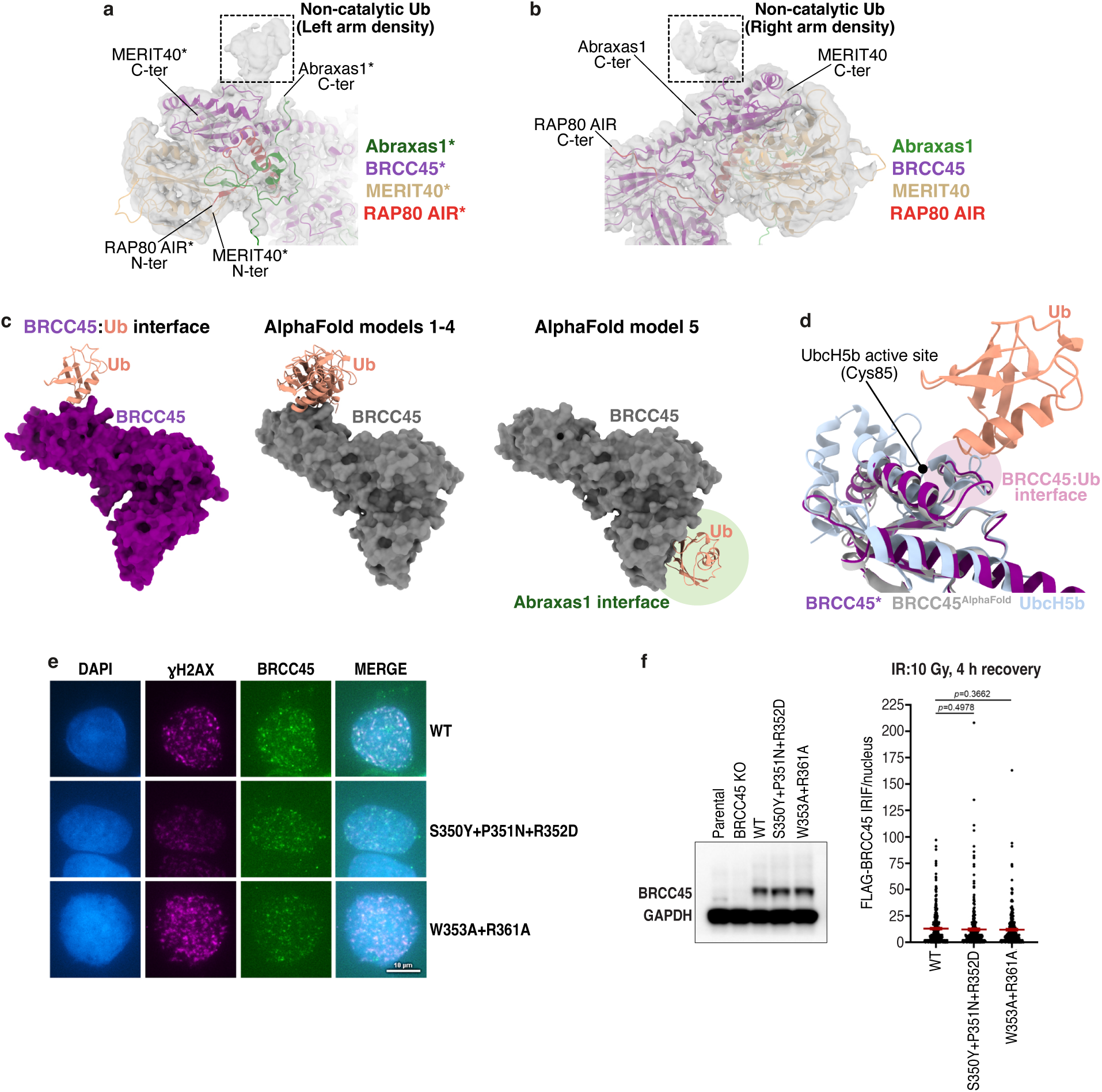
Structural analyses and validation of the ARISC–RAP80 non-catalytic ubiquitin binding sites. **a,b**, Close-up views of the left (**a**) and right (**b**) ARISC–RAP80 arms from the cryo-EM composite map of the ARISC(E33A)–RAP80:K63-Ub7 complex. The cryo-EM map at contour level of 0.0247 is depicted in transparent surface; cartoon models of ARISC–RAP80 subunits were rigid-body fitted into the cryo-EM density and colored as indicated. Dashed black rectangles highlight the non-catalytic ubiquitin sites located at top of each BRCC45* UEV-C domain (see Fig. 7b). AIR, Abraxas1-interacting region; N-ter, N-terminus; C-ter, C-terminus. **c,** Structural comparisons between the non-catalytic BRCC45*:Ub interface as seen in the ARISC(E33A)–RAP80:K63-Ub7 structure (purple and salmon) and the corresponding interfaces predicted in the AlphaFold 3 models (grey and salmon). BRCC45 and ubiquitin models are shown as surface and cartoon representations, and colored as in Fig. 1a and Fig. 7a. **d,** Structural overlays between the non-catalytic BRCC45*:Ub interface as seen in the ARISC(E33A)–RAP80:K63-Ub7 structure (purple and salmon) and the corresponding BRCC45 region predicted in AlphaFold 3 model 1 (light grey) as well as UbcH5b (PDBid = 3A33) (light blue). The UbcH5b active site cysteine residue (Cys85) and the BRCC45 or UbcH5b regions at the interface with ubiquitin are indicated. Structural models are shown as cartoon; the ubiquitin moiety loaded onto UbcH5b was omitted for clarity. Ub, ubiquitin. **e,** Representative immunofluorescence images of FLAG-BRCC45 IRIF in HEK293T cells expressing WT or indicated BRCC45 mutants. Cells were irradiated with 10 Gy and recovered for 4 h before fixation. Scale bar is 10 µm. **f,** Western blots showing expression of FLAG-BRCC45 in BRCC45 KO HEK293T cells transfected with WT or mutant BRCC45 expressing plasmids. Cells were harvested 48 h post transfection. GAPDH serves as loading control (*left*). Scatter plot showing quantification of the data described in **e**. Data represent mean ± SEM from n ≥ 336 nuclei examined over two independent experiments; p values are indicated, unpaired two-tailed t test (*right*). KO, knockout; WT, wild type.

**Supplementary Table 1:** Cryo-EM data collection, refinement and validation statistics.

|  | ARISC(E33A)-RAP80:K63-Ub7<br>EMD-55122<br>PDBid: 9SQY | ARISCΔC(E33A):K63-Ub7<br>EMD-55119<br>PDBid: 9SQW | ARISCΔC(E33A):K63-Ub4<br>EMD-55118<br>PDBid: 9SQV |
| --- | --- | --- | --- |
| **Data Collection** |  |  |  |
| Microscope |  | FEI Titan KRIOS |  |
| Detector |  | Falcon 4i |  |
| Voltage (keV) |  | 300 |  |
| Mode |  | Counting |  |
| Energy Filter (eV) |  | 10 |  |
| Pixel size (Å) |  | 0.74 |  |
| Magnification (x) |  | 165,000 |  |
| Electron dose (e<sup>-</sup>/Å<sup>2</sup>) | 45.5 | 45.4-45.5 | 45.6-45.9 |
| Exposure (s) | 3.4 | 3.36-3.75 | 3.4 |
| Electron dose per frame (e<sup>-</sup>/Å<sup>2</sup>) | 1.17 | 1.06-1.17 | 1.17-1.18 |
| Fractions | 39 | 39-43 | 39 |
| Total no. of movies | 10,649 | 55,093 | 58,037 |
| Defocus range (μm) |  | -0.9 to -3.0 |  |
| AFIS range (μm) |  | 10 |  |
| **Data Processing** |  |  |  |
| Symmetry Point Group |  | C1 |  |
| Initial particle number | 789,840 | 2,206,570 | 2,194,750 |
| Final particle number | 214,650 | 100,036 | 54,452 |
| Map resolution (Å) |  |  |  |
| Consensus | 2.92 | 3.20 | 3.21 |
| Right/Left Arms | 4.09/4.02 | 3.44/3.57 | 3.88/3.76 |
| Ub (P1') | 3.16 | 3.41 | 3.53 |
| Ub (P2) & Ub (P3) | 3.17 | 3.70 | - |
| Non-catalytic Ub Right/Left Arms | 3.99/3.91 | - | - |
| FSC threshold |  | 0.143 |  |
| Map resolution range (Å) |  |  |  |
| Consensus | 1.6-6.0 | 2.0-5.0 | 2.0-5.0 |
| Right/Left Arms | 2.0-6.0/2.0-6.0 | 2.0-5.0/2.0-5.0 | 2.5-6.0/2.5-6.0 |
| Ub (P1') | 2.0-4.0 | 2.0-4.0 | 2.0-5.0 |
| Ub (P2) & Ub (P3) | 2.5-5.0 | 2.5-5.0 | - |
| Non-catalytic Ub Right/Left Arms | 2.0-6.0/2.0-6.0 | - | - |
| **Refinement** |  |  |  |
| Model resolution (Å) | 1.8 | 1.9 | 1.9 |
| FSC threshold |  | 0.143 |  |
| Model composition |  |  |  |
| Chains | 17 | 11 | 9 |
| Non-hydrogen atoms | 23,659 | 13,761 | 12,545 |
| Protein residues | 2,956 | 1,722 | 1,572 |
| Ligands | Zn: 2 | Zn: 2 | Zn: 2 |
| B factors (Å<sup>2</sup>) |  |  |  |
| Protein (min/max/mean) | 19.92/264.02/94.61 | 32.53/201.57/91.13 | 23.73/174.89/85.20 |
| Ligands (min/max/mean) | 84.40/91.04/87.72 | 58.95/105.71/82.33 | 62.47/131.97/97.22 |
| Map: model CC |  |  |  |
| CC (mask/box/peaks/volume) | 0.72/0.77/0.73/0.75 | 0.75/0.75/0.74/0.77 | 0.72/0.75/0.72/0.74 |
| Mean CC for ligands | 0.75 | 0.80 | 0.76 |
| R.m.s. deviations |  |  |  |
| Bond lengths (Å) | 0.003 | 0.003 | 0.002 |
| Bond angles (°) | 0.641 | 0.484 | 0.507 |
| **Validation** |  |  |  |
| MolProbity score | 1.79 | 1.92 | 2.05 |
| Clash score | 9.06 | 12.9 | 18.01 |
| Rotamers outliers (%) | 0.53 | 0.19 | 0.42 |
| CB outliers (%) | 0 | 0 | 0 |
| CaBLAM outliers (%) | 2.60 | 2.40 | 2.50 |
| Ramachandran plot |  |  |  |
| Favored (%) | 95.64 | 95.73 | 95.71 |
| Allowed (%) | 4.26 | 4.27 | 4.29 |
| Outliers (%) | 0.10 | 0 | 0 |

**Supplementary Table 2:** List of expression constructs.

| ARISC & BRISC variants |  |  |  |  |  |  |  |  |  |
| --- | --- | --- | --- | --- | --- | --- | --- | --- | --- |
| Expression Construct | Species | Cloning/Gene Synthesis | Protein/Sub-complex 1 (name) | Protein/Sub-complex 1 (aa) | Tag | Protein/Sub-complex 2 (name) | Protein/Sub-complex 2 (aa) | Tag | Expression System |
| pFL-pUCDM:FL ARISC WT | Human | codon-optimised (GenScript) | BRCC36-Abraxas1 | 1-316 (BRCC36); 1-409 (Abraxas1) | N-dStrepII (Abraxas1) | BRCC45-MERIT40 | 1-383 (BRCC45); 1-329 (MERIT40) | N-6xHis (BRCC45) | *Sf9* |
| pFL-pUCDM:FL BRISC WT | Human | codon-optimised (GenScript) | BRCC36-Abraxas2 | 1-316 (BRCC36); 1-415 (Abraxas2) | N-dStrepII (Abraxas2) | BRCC45-MERIT40 | 1-383 (BRCC45); 1-329 (MERIT40) | N-6xHis (BRCC45) | *Sf9* |
| pFL-pUCDM:ARISCΔN WT | Human | cloning (see **Supplementary Table 3**) | BRCC36-Abraxas1 | 1-316 (BRCC36); 1-409 (Abraxas1) | none | BRCC45-MERIT40ΔN | 1-383 (BRCC45); 72-329 (MERIT40ΔN) | N-6xHis (BRCC45) | *Sf9* |
| pFL-pUCDM:ARISCΔC WT | Human | cloning (see **Supplementary Table 3**) | BRCC36-Abraxas1ΔC | 1-316 (BRCC36); 1-265 (Abraxas1ΔC) | none | BRCC45-MERIT40 | 1-383 (BRCC45); 1-329 (MERIT40) | N-6xHis (BRCC45) | *Sf9* |
| pFL-pUCDM:ARISCΔNΔC WT | Human | cloning (see **Supplementary Table 3**) | BRCC36-Abraxas1ΔC | 1-316 (BRCC36); 1-265 (Abraxas1ΔC) | none | BRCC45-MERIT40ΔN | 1-383 (BRCC45); 72-329 (MERIT40ΔN) | N-6xHis (BRCC45) | *Sf9* |
| pFL-pUCDM:FL ARISC E33A | Human | codon-optimised (GenScript) | BRCC36 E33A-Abraxas1 | 1-316 (BRCC36 E33A); 1-409 (Abraxas1) | N-dStrepII (Abraxas1) | BRCC45-MERIT40 | 1-383 (BRCC45); 1-329 (MERIT40) | N-6xHis (BRCC45) | *Sf9* |
| pFL-pUCDM:FL ARISC E33A<sup>BRCC36(S98K)</sup> | Human | codon-optimised (GenScript) | BRCC36 E33A+S98K-Abraxas1 | 1-316 (BRCC36 E33A+S98K); 1-409 (Abraxas1) | N-dStrepII (Abraxas1) | BRCC45-MERIT40 | 1-383 (BRCC45); 1-329 (MERIT40) | N-6xHis (BRCC45) | *Sf9* |
| pFL-pUCDM:ARISC E33A<sup>Abraxas1(Δ42-55)</sup> | Human | codon-optimised (GenScript) | BRCC36 E33A-Abraxas1 Δ42-55 | 1-316 (BRCC36 E33A); 1-409 (Abraxas1 Δ42-55) | N-dStrepII (Abraxas1) | BRCC45-MERIT40 | 1-383 (BRCC45); 1-329 (MERIT40) | N-6xHis (BRCC45) | *Sf9* |
| pFL-pUCDM:ARISC E33A<sup>BRCC45(ΔLoop)</sup> | Human | codon-optimised (GenScript) | BRCC36 E33A-Abraxas1 | 1-316 (BRCC36 E33A); 1-409 (Abraxas1) | N-dStrepII (Abraxas1) | BRCC45 ΔLoop-MERIT40 | 1-383/Δ161-170 (BRCC45 ΔLoop); 1-329 (MERIT40) | N-6xHis (BRCC45) | *Sf9* |
| pFL-pUCDM:ARISCΔC E33A | Human | codon-optimised (GenScript) | BRCC36 E33A-Abraxas1ΔC | 1-316 (BRCC36 E33A); 1-265 (Abraxas1ΔC) | N-dStrepII (Abraxas1) | BRCC45-MERIT40 | 1-383 (BRCC45); 1-329 (MERIT40) | N-6xHis (BRCC45) | *Sf9* |
| pFL-pUCDM:FL ARISC S98K | Human | codon-optimised (GenScript) | BRCC36 S98K-Abraxas1 | 1-316 (BRCC36 S98K); 1-409 (Abraxas1) | N-dStrepII (Abraxas1) | BRCC45-MERIT40 | 1-383 (BRCC45); 1-329 (MERIT40) | N-6xHis (BRCC45) | *Sf9* |
| pFL-pUCDM:FL ARISC S132A | Human | codon-optimised (GenScript) | BRCC36 S132A-Abraxas1 | 1-316 (BRCC36 S132A); 1-409 (Abraxas1) | N-dStrepII (Abraxas1) | BRCC45-MERIT40 | 1-383 (BRCC45); 1-329 (MERIT40) | N-6xHis (BRCC45) | *Sf9* |
| pFL-pUCDM:FL ARISC V134K | Human | codon-optimised (GenScript) | BRCC36 V134K-Abraxas1 | 1-316 (BRCC36 V134K); 1-409 (Abraxas1) | N-dStrepII (Abraxas1) | BRCC45-MERIT40 | 1-383 (BRCC45); 1-329 (MERIT40) | N-6xHis (BRCC45) | *Sf9* |
| pFL-pUCDM:FL ARISC M145A | Human | codon-optimised (GenScript) | BRCC36 M145A-Abraxas1 | 1-316 (BRCC36 M145A); 1-409 (Abraxas1) | N-dStrepII (Abraxas1) | BRCC45-MERIT40 | 1-383 (BRCC45); 1-329 (MERIT40) | N-6xHis (BRCC45) | *Sf9* |
| pFL-pUCDM:ARISC Δ42-55 | Human | codon-optimised (GenScript) | BRCC36-Abraxas1 Δ42-55 | 1-316 (BRCC36); 1-409/Δ42-55 (Abraxas1 Δ42-55) | N-dStrepII (Abraxas1) | BRCC45-MERIT40 | 1-383 (BRCC45); 1-329 (MERIT40) | N-6xHis (BRCC45) | *Sf9* |
| pFL-pUCDM:FL ARISC F140A | Human | codon-optimised (GenScript) | BRCC36-Abraxas1 | 1-316 (BRCC36); 1-409 (Abraxas1) | N-dStrepII (Abraxas1) | BRCC45 F140A-MERIT40 | 1-383 (BRCC45 F140A); 1-329 (MERIT40) | N-6xHis (BRCC45) | *Sf9* |
| pFL-pUCDM:ARISC ΔLoop | Human | codon-optimised (GenScript) | BRCC36-Abraxas1 | 1-316 (BRCC36); 1-409 (Abraxas1) | N-dStrepII (Abraxas1) | BRCC45 ΔLoop-MERIT40 | 1-383/Δ161-170 (BRCC45 ΔLoop); 1-329 (MERIT40) | N-6xHis (BRCC45) | *Sf9* |
| pFL-pUCDM:FL ARISC W165A | Human | codon-optimised (GenScript) | BRCC36-Abraxas1 | 1-316 (BRCC36); 1-409 (Abraxas1) | N-dStrepII (Abraxas1) | BRCC45 W165A-MERIT40 | 1-383 (BRCC45 W165A); 1-329 (MERIT40) | N-6xHis (BRCC45) | *Sf9* |
| pFL-pUCDM:FL ARISC Y263A+R320A | Human | codon-optimised (GenScript) | BRCC36-Abraxas1 | 1-316 (BRCC36); 1-409 (Abraxas1) | N-dStrepII (Abraxas1) | BRCC45 Y263A+R320A-MERIT40 | 1-383 (BRCC45 Y263A+R320A); 1-329 (MERIT40) | N-6xHis (BRCC45) | *Sf9* |
| RAP80 variants |  |  |  |  |  |  |  |  |  |
| Expression Construct | Species | Cloning/Gene Synthesis | Protein/Sub-complex 1 (name) | Protein/Sub-complex 1 (aa) | Tag | Protein/Sub-complex 2 (name) | Protein/Sub-complex 2 (aa) | Tag | Expression System |
| pFastBac-HTB:FL RAP80 WT | Human | cloning (see **Supplementary Table 3**) | RAP80 | 1-719 | N-6xHis | none | none | none | *Sf9* |
| pFastBac-HTB:RAP80 AIR | Human | cloning (see **Supplementary Table 3**) | RAP80 AIR | 273-402 | N-6xHis | none | none | none | *Sf9* |
| pFastBac-HTB:RAP80 ΔUIMs | Human | cloning (see **Supplementary Table 3**) | RAP80 ΔUIMs | 1-719/Δ74-131 | N-6xHis | none | none | none | *Sf9* |
| pFastBac-HTB:RAP80 ΔZnF | Human | cloning (see **Supplementary Table 3**) | RAP80 ΔZnF | 1-719/Δ499-588 | N-6xHis | none | none | none | *Sf9* |
| Ubiquitin variants |  |  |  |  |  |  |  |  |  |
| Expression Construct | Species | Cloning/Gene Synthesis | Protein/Sub-complex 1 (name) | Protein/Sub-complex 1 (aa) | Tag | Protein/Sub-complex 2 (name) | Protein/Sub-complex 2 (aa) | Tag | Expression System |
| pProEx-HTB:Cys0-ubiquitin K63R | Human | cloning (see **Supplementary Table 3**) | Cys0-ubiquitin K63R | 1-76 | N-6xHis | none | none | none | *E. coli* |
| pET3a:ubiquitin R74C ΔGG | Human | codon-optimised (GenScript) | ubiquitin R74C ΔGG | 1-74 | none | none | none | none | *E. coli* |
| pET3a:ubiquitin K63R | Human | codon-optimised (GenScript) | ubiquitin K63R | 1-76 | N-8xHis | none | none | none | *E. coli* |

**Supplementary Table 3:**
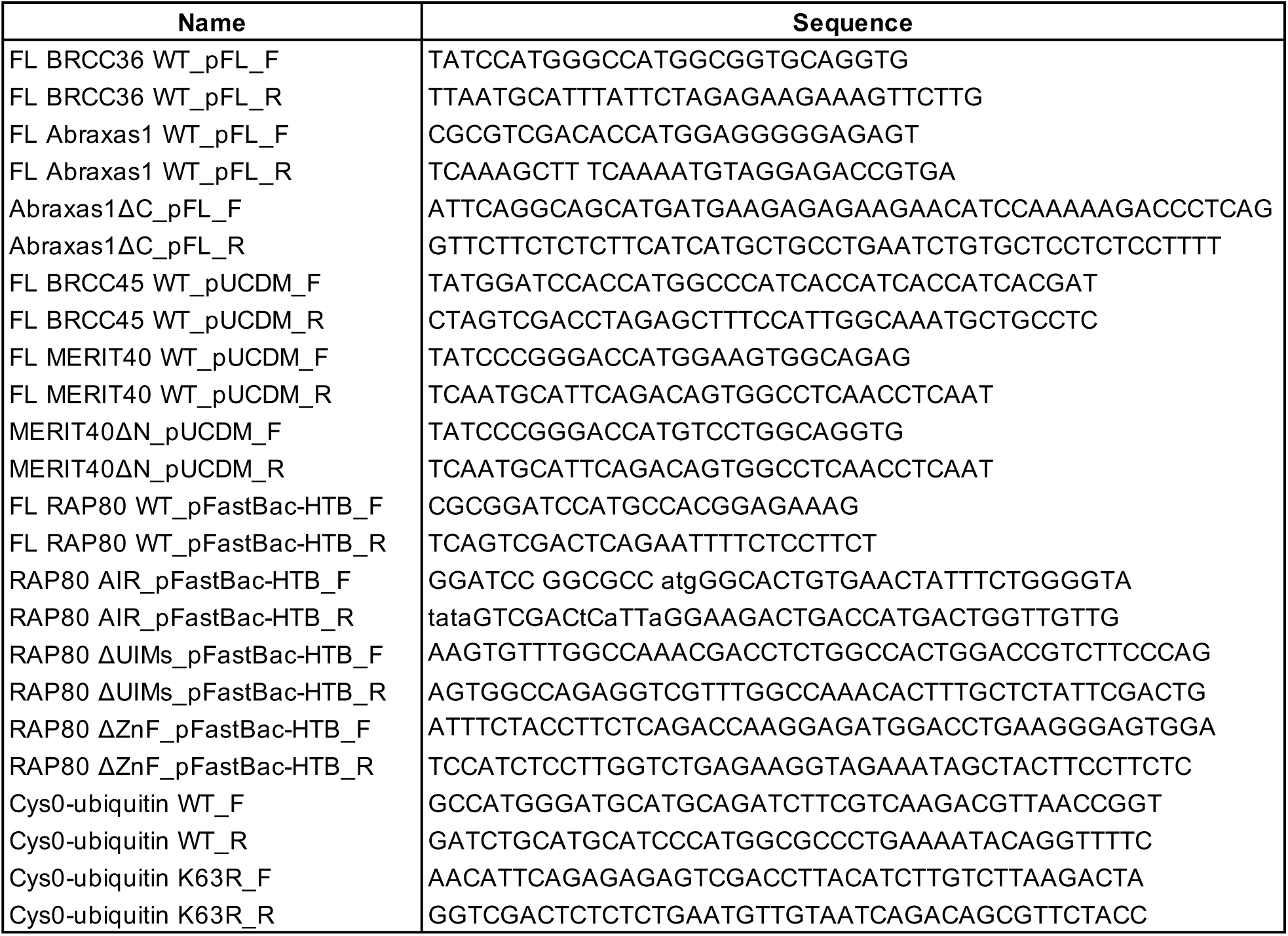
List of primers.

